# Domino-like Effect of C112R Mutation on ApoE4 Aggregation and Its Reduction by Alzheimer’s Disease Drug Candidate

**DOI:** 10.1101/2022.10.09.511473

**Authors:** Michal Nemergut, Sergio M. Marques, Lukas Uhrik, Tereza Vanova, Marketa Nezvedova, Darshak Chandulal Gadara, Durga Jha, Jan Tulis, Veronika Novakova, Joan Planas-Iglesias, Antonin Kunka, Anthony Legrand, Hana Hribkova, Veronika Pospisilova, Jiri Sedmik, Jan Raska, Zbynek Prokop, Jiri Damborsky, Dasa Bohaciakova, Zdenek Spacil, Lenka Hernychova, David Bednar, Martin Marek

## Abstract

**Background:** Apolipoprotein E (ApoE) ε4 genotype is the most prevalent risk factor for late-onset Alzheimer’s Disease (AD). Although ApoE4 differs from its non-pathological ApoE3 isoform only by the C112R mutation, the molecular mechanism of its proteinopathy is unknown.

**Methods:** Here, we reveal the molecular mechanism of ApoE4 aggregation using a combination of experimental and computational techniques, including X-ray crystallography, site-directed mutagenesis, hydrogen-deuterium mass spectrometry (HDX-MS), static light scattering and molecular dynamics simulations. Treatment of ApoE ε3/ε3 and ε4/ε4 cerebral organoids with tramiprosate was used to compare the effect of tramiprosate on ApoE4 aggregation at the cellular level.

**Results:** We found that C112R substitution in ApoE4 induces long-distance (>15 Å) conformational changes leading to the formation of a V-shaped dimeric unit that is geometrically different and more aggregation-prone than the ApoE3 structure. AD drug candidate tramiprosate and its metabolite 3-sulfopropanoic acid induce ApoE3-like conformational behavior in ApoE4 and reduce its aggregation propensity. Analysis of ApoE ε4/ε4 cerebral organoids treated with tramiprosate revealed its effect on cholesteryl esters, the storage products of excess cholesterol.

**Conclusions:** Our results connect the ApoE4 structure with its aggregation propensity, providing a new druggable target for neurodegeneration and ageing.

## Background

Seeing loved ones losing their ability to recall their memories is devastating. However, this is the reality of life for the families of an increasing number of individuals with Alzheimer’s disease (AD) [1]. Currently, more than 55 million people live with dementia worldwide, and nearly 10 million new cases are diagnosed every year [2,3]. Despite the considerable funding for AD drug development [4], no disease-modifying therapies currently exist, and numerous clinical trials have failed to demonstrate any benefits [5,6]. There is an urgent need to improve our understanding of the molecular basis of this devastating disease and provide validated therapeutic targets.

Most of the drug candidates in ongoing clinical trials target two pathological features of AD: (i) neurofibrillary tangles comprised of hyperphosphorylated tau protein and (ii) amyloid plaques comprised of Aβ peptides [7,8]. One such candidate for the treatment of AD is ALZ-801, a prodrug of tramiprosate [9]. Orally administered tramiprosate was reported to inhibit Aβ -amyloid aggregation by modulation of its conformational flexibility [10,11]. ALZ- 801 and tramiprosate are metabolized into the 3-sulfopropanoic acid (SPA), an endogenous molecule of the human brain with similar biological activity to tramiprosate [12]. Despite the great potential of tramiprosate to modify the progression of AD, a positive clinical effect was only observed in homozygotes for apolipoprotein E4 (ApoE4 ε4/ε4), a well-established and long-standing genetic risk factor for AD [13–18].

Human ApoE is a 299-residue protein component of lipoprotein particles [19] and a key regulator of Aβ aggregation and clearance [20,21]. The three polymorphic isoforms of ApoE, i.e., ApoE2, ApoE3, and ApoE4, differ in amino acid composition at positions 112 and 158 [22]. Compared to the least frequent isoform ApoE2 (C112/C158), the most common isoform ApoE3 differs by a single mutation (C112/R158), whereas the AD-associated ApoE4 contains two mutations (R112/R158) [23]. Recently, it has been proposed that ApoE4 could be a new druggable target for AD treatment [24,25]. Various therapeutic approaches have been proposed to reduce ApoE4 toxicity or modify its physiological activity, including immunotherapy, antisense oligonucleotides, and small molecule correctors [26–29].

In its lipid-free state, ApoE exists as a mixture of monomers, dimers, tetramers and octamers, with tetramers being the prevalent species [30]. The colloidal heterogeneity of full-length ApoE and its high propensity for fragmentation has made all crystallization attempts to date unsuccessful. The lipid-free structure of ApoE3 in solution was resolved using NMR spectroscopy only when five “monomerizing” mutations (ApoE3_M_; F257A/W264R/V269A/L279Q/V287E) were introduced [31]. The ApoE protein comprises of the N-terminal domain formed by a four-helix bundle (NTD, residues 1-167, helices N1, N2, and H1-H4), followed by the hinge domain (HD, residues 168-205, helices HD1-HD2), and the C-terminal domain (CTD, residues 206-299, helices C1-C3) [32]. Several biophysical studies reported that lipid-free ApoE4 is more aggregation-prone than lipid-free ApoE3, suggesting that aggregation may play an essential role in driving the pathological effects of ApoE4 [33,34]. Moreover, ApoE4 aggregates have been demonstrated to be toxic to neuronal cells and have been found in amyloid plaques of AD patients [35–37]. These aggregates adopt a protofilament-like morphology mediated by the NTD [38]. However, the underlying mechanism of how C112R substitution increases the pathological effect of ApoE4 in AD remains poorly understood.

Here, we utilized a comprehensive structural analysis of newly obtained ApoE structures to reveal the molecular features and protein-protein contacts distinguishing ApoE4 from its non-pathological isoforms. Our work delineates the long-distance structural effects induced by a single mutation (C112R) directly impacting the ApoE aggregation tendency. We propose a mechanism for NTD-mediated ApoE aggregation that (i) effectively explains the high aggregation propensity of ApoE4; (ii) explains an unequal efficacy of tramiprosate in AD patients with different ApoE genotypes; and (iii) provides a validated therapeutic target for AD drug development. The proposed mechanism is further supported by biophysical, biochemical, mass spectrometric, computational and cell biology experiments.

## Methods

(MOLECULAR BIOLOGY METHODS)

### Mutagenesis

Megaprimer PCR-based mutagenesis [39] was applied to create single-point (W34A) and four-point (R38A/E45A/E49A/R145A) mutations in both ApoE3/4_WT_ and ApoE3/4_M_ variants purchased from GeneArt (Thermo Fisher Scientific). A megaprimer with the desired mutations was synthesized in the first PCR reaction using primers listed in **Supplementary Table 1**. The megaprimer was purified by gel electrophoresis and used as a primer in the second round of PCR to generate the complete DNA sequence. After temperature cycling, the parental template was digested by *Dpn*I (2 h at 37 °C), and the mutated plasmid was transformed into homemade competent Dh5α cells. The truncated ApoE variants (deletion of residues 1-23) were amplified by PCR, cloned into *Nde*I/*BamH*I sites of the pET-21b expression vector, and transformed into homemade competent Dh5α cells. All plasmids were isolated from three randomly selected colonies and sent for DNA sequencing (Eurofins Genomics, Germany) to verify the presence of the mutation.

### Overexpression and purification

*Escherichia coli* BL21(DE3) cells (NEB, USA) were transformed with pET-21b plasmid encoding His-thioredoxin-ApoE, plated on agar plates with ampicillin (100 μg/ml) and grown overnight at 37 °C. A single colony was used for inoculation of 50 ml of LB medium containing 100 µg/ml ampicillin followed by overnight incubation at 37 °C. The next day, 1 l of LB Broth medium containing 100 µg/ml ampicillin was inoculated to OD_600_ ∼ 0.1 with an overnight pre-culture harbouring a corresponding expression plasmid. Protein expression at 37 °C was induced by the addition of IPTG to a final concentration of 1 mM upon reaching OD_600_ ∼ 0.6 and continued for the next 3 hours. After cultivation, the cells were harvested by centrifugation (15 min, 4000 g, 4 °C) and resuspended in TBS buffer A (10 mM Tris-HCl, 50 mM NaCl, 20 mM imidazole, pH 7.5) containing DNase (20 μg/ml). The cells were then disrupted by sonification using Sonic Dismembrator Model 705 (Fisher Scientific, USA). The lysate was clarified by centrifugation (50 min, 21000 g, 4 °C) using a Sigma 6-16K centrifuge (SciQuip, UK). The filtrated supernatant containing His-thioredoxin-ApoE was applied to a 5 ml Ni-NTA Superflow Cartridge (Qiagen, Germany) pre-equilibrated with TBS buffer A. His-thioredoxin-ApoE was eluted with TBS buffer B (10 mM Tris-HCl, 50 mM NaCl pH, 250 mM imidazole, pH 7.5). Eluted His-thioredoxin-ApoE was treated with homemade 3C protease (overnight at 4 °C) to remove the His-thioredoxin tag. Finally, ApoE was purified by size exclusion chromatography using Äkta pure system (Cytivia, USA) equipped with Superdex 200 Increase 10/300 GL column equilibrated with TBS buffer A. The protein purity was verified by SDS-PAGE. The same protocol was used for all ApoE variants listed in **Table 1**.

(BIOPHYSICAL AND STRUCTURAL STUDIES)

### Circular dichroism

Ellipticity of ApoE variants was recorded at 4 °C using a Chirascan spectropolarimeter (Applied Photophysics, UK). Spectra were collected in the range of 195 to 260 nm with a 0.25 s response time and 1 nm bandwidth in 1 cm quartz cuvettes. The final spectrum represents an average of three individual scans, corrected for absorbance of the buffer.

### Dynamic light scattering

The size of ApoE variants was determined using the Uncle instrument (USA). Capillaries were filled with the protein samples with a concentration of ∼ 0.2 mg/ml and the size distribution profiles were collected at room temperature. The final mass distribution functions represent the average values of twenty individual repeats.

### Static light scattering

The aggregation of ApoE variants was studied by static light scattering (SLS) using the Uncle instrument (USA). ApoE proteins at a concentration of ∼10µM were mixed with 3- sulphopropanoic acid (SPA) in 2000 molar excess (20 mM). Capillaries were filled with the ApoE-SPA complexes and incubated at 37 °C. The aggregation process was monitored as SLS at 266 nm for 15 hours. We applied a lowpass filter with a passband frequency of 0.03183 to suppress high frequency fluctuations in the signal by using the *lowpass (x,wpass,′ImpulseResponse′,′iir′)* function in MATLAB R2019b. Final aggregation curves represent an average of three individual curves.

### Differential scanning fluorimetry

Thermal denaturation of ApoE proteins was studied by using a NanoDSF Prometheus instrument (NanoTemper, Germany). The protein samples at a concentration of 0.2 mg/ml were heated at a scan rate of 1 °C/min from 25 to 95 °C. The intrinsic tryptophan fluorescence intensity ratio at 350 nm 330 nm (F350/F330) and scattering were used to monitor the unfolding process and aggregation, respectively.

### Transmission electron microscopy

ApoE samples at a concentration of ∼ 0.4 mg/ml were applied to carbon-coated grids for 2 min. The unabsorbed liquid material was removed using filter papers and the grids were subsequently negatively stained with a 2 % solution of ammonium molybdate for 1 min. After staining, the excess staining solution was removed using filter papers. ApoE samples were imaged in a Morgagni 268D transmission electron microscope (FEI Company) equipped with a Veleta digital camera (Olympus).

### Crystallization

To determine the structure of ApoE4, we crystallized ApoE4_M_ in fusion with maltose-binding protein (His-MBP-ApoE4_M_). Purification of His-MBP-ApoE4_M_ was the same as for the other variants mentioned above except for the cleavage step. Whole purified His-MBP-ApoE4_M_ was concentrated to a final concentration of ∼ 8 mg/ml using Centrifugal Filter Units Amicon^R^ Ultra-15 Ultracel^R^-3K (Merck Millipore Ltd., Ireland). The crystallization was performed in Easy-Xtal 15-well crystallization plates by a hanging drop vapour diffusion, where 1 μl of His-MBP-ApoE4_M_ was mixed with the reservoir solution (8AX9: 100 mM HEPES pH 7.5, 11% PEG 3350; 8AX8: 100 mM NaCl, 20 mM KMES pH 6.7, 6.6% PEG 4000; 8CDY: 100 mM NaHEPES pH 7.0, 200 mM NaCl, 8% PEG 8000; 8CE0: 100 mM HEPES pH 7.0, 10% PEG 6000) in the ratio 1:1 and equilibrated against 500 μl of the reservoir solution. The best-looking crystals were flash-frozen in liquid nitrogen and stored to be taken for X-ray diffraction analysis.

### Diffraction data collection and processing

Diffraction data were collected at the Swiss Light Source synchrotron at the wavelength of 1.0 Å. The data were processed using XDS [40], and Aimless [41] was used for data merging. Initial phases of ApoE were solved by molecular replacement using Phaser [42] implemented in Phenix^5^. The structure of the N-terminal domain (NTD) ApoE (PDB ID: 5GS9) was employed as a search model for molecular replacement. The refinement was carried out in several cycles of automated refinement in phenix.refine program [43] and manual model building performed in Coot [44]. The output PDB deprived of hydrogens in PyMOL (Schrödinger, LLC) was used in Coot to fit into the experimental ligand electron density. The final models were validated using tools provided in Coot [44]. Structural data were graphically visualized with PyMOL Molecular Graphics System (Schrödinger, LLC). Atomic coordinates and structure factors for both ApoE structures were deposited in the Protein Data Bank under the codes 8AX8 and 8AX9, respectively.

### Hydrogen Deuterium Exchange (HDX) mass spectrometry (MS)

In order to prepare peptide mapping samples and undeuterated controls, both ApoE3/4_WT_ and ApoE3/4_M_ were diluted with H_2_O buffer (10 mM Tris-HCl, 50 mM NaCl, pH 7.5) to a final concentration of 2 µM. All deuterated ApoE samples either free or with SPA in the molar ratio 1: 1000 (ApoE: SPA) were preincubated for 20 min at laboratory temperature and then diluted with D_2_O buffer (10 mM Tris-HCl, 50 mM NaCl, pHread 7.1). HDX was carried out at room temperature and was quenched after 1 min, 2 min, or 10 min by adding quench buffer (4 M urea, 0.5 M TCEP-HCl in 1 M glycine, pH 2.3) and pepsin followed by 3 min incubation and rapid freezing in liquid nitrogen. Each sample was thawed and injected into an LC system (UltiMate 3000 RSLCnano, Thermo Scientific) where the protein was digested within a dual-protease enzymatic column (nepenthesin-1 and pepsin, 15 µl bed volume, Affipro s.r.o., CZ). Peptides were trapped and desalted on-line on a peptide microtrap column (Michrom Bioresources, Auburn, CA) for 3 min. Both digestion and desalting were provided by loading buffer (2% acetonitrile/0.05% trifluoroacetic acid) at a flow rate of 100 µl/min. Next, the peptides were eluted onto an analytical column (Jupiter C18, 0.5 x 50 mm, 5 µm, 300 Å, Phenomenex, CA) and separated by a 17 min two-stage linear gradient elution which was starting with 10% buffer B in buffer A and rising to 65% buffer B (began with 13 min stage 10-30% B followed by 4 min 30-65% B) at a flow rate of 50 µl/min. Buffers A and B consisted of 0.1% formic acid in water and 80% acetonitrile in 0.1% formic acid, respectively. The dual-protease, trap, and analytical columns were kept at 1 °C.

Mass spectrometric analysis was carried out using an Orbitrap Elite mass spectrometer (Thermo Fisher Scientific) with ESI ionization connected on-line to a robotic system based on the HTS-XT platform (CTC Analytics, Zwingen, Switzerland). The instrument was operated in a data-dependent mode for peptide mapping (HPLC-MS/MS). Each MS scan was followed by MS/MS scans of the three most intensive ions from both CID fragmentation spectra. Tandem mass spectra were searched using SequestHT against the cRap protein database (ftp://ftp.thegpm.org/fasta/cRAP) containing the sequence of ApoE3, ApoE4 (WT and MM) recombinant proteins with the following search settings: mass tolerance for precursor ions of 10 ppm, mass tolerance for fragment ions of 0.6 Da, no enzyme specificity, two maximum missed cleavage sites and no-fixed or variable modifications. The false discovery rate at the peptide identification level was set to 1%. Sequence coverage was analyzed with Proteome Discoverer version 1.4 (Thermo Fisher Scientific) and graphically visualized with the MS Tools application [45]. Analysis of deuterated samples was done in LC-MS mode with ion detection in the orbital ion trap. The MS raw files together with the list of peptides (peptide pool) identified with high confidence characterized by requested parameters (amino acid sequence of each peptide, its retention time, XCorr, and ion charge) were processed using HDExaminer version 2.2 (Sierra Analytics, Modesto, CA). The software analyzed protein and peptides behavior and created the uptake plots that showed peptide deuteration over time with a calculated confidence level (high and medium confidence are accepted, and low confidence is rejected). Each of the accepted peptides was mapped to the amino acid sequences of the analyzed proteins via the following procedure. Each residue was assigned the uptake data from any peptide solved with high confidence. Medium confidence peptides were also considered for positions without previously assigned data. Low-confidence peptides were rejected. The final uptake value (expressed as % of deuteration) assigned to each amino acid corresponded to the average of all assigned values for its position. HDX-MS measurements were mapped to 3D representations of proteins by assigning such averaged values to the PyMOL B-factor field of the corresponding amino-acid alpha carbon using data2bfactor.py script from Queens University, Canada (http://pldserver1.biochem.queensu.ca/∼rlc/work/pymol/), and visualized using putty representation. Amino acids with missing HDX-MS information were represented with minimal width and grey color.

The mass spectrometry proteomics data have been deposited to the ProteomeXchange Consortium via the PRIDE [46] partner repository with the dataset identifier PXD036416; Username, Password: UcoTxYaM.

### Analytical Ultracentrifugation

Analytical ultracentrifugation (AUC) experiments were performed using ProteomeLab XL-I analytical ultracentrifuge (Beckman Coulter, Indianapolis, IN, USA) equipped with an An-60 Ti rotor. Sedimentation velocity experiments were conducted in 12mm titanium double-sector centerpiece cells (Nanolytics Instruments, Potsdam, Germany) loaded with 380 µL of both protein sample and reference solution (10mM Tris/HCl pH 7.5, 50 mM NaCl). Data were collected using absorbance and interference optics at 20 °C at a rotor speed of 50,000 rpm. Scans were collected at 280 nm in 5-min intervals and 0.003 cm spatial resolution in continuous scan mode. The partial specific volume of the protein and the solvent density and viscosity were calculated from the amino acid sequence and buffer composition, respectively, using the software Sednterp (http://bitcwiki.sr.unh.edu). The data were analyzed with the continuous c(s) distribution model implemented in the program Sedfit 15.01c [47]. For the regularization procedure, a confidence level of 0.68 was used. The plots of c(s) distributions were created in GUSSI 1.3.1 [48].

(MOLECULAR DYNAMICS)

### Ligand preparation for molecular dynamics experiments

The structure of SPA f The treatment with tramiprosate was performed using Avogadro 2 [49]. As expected at pH 7.4, SPA was considered deprotonated on both the sulfonate (SO_3_^-^) and carboxylate (COO^-^) groups, with a net charge of −2. The minimization step was performed by the Auto Optimization Tool of Avogadro, using the UFF force field [50] with the steepest descent algorithm. The resulting structure was then submitted to further optimization and calculation of their partial atomic charges using Gaussian 09 [51], with the Hartree-Fock method and 6- 31G(d) basis set in a vacuum. The *antechamber* module of AmberTools 14 [52] was used to extract the RESP charges of the ligand from the Gaussian output files. To prepare the PAR and RTF parameter files compatible with the CHARMM force field we used the SwissParam webserver [53].

### Preparation of the systems

The three-dimensional structure of the ApoE3 NTD was obtained from the RCSB Protein Data Bank [54] (PDB ID 1BZ4). The structure of the ApoE4 NTD was obtained in this work as described above (PDB ID 8AX9). PyMOL was used to generate the dimeric units from those structures with the symmetry information contained in the original PDB files. The two chains of the dimers were renamed as A and B. The crystallographic water molecules were maintained.

To determine the angle between chains A and B of the dimeric unit, we calculated the center of mass and the principal axis of inertia of the two chains. The final value of the angle corresponds to the angle between the principal axes of inertia of each chain.

The three-dimensional structure of the full-length ApoE3_M_ was obtained from the RCSB Protein Data Bank [54] (PDB entry 2L7B). This structure resulted from NMR experiments and contains 20 models. Only the first of these models was extracted and used as a starting point for molecular dynamics (MD). The structure of the full-length ApoE4_M_ was obtained by homology modelling with the SwissModel webserver [55], using the respective FASTA sequence and the PDB entry 2L7B (ApoE3_M_) as a template.

The following steps were performed with the High Throughput Molecular Dynamics (HTMD) [56] python scripts. The protein was protonated with PROPKA 2.0 at pH 7.4 [57]. Each protein was simulated free and with an excess of SPA ligands. 1) The free protein was solvated with a cubic box of TIP3P [58] water with the edges at least 25 Å away from the protein atoms, using the *solvate* module of HTMD. 2) For the proteins with SPA, we used a python routine in the preparation protocol of HTMD, to add 100 molecules of SPA randomly placed on a sphere with a radius of 3 Å further from any atom of the protein, and at least 5 Å away from each other; the system was then solvated with a cubic box of TIP3P water with the edges at least 5 Å away from the ligand atoms. Cl^-^ and Na^+^ ions were added to neutralize the charge of the system and get a final salt concentration of 0.1 M. Considering the resulting box volumes, the final concentrations varied between 1.0–2.1 mM of protein and 0.10–0.21 M of SPA. The topology of the system was built, using the *charmmbuild* module of HTMD, with the modified CHARMM36m [59] force field and the parameters for the modified mTIP3P [58] solvent model, and the respective parameters for the ligands. The CHARMM36m/mTIP3P combination is expected to provide more accurate ensembles for intrinsically disordered proteins, which is the case of the CTD of ApoE [60].

### System equilibration

Each system was equilibrated using the *Equilibration_v2* module of HTMD [56]. It was first minimized using the conjugate-gradient method for 500 steps. Then the system was heated to 310 K and minimized as follows: (i) 500 steps (2 ps) of NVT thermalization with the Berendsen barostat with 1 kcal·mol^-1^·Å^-2^ constraints on all heavy atoms of the protein, (ii) 1,250,000 steps (5 ns) of NPT equilibration with Langevin thermostat and same constraints, and (iii) 1,250,000 steps (5 ns) of NPT equilibration with the Langevin thermostat without any constraints. During the equilibration simulations, holonomic constraints were applied on all hydrogen-heavy atom bond terms and the mass of the hydrogen atoms was scaled with factor 4, enabling 4 fs time step [61–64]. The simulations employed periodic boundary conditions, using the particle mesh Ewald method for treatment of interactions beyond 9 Å cut-off, and the smoothing and switching of the van der Waals interaction for a cut-off at 7.5 Å [62].

### Adaptive sampling

HTMD was used to perform adaptive sampling to study (1) the dissociation process of the ApoE dimers, and (2) the conformational diversity of the monomeric full-length ApoE_M_. For that, we performed production MD runs of 50 ns (for the dimers) or 100 ns (for the monomers), which started from the systems that resulted from the equilibration cycle and employed the same settings as the last step of the equilibration. The trajectories were saved every 0.1 ns. Adaptive sampling was performed using the root-mean-square deviation (RMSD) of the C_αα_ atoms with respect to the initial protein structure of each system, and time-lagged independent component analysis (tICA) [65] in 1 dimension. Several epochs of 10 MDs each were performed for each system: 1) for the dimer, we ran the simulations until the dissociation of the dimers was observed or a maximum of 20 epochs was reached (corresponding to a cumulative time of 10 μs); 2) for the monomers, 20 epochs of 10 MDs were performed for each system, corresponding to a cumulative time of 20 μs.

### Markov state model construction

The adaptive simulations of the ApoE_M_ systems were made into a simulation list using HTMD [66], the water and ions were filtered out, and unsuccessful simulations with lengths less than 100 ns were omitted. This resulted in 20 µs of cumulative simulation time (200 × 100 ns). The ApoE conformations were studied by the same metric used in the adaptive sampling: the RMSD of the C_αα_ atoms of the protein with respect to the initial structure. The data were clustered using the MiniBatchKmeans algorithm to 1000 clusters. A 30 ns lag time was used in the models to construct 4 Markov states, and the Chapman-Kolmogorov test was performed to assess the quality of the constructed states.

### Analysis of MD trajectories

The ParmEd program [67] was used to convert the CHARMM topologies to AMBER topologies. The *cpptraj* [68] module of AmberTools 14 [52] was used to concatenate the filtered trajectories of each system, ordering the epochs chronologically, center and align them by their backbone atoms, and save the combined trajectory in a single file. *Cpptraj* was also used to calculate the B-factors of the protein backbone atoms, dihedral angles of some residues, and the angles formed by some atoms in helix 3. The *do_dssp* module of Gromacs 5.1 [69], patched with DSSP 3.0 [70], was used to compute the secondary structures for every snapshot of the combined trajectories to obtain the total secondary content.

An in-house script that applies DSSP 3.0 [70] was used to calculate the secondary elements by residue for an ensemble containing 10 ns-spaced snapshots (every 100^th^ frame) of each simulation. The analyzed snapshots were subsequently grouped in ten different bins. The average secondary structure type content values were calculated for each bin, and the dispersion of each secondary structure type value was reported as the standard error of the mean (as a proxy of the certainty of the content value calculated for that particular secondary structure type).

### Interaction energies

The linear interaction energy (LIE) [71] was computed to assess the free binding energy of the ApoE dimers with the SPA molecules, expressed as the respective electrostatic and van der Waals components, in the adaptive simulations of the dimers of ApoE3 and ApoE4. For that, we used *cpptraj* to calculate the LIE on every snapshot of the respective combined MD trajectories of the adaptive simulations. This method was also used to assess the contribution of each residue to hold the dimers together. For that, we computed the interaction energy of every residue in each chain with the whole opposite monomeric unit. In both calculations, we evaluated the interactions when the dimers were still associated, and for that, we considered only the first 500 ns of the concatenated MD.

(CELL BIOLOGY EXPERIMENTS)

### Cell culture of induced pluripotent stem cells (iPSCs)

Two human iPSC lines used in this study were passaged and maintained using standard feeder-free culture protocols. In brief, feeder-free cultures were grown on Matrigel-coated plates (Corning) in mTeSR^TM^1 medium (STEMCELL Technologies) supplemented with half of the recommended dose of ZellShield® (Minerva Biolabs). Cells were passaged using 0.5 mM EDTA (Thermo Fisher Scientific) in PBS or manually. The “sAD-E4” cell line was derived from a patient with a sporadic form of AD with ApoE4/4 status. The isogenic cell line “sAD-E3” was obtained by correction of ApoE4/4 to ApoE3/3. Both isogenic iPSC lines were kindly provided by Dr. Li-Huei Tsai and described previously by Lin and co-workers [72].

### Cerebral organoid culture and treatment with tramiprosate

Cerebral organoids were generated using the protocol described previously [73,74]. Briefly, for spheroid formation, cells were plated at day 0 into non-adherent V-shaped 96-well plates at 2000-3000 cells in 150 μl of mTeSR^TM^1 medium with 50 μM ROCK inhibitor (S1049, Selleckchem). Plates were centrifuged for 2 min at 200 g to facilitate spheroid formation. Non-adherent cell culture plates were prepared with poly(2-hydroxyethyl methacrylate) (poly-HEMA; P3932, Merck) coating; 3) On day 2, the cell culture medium was exchanged for fresh mTeSR^TM^1 without ROCK inhibitor. When spheroids reached the size of 400 - 600 µm, fresh Neural Induction Medium [74] was added every day for six days (usually from day 3 to day 8). The next day, twelve organoids were transferred to one 6 cm cell culture dish. The remaining medium was aspirated, and dry organoids were embedded in 7 µl of cold Geltrex^TM^ (Thermo Fisher). Geltrex^TM^ was left to solidify as hanging drops on the inverted cell culture dish for 10 min at 37 °C. Solidified Geltrex^TM^ drops with organoids were gently detached from the bottom and cultured without shaking in Cerebral Organoid Differentiation Medium (CODM) [74] without vitamin A for seven days. Subsequently, organoids were cultured in CODM with vitamin A and were moved on an orbital shaker on day 26 (± two days). CODM was changed three times a week by aspirating at least half of the old medium and replacing it with a fresh medium. ZellShield® (Minerva Biolabs) was used as the contamination preventive in all media.

The treatment with tramiprosate was performed for 50 days, from day 50 to day 100. During the treatment, CODM was continuously supplemented with 100 μM tramiprosate. Samples were harvested from two independent batches of CO differentiation. Control samples were treated with solvent only.

### Cryo-sections and immunohistochemistry (IHC)

Harvested cerebral organoids were fixed with 3.7% paraformaldehyde for one hour and washed with PBS. For cryo-sections, fixed organoids were saturated with 30% sucrose (Merck), embedded in O.C.T. medium (Tissue-Tek), and frozen. 10 µm sections were prepared on cryostat Leica 1850. Excessing O.C.T. medium was removed by 15 min PBS wash prior to IHC staining. Sections were permeabilized in 0.2% Triton-X (Merck) in PBS and blocked in 2% normal goat serum (Merck) in permeabilization solution. They were then incubated with primary antibodies diluted in blocking solution at 4 °C overnight, followed by the incubation with secondary antibodies (Alexa FluorTM; Thermo Fischer Scientific) for 1 hour at room temperature. Nuclei were visualized by Hoechst 33342 (Thermo Fischer Scientific). Following primary antibodies were used: PAX6 (#60433; Cell Signaling Technologies - CST), TUJ (#4466; CST), MAP2 (#8707; CST), GFAP (#12389; CST), IBA1 (#019-19741; Labmark), S100β (ab11178; Abcam), ApoE (NBP2-49450; NovusBio), DCX (sc-271390; Santa Cruz Biotechnology).

### Western blotting

Protein lysis and Western blot were performed as described previously [75]. Briefly, protein samples were lysed in 1% SDS-lysis buffer, and concentration was measured with DC™ Protein Assay (Bio-Rad). Samples were then mixed with 10x Laemmli buffer and incubated at 95°C for 10 min. Proteins were separated on 10% Acrylamide gel, transferred onto PVDF membranes (Merck), blocked, and subsequently incubated with antibodies in 5% skimmed milk or BSA. The results were visualized via ECL™ Prime (Amersham) using ChemiDoc™ Touch Imaging System (Bio-Rad). Densitometric analysis was done using ImageJ software and plotted as a bar graph with individual values – each dot represents a sample of 2-3 organoids; 2 samples from each batch were analyzed (outliers were identified by Grubbs test, α=0.01 and excluded); 2 independent batches of organoids were used for this study. Following primary antibodies were used: ERK (#4695; CST), P-ERK (#4370; CST), AKT (#9272; CST), P-AKT (#9271; CST), β-Actin (#3700; CST).

### Sample preparation for metabolomic, proteomic, and lipidomic assays

For metabolomic, proteomic, and lipidomic assays, individual organoids on day 100 were transferred to 1-1.5 ml of E6 minimal medium (Thermo Fischer Scientific) for 72 hours. Before proteomic and lipidomic analyses, organoids were washed with PBS, incubated in Cell Recovery Solution (Corning) for 1 hour, washed with PBS, and stored at −80 °C. For metabolomic analysis, cerebral organoid conditioned E6 medium was transferred to individual tubes, centrifuged to remove cell debris, and frozen at −80 °C. 5-6 organoids from two individual batches were harvested for all analyses.

<u>LIPIDOMICS, GANGLIOSIDE AND METABOLITE ANALYSIS</u>

### Materials and reagents

Ammonium acetate (Cat# A11450), and ammonium formate (Cat# A11550) were purchased from Fisher Chemical (New Hampshire, USA). LC-MS grade acetonitrile (Cat# 34967), isopropanol (Cat# 34965), and formic acid (FA) were purchased from Honeywell (Charlotte, USA). LC-MS grade methanol (Cat# 0013687802BS) was purchased from Biosolve Chimie (Dieuze, France). Acetic acid (≥99.8%) was purchased from Penta Chemicals (Chrudim, Czech Republic). Lipidomics standards SPLASH Lipidomix (cat. #330707) and Cer/Sph Mixture I (cat. #LM6002) were purchased from Avanti Polar Lipids (Alabaster, USA). 3- sulfopropionic acid (cat. #BD01044758) was purchased from BLD pharma. Tramiprosate (cat. # A76109) was purchased from Sigma Aldrich. GM1 and GM3, isotopically-labeled ganglioside (GS) internal standards (IS), were synthesized in-house.

### Lipids, gangliosides, and metabolite extraction

Cerebral organoids (COs) were washed with PBS, treated for 1 hour with cell recovery solution (CRS, Corning, NY, USA) at 4 °C, and washed with PBS again. CO was homogenized in a microtube with a glass bead (Benchmark Scientific, Edison, NJ, USA). The COs were stored at −80 °C before processing. COs were freeze-dried (Savant SDP121 P, SpeedVac, ThermoFisher Scientific, USA) and homogenized using a glass bead. Lipid, ganglioside, and metabolite extraction were performed by adding 100 µl of 80% isopropanol to the homogenate. It was followed by a brief vortex, sonication (37Hz, 5 min), and vortexing (200 rpm, 10 min). The extract was then centrifuged (12.3 RCF, 5 min), and the filtrate was removed and mixed 1:1 with a mixture of lipid and metabolite internal standards (**Supplementary Table 2**) for lipidomics, and metabolite assays or with a mixture of isotopically labeled GM1 and GM3 internal standards for ganglioside assay, and stored in −20 °C until LC-MS analysis. The protein pellet was dried using the SpeedVac vacuum concentrator (Savant SDP121 P, ThermoFisher Scientific). The dried protein pellet was reconstituted to perform the BCA assay protein determination. Control (N=10) and tramiprosate-treated COs (N=10) were utilized for the selective reaction monitoring - mass spectrometry (SRM-MS) analysis. The protein pellet was dried to be processed for the proteomics assay.

### Lipidomics assay and data processing

The cerebral organoid extract was analyzed using a 1290 Infinity II UHPLC (Agilent) system coupled with the 6469 Triple Quadrupole mass spectrometer (Agilent). 1 µl of lipid extract was injected twice on the reverse phase microbore column (CSH, 1 mm *100 mm, 1.7µm, Waters), separated at 100 µl/min flow rate over 15 min. For the gradient elution, mobile phase A was 10 mM ammonium formate in acetonitrile: water (60:40), and mobile phase B was 10 mM ammonium formate in isopropanol: acetonitrile (90:10). The gradient elution program was: 0 min 15 % B, 1.86 min 30% B, 2.32 min 48%, 9.5 min 82% B, 12.5 to 13.5 min 99% B and 13.5 to 15 min column re-equilibration. The positive ion mode source parameters were: gas temp 200 °C, gas flow 14 l/min, nebulizer pressure 45 psi, sheath gas temp 400 °C, and sheath gas flow 8 l/min capillary voltage 4 kV, nozzle voltage 500 V, and unit resolution for Q1 and Q3. Data were acquired using the dynamic SRM mode [76–78], with a 2 min retention time window per transition. Raw data files were processed using Mass Hunter Quantitative analysis (B.07.00, Agilent Technologies) software. The concentrations of lipid species were calculated from the respective internal standard and further normalized to total lipid content to account for the variable sample amount.

### Metabolite (tramiprosate and SPA) assay and data processing

The cerebral organoid extract was analyzed on Agilent 1290 UHPLC coupled with the 6469 QQQ (Agilent) system. Each sample was injected twice (2µl) for the hydrophilic interaction chromatographic (HILIC) separation on BEH Amide Column (2.1×100 mm, 1.7 µM, Waters Corporation, Milford, MA) at 0.4 ml/min flow rate. Mobile phase A was Milli-Q water containing 10mM ammonium acetate (pH = 9) and mobile phase B was 95% acetonitrile containing 10 mM ammonium acetate (pH = 9), ammonium hydroxide was used to adjust the pH. The total run time was 12 min. The gradient elution program was: 0 min 99% B, 0.1 min 99% B, 6.0 min 70%, 6.5 min 99% B, and column re-equilibration till 12 min. The negative mode jet stream source parameters were gas temp 190 °C, gas flow 14 l/min, nebulizer pressure 25 psi, sheath gas temp 400 °C, and sheath gas flow 10 l/min capillary voltage 4 kV, nozzle voltage 1500 V, and unit resolution for Q1 and Q3. The SRM transitions was optimized injecting the respective metabolite standards, tramiprosate (138>80) and SPA (153>81). The SRM peaks were integrated using Agilent MassHunter Quantitative Data Analysis (Santa Clara, CA). Relative response factor calculated to perform the quantification of tramiprosate and SPA. Metabolite concentration further normalized to total protein content.

### Gangliosides assay and data processing

The Agilent UHPLC system was equipped with a C18 CSH column (CSH 50 x 2.1 mm x 1.7 μm, Waters Corp., MA, USA). Positive ion mode mobile phase A consisted of 0.5 mM ammonium formate in water. Mobile phase B consisted of methanol and isopropanol (50:50 v/v). For the analysis, the lipid extracts were diluted two-fold with the addition of 0.3 μM of isotopically labeled GM1 and GM3. Gangliosides were separated during a linear gradient elution shifting from mobile phase B 30% to 70% over 2 min and then to 95% over 9 min. The column was conditioned with 95% B from 9 to 13 min and then 95% A for an additional 1.5 min. Re-equilibration was done from 14.5 to 17.1 min. Negative mode mobile phase A consisted of 0.5 mM ammonium formate and 10 mM ammonium acetate in water and B of acetonitrile and isopropanol (50:50 v/v). The gradient elution profile began at 10% B for 4 min and increased to 85% B till 6.2 min, then to 95% B to 10.4 min. The column was flushed with 10% B from 10.4 to 14.4 min and 95% B from 14.4 to 16.4 min, with re-equilibration from 16.4 – 19.1 min at 10% B. A flow rate of 300 µl/min was used for all separations. The injection volume was 2 µl in negative mode and 3 µl in positive mode.

The GS assay was performed on a triple quadrupole mass spectrometer (Agilent 6495B, Agilent Technologies) operated in positive and negative ion mode. The capillary voltage was 3.5 kV and 3 kV, respectively, in positive and negative ion modes. The selected reaction monitoring (SRM) mode. The SRM library for the quantification of gangliosides is given in **Supplementary Table 3**. Additionally, the gas flow rate of 16 l/min at 190 °C with sheath gas pressure at 20 psi and 350 °C was utilized for positive mode. In negative mode, the gas flow rate was 14 l/min at 190 °C with sheath gas pressure at 25 psi and 400 °C.

We processed raw data in the Skyline (MacCoss Lab., UW, USA). The labeled GM3 IS was used to determine concentrations for all GSs species, except for GM1, which utilized the labeled GM1 IS. Respective response factors (RF) were calculated for all GS species for which accurate IS was unavailable. All concentrations were reported as the average of two technical replicates and normalized to the total protein level. Graphical representation and Mann-Whitney tests were performed using GraphPad Prism version 8.0.0 for Windows (GraphPad Software, San Diego, California, USA, www.graphpad.com).

<u>PROTEOMICS</u>

### Materials and reagents

Ammonium bicarbonate (AmBic, BioUltra, ≥99.5% purity, 09830-500G), sodium deoxycholate (SDC, BioXtra, ≥98.0% purity, 30970-25G), iodoacetamide (IAA, ≥99% purity, I6125-5G) were obtained from Sigma Aldrich (St. Louis, MO, USA), 1,4-dithiothreitol (DTT, ≥99% purity, Art.-Nr. 6908.1) from Carl Roth GmbH + Co. KG (Karlsruhe, Germany). Trypsin gold, Mass Spec Grade, was from Promega (Madison, WI, USA). Synthetic isotopically labeled (SIL) peptide standards (SpikeTides_L crude) were purchased from JPT Peptide Technologies Inc. (Acton, MA, USA). Pierce BCA Protein Assay Kit reagents were from ThermoFisher Scientific, Waltham, MA, USA. The ultrapure water was prepared in the purification system (arium® Comfort System, Sartorius).

### Protein pellet solubilization and enzymatic proteolysis

Briefly, after lipid extraction, the dried protein pellet with a glass bead was solubilized in 100 μl AmBic buffer (100 mM) with SDC (5 mg/ml)39. Samples were vortexed (10 s, 2000 rpm, VELP Scientifica), homogenized (4 m/s, 10 s, two cycles with 10 s inter-time, BeadBlasterTM 24, Benchmark), sonicated (1 min, 80 kHz, Elmasonic P, Elma Schmidbauer GmbH), mixed (10 min, 2035 rpm, HeidolphTM MultiReax) and centrifuged (1 min, 12300 RCF, Micro-Star 12, VWR®, Radnor, PA, USA). The total protein concentration in each sample was determined using the BCA assay in 10-20 μl sample aliquots, and the protein concentration was adjusted to 0.1-0.3 μg/μl by adding the solubilization buffer. Samples were centrifuged, and aliquots of 6-18 µg of total protein (60 μl) were reduced (20 mM DTT in 50 mM AmBic; 10 min; 95 °C) and alkylated (40 mM IAA in 50 mM AmBic; 30 min; room temperature in the dark). The remaining sample homogenates from individual COs were mixed to prepare a QC sample for calibration curve measurements following the same protocol as for individual COs. Trypsin was added in the ratio of 1:20-40 (enzyme: total protein content, w/w), and the Parafilm®-sealed samples were incubated (37 °C; 16 h; gentle shaking). The isotopically labeled synthetic peptides were added (final sample conc. ≈22-29 nmol/l) after quenching the digestion with 200 µl of 2% FA. Samples were loaded on the mixed-mode cartridge (Oasis® PRiME HLB – 30 mg, Waters Corp. Milford, MA, USA) for solid-phase extraction (SPE). Peptides were washed with 2% FA, eluted with 500 µl of 50% ACN with 2% FA and dried in SpeedVac. The sample preparation details are provided in Supplementary data (Sample processing details).

### Mass spectrometry protein assays and data processing

The dried, purified peptides were reconstituted in 15-20 µl of 5% ACN with 0.1% FA. Peptides were analyzed in positive ion detection mode using the same UHPLC-MS/MS system as described for metabolite analysis. Samples (3 µl) were injected into the C18 analytical column (Acquity UPLC CSH 1.7 µm, 2.1 mm x 100 mm, Waters Corporation, Milford, MA, USA). The mobile phase flow rate was 0.3 ml/min; buffer A (0.1% FA) and buffer B (0.1% FA in 95% ACN). Linear gradient elution: initial 5% B; 25 min 30% B; 25.5 min 95% B; 30 min 95% B; and from 31 to 35 min with 5% B. The ESI source temperature was 200 °C, and the capillary voltage was 3500 V. We used a multiplex SRM assay for monitoring of 32 proteins detectable in cerebral organoids (**Supplementary Table 4**). The nextprot.org database was used to select unique peptide surrogates for targeted proteins. Srmatlas.org (online) was used to generate SRM assays preferentially based on experimental data. We used a dynamic SRM mode with a 2 min window centered at a peptide experimental retention time. Data were manually inspected and processed in Skyline software (version 21.2.0.369, MacCoss Lab, UW, USA). Transitions corresponding to (>5) y ions with a reproducible signal (on average, %CV < 15 %, n = 2) were selected for the relative protein quantification (**Supplementary Table 5**). The average value of the two technical replicates was used for further normalization. Relative concentrations determined using quantifier transition of standard (ST) and light peptide peak area (light peptide peak area/ST peptide peak area*ST peptide concentration) were normalized to the GAPDH levels. The final values (Supplementary data: Expression levels of proteins) in tramiprosate-treated organoids were related to the corresponding non-treated organoids (ApoE ε3/ε3 and ApoE ε4/ε4) in each batch (Supplementary data: Protein levels normal. to NTR) to see the effect of tramiprosate relatively to organoids without the treatment.

## Results

### ApoE4 prefers a geometrically different crystal packing than ApoE2 and ApoE3

Despite the crystallization of full-length ApoE4, we could only obtain ApoE4 NTD structures (residues ∼22-165), but with well-defined loops conformation (**Supplementary Table 6**). Although our four new ApoE4 crystals belonged to two different space groups (orthorhombic P2_1_2_1_2_1_ versus trigonal P3_1_21), they shared similar crystal packing contacts. Specifically, during the comprehensive analysis of the previously solved ApoE crystal structures and our novel ones, we observed that all three ApoE isoforms shared similar, but not identical, crystal packing patterns (**Figure 1A, Supplementary Figure 1** and **Supplementary Table 7**). The crystal lattices consisted of a unique protein-protein contact region that stabilized the regular arrangement of ApoE molecules. Two NTDs (chains A and B) formed a T-shaped or V-shaped molecular building dimeric unit (**Figure 1B**) that stacked with another adjacent dimeric unit, resulting in an elongating crystallographic fibril. While the angle between chains A and B in the T-shaped dimeric unit was ∼100°, the corresponding angle in the V-shaped dimeric unit was significantly smaller ∼75°. This ∼25° angle difference between T-shaped and V-shaped packing was retained in both the orthorhombic and trigonal space groups.

**Figure 1.**
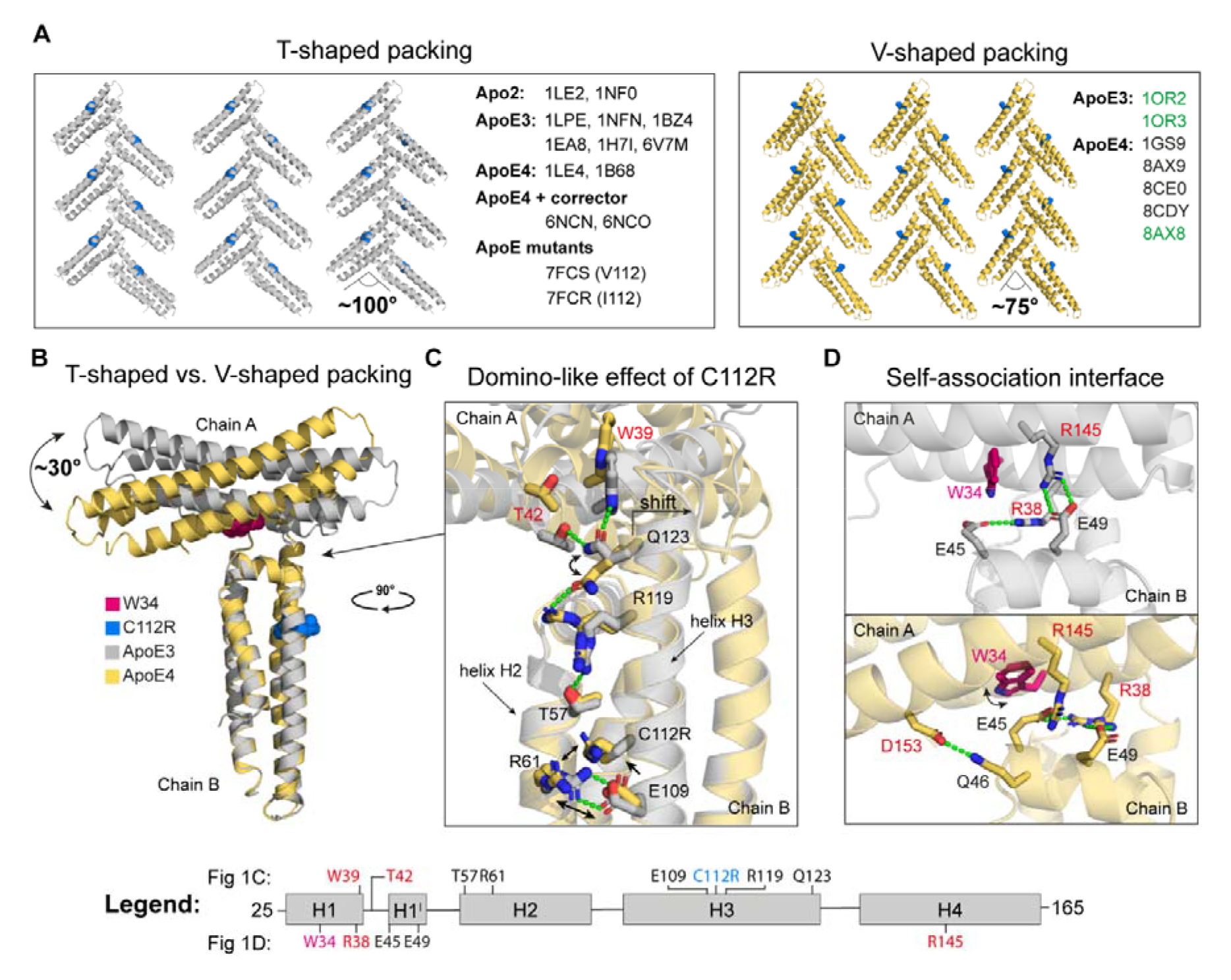
Structural insight into the ApoE crystal packing topologies. **(A)** Comparison of T-shaped (grey) and V-shaped (yellow) crystal packing patterns of all inspected ApoE structures with position 112 visualized as a blue sphere. ApoE structures with PDB identification codes marked in green have a V-shaped packing but residue organization at the self-association interface as in the T-shaped packing. **(B)** Superposition of ApoE3 (grey) and ApoE4 (yellow) NTD dimeric units with a marked W34 (pink sphere) and R112 (blue sphere). **(C)** “Domino-like” effect of C112R substitution leading from small local changes up to a change in the tilt angle between chain A and B. **(D)** Comparison of the self-association interface of ApoE3 (the upper panel) and ApoE4 (the lower panel). The critical interaction residues of chain A (W34, R38, R145, and D153) and chain B (E45, Q46, and E49) are in black and red, respectively.

A comparison of previously solved ApoE structures revealed that ApoE2 and ApoE3 prefer T-shaped packing, as 2 out of 2 ApoE2 structures and 6 out of 8 ApoE3 structures display T-shaped packing. On the other hand, 5 of the 7 ApoE4 structures have V-shaped packing, indicating an ApoE4 preference for this packing pattern. However, when the ApoE4 was complexed with the previously designed “corrector” molecules [29], it crystallized in the T-shaped packing pattern. On the other hand, both ApoE variants V112 and I112 (PDB IDs: 7FCS and 7FCR) display T-shaped packing, suggesting that the V-shaped packing is a consequence of the C112R substitution.

### The C112R substitution triggers long-distance structural changes in ApoE4

The C112R substitution in ApoE4 (**Figure 1C** and **Supplementary Figure 2**) led to the formation of a new salt bridge between R112 and E109 and the repositioning of R61. Consequently, the helix H2-H3 interaction between R61 and E109 observed in the T-shaped packing was abolished, and the four-helix bundle was destabilized. The “domino-like” effect of the C112R mutation was further exacerbated through the H3 helix up towards Q123, which directly influenced a tilt between chains A and B. The side-chain of Q123 was oriented upwards in the T-shaped packing, while it was rotated downwards in the V-shaped packing. The upward orientation of Q123 in chain B was stabilized by interactions with W39 and T42 of chain A. In contrast, the absence of this interaction in ApoE4 structures resulted in a tilt (∼30°) between chains A and B and the subsequent formation of a geometrically distinct V-shaped dimeric unit (**Figure 1B** and **Video** in **Supplementary data**).

One of our ApoE4 structures (PDB ID: 8AX8) further revealed that R119 could adopt two conformational states. The “downward-facing” conformation of R119 was found in all previously solved ApoE structures. In the newly observed “upward-facing” conformation, R119 formed a stabilizing interaction with Q123, locking it in an orientation at which Q123 could no longer interact with W39 and T42 (**Figure 1C**). Despite a different tilt in T-shaped and V-shaped dimeric units, the main self-association interface formed by chains A and B remained the same in all ApoE isoforms (**Figure 1D**). The key interacting residues that held the two chains together were R38 (H1) and R145 (H4) from chain A and E45 and E49 located in helix 1’ (H1’) of chain B. Additionally, a new specific interaction in ApoE4 was formed between D153 from chain A and Q46 from chain B as both chains were tilted in the ApoE4 structure.

Moreover, E45 which interacts with R38 and R145, moved to the center of the “self-association interface”, causing a re-orientation of the W34 side chain upwards. This “flip-out” orientation of W34 in the parallel direction to the chain A axis is typical for ApoE4 preferred V-shaped packing. In contrast, the “flip-in” orientation of the W34 side chain in the direction perpendicular to the chain A axis was observed in the ApoE2 and ApoE3 preferred T-shaped packing. Interestingly, the W34A mutation led to an inversion of the fluorescence signal during the thermal denaturation of ApoE_M_, which suggests that W34 and thus the “self-association” interface may be involved in the aggregation process of ApoE (**Supplementary Figure 3**).

### Destabilization of the helix H2-H3 interaction by the loop that connects them

Contrary to the domino-like effect of C112R substitution described above, 2 of the 8 ApoE3 structures (PDB IDs: 1OR2 and 1OR3) have V-shaped packing. When we compared these ApoE3 structures with the ApoE4 structures, we noticed a conformational change in the loop that connects the H2 and H3 helices (**Figure 2**). While this loop was formed by residues 81-88 in ApoE4, it was extended by three residues (81-91) in the case of these ApoE3 structures. The unwinding of the beginning of helix 3 (residues 89-91) probably led to a weakening of the interaction between helices H2 and H3 and, consequently, to conformational changes similar to those caused by the C112R substitution. Although these ApoE3 structures exhibit the V-shaped packing preferred by ApoE4, the orientation of the residues at the self-association interface is similar to that of the T-shaped packing, suggesting a sort of intermediate state between T-shaped and V-shaped packing (**Figure 2**). We found the same conformational arrangement in our trigonal form of ApoE4. Interestingly, in our simulations, we also observed the unwinding of the beginning of helix H3 in ApoE3, whereas in ApoE4, there was the unwinding of helix H3 at the opposite end, confirming the domino-like effect of the C112R substitution. (**Supplementary Figure 4**).

**Figure 2.**
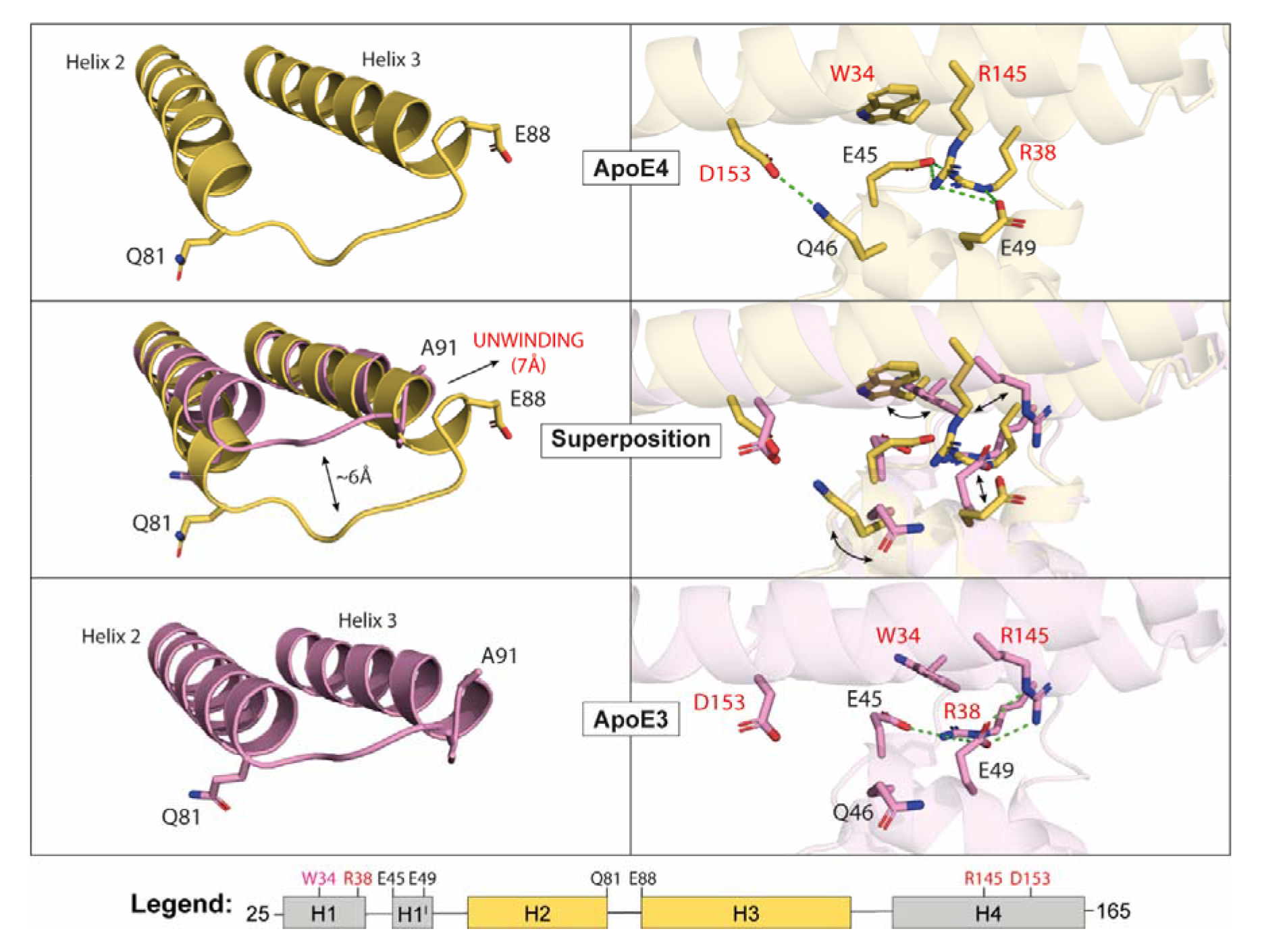
Comparison of V-shaped packing in ApoE4 and ApoE3 structures. The conformational change of the loop connecting helices H2 and H3 in ApoE3 structures (PDB IDs: 1OR2 and 1OR3) leads to V-shaped packing but to a different residue arrangement at the self-association interface than in ApoE4 structures.

### The self-association interface is involved in ApoE aggregation

Next, we strove to link the self-association interface with the ApoE aggregation propensity. Using the static light scattering at 37 °C, we observed increased aggregation of ApoE4 compared to ApoE3, as described previously [33,34]. Moreover, we observed that the five mutations previously introduced into the ApoE code to monomerize both isoforms (ApoE_M_) simultaneously inhibit their aggregation (**Supplementary Figures 5 and 6**). Based on these observations, we hypothesized that if our proposed self-association interface is involved in ApoE aggregation, it will not be possible to model a T-shaped dimer from ApoE3_M_. Therefore, we created a full-length T-shaped ApoE3_M_ dimer structural model and analyzed whether it contains any clashes (**Figure 3A**). In this model, the two CTDs are distant and do not interact. However, the C1 helix (residues 204-223) in both chains appears to play an important role in the interchain interactions. The helix C1 of chain B is in direct contact with the NTD of chain A within the self-association interface, whereas the helix C1 of chain A is close to the end of HD2 and helix H3 of chain B containing Q123 (**Figure 3B**). Importantly, helix C1 of chain B is in a clash with helices N1 and N2 (residues 1-23) of chain A. In the monomeric structure of ApoE3_M_, these helices are structurally adjacent to the CTD end that carries the five monomerizing mutations. Therefore, we investigated whether the position of these helices can be affected by the introduced “monomerizing” mutations. The superposition of the ApoE3_M_ structure with the NMR structure of the ApoE3 NTD (PDB ID 2KC3) showed that the position of the helices N1 and N2 is indeed different (**Figure 3C**). This implies that the monomerization of ApoE_M_ and the fact that ApoE_M_ does not aggregate may be caused indirectly by the mutations displacing the helices N1 and N2 to a new position.

To further support the involvement of the self-association interface in ApoE aggregation, we expected that the removal of helices N1 and N2 should lead to ApoE_M_ aggregation. To test this hypothesis experimentally, we created two truncated ApoE_M_ variants: (i) aggregation-prone ApoE_M_-AP with truncated helices N1 and N2 (residues 1-23), and (ii) aggregation-suppressed ApoE_M_-AS with the same truncation, but carrying additional four mutations (R38A, E45A, E49A, and R145A). These mutations were introduced in order to impair the self-association interface of the dimeric unit (**Figure 3D**). Both ApoE_M_ variants displayed α-helical content indicating their proper folding (**Figure 3E**). However, the dynamic light scattering revealed the presence of higher structures in the case of ApoE_M_-AP (**Figure 3F**). The aggregation of ApoE_M_-AP was so strong that the sample became turbid immediately after purification by size exclusion chromatography (**Figure 3G**). Interestingly, while the aggregation of ApoE4_M_-AP occurred almost instantaneously, it took several hours to observe the aggregation of ApoE3_M_-AP.

**Figure 3.**
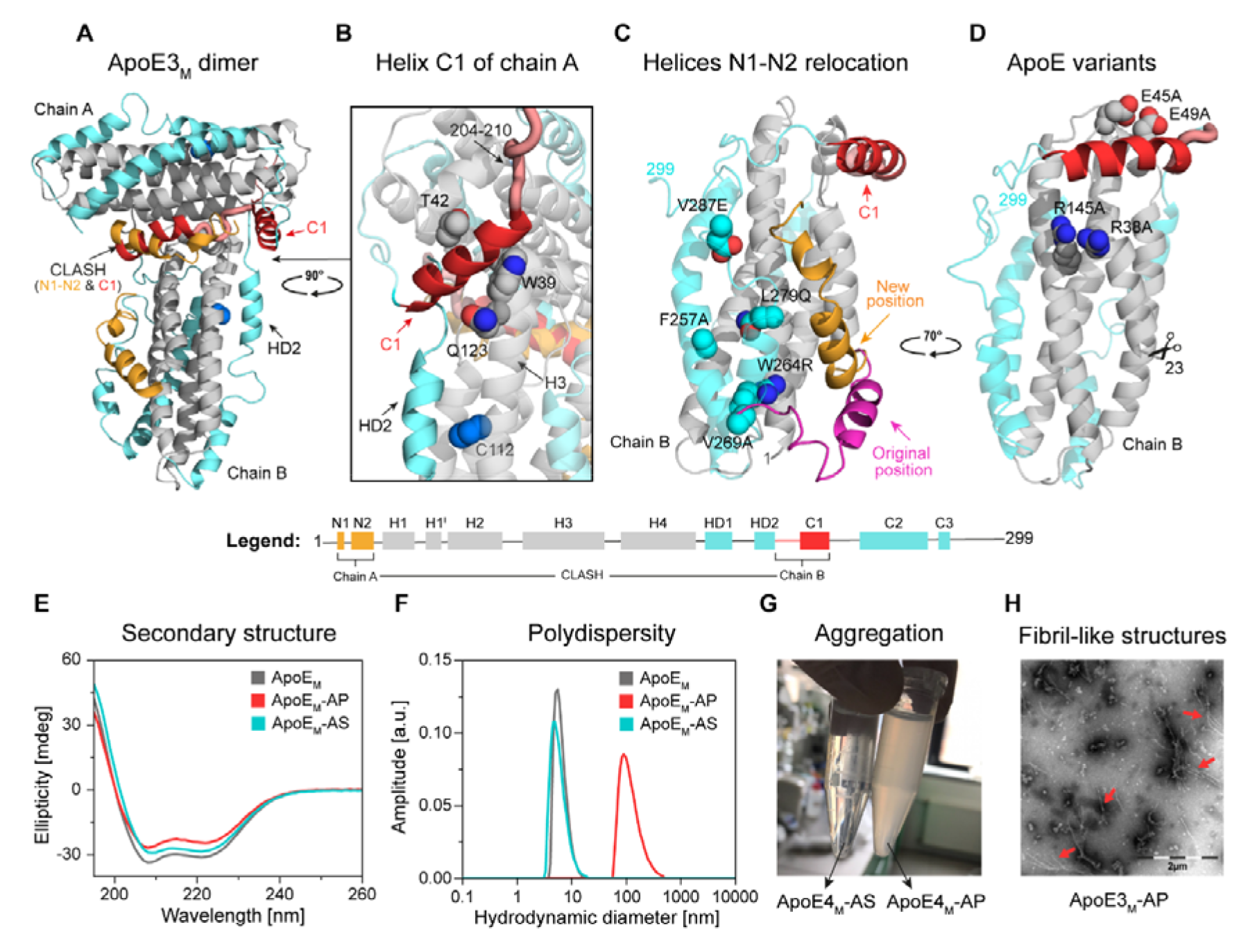
Structural determinants of ApoE self-association. **(A)** The model of the full-length ApoE3_M_ T-shaped dimer is colored according to the legend, with C112 marked as a blue sphere. **(B)** Detailed view of helix C1 of chain A located near residues W39, T42, and Q123 (spheres). **(C)** Superposition of NMR structures of ApoE3_M_ (PDB ID 2L7B; colored according to the legend) and NTD ApoE3 (PDB ID 2KC3; residues 1-23 and 24-165 are shown in pink and grey, respectively). The five “monomerizing” mutations (F257A, W264R, V269A, L279Q, and V287E) in the CTD of ApoE3_M_ are shown as spheres. **(D)** The structure of truncated ApoE_M_ with marked four mutations (R38A, E45A, E49A, and R145A) was introduced at the self-association interface. **(E)** Circular dichroism spectra and **(F)** dynamic light scattering of ApoE_M_ (grey), ApoE_M_-AP (red), and ApoE_M_-AS (cyan). The data represent averages of twenty individual repeats. **(G)** Photo showing soluble ∼ transparent ApoE_M_-AS (left) and an aggregated ∼ turbid ApoE_M_-AP sample two hours after purification (right). Similar results were obtained in at least two independent experiments. **(H)** TEM of ApoE3_M_- AP fibrils (red arrows). Similar results were obtained in at least two independent experiments.

Subsequent transmission electron microscopy (TEM) analysis of these aggregates revealed fibril-like morphology in both cases (**Figure 3H**). Conversely, no aggregation was observed for ApoE3_M_-AS and ApoE4_M_-AS. These results imply that removing the helices N1 and N2 clashing in the modelled T-shaped dimeric unit results in protein aggregation. The alanine substitution of four key interacting residues at the self-association interface suppressed this aggregation, strongly supporting our hypothesis that the self-association interface is involved in the early stages of ApoE aggregation.

We conducted extensive hydrogen-deuterium exchange coupled with mass spectrometry detection (HDX) experiments and molecular dynamics (MD) simulations of different ApoE structures to further test this proposal. Based on LC-MS/MS bottom-up proteomics analysis of both ApoE_WT_ and ApoE_M_, we found that wild-type ApoE3 and ApoE4 isoforms lack peptides in NTD corresponding to residues ∼ 31-50 and 44-52, respectively (**Figure 4A,** CTD in **Supplementary Figure 7**). Interestingly, this region correlates with the self-association interface used by dimeric units in the crystallographic lattices. This implies that these regions in the homotetrameric form of ApoE are covered and inaccessible to proteases. The main differences between wild-type ApoE3 and ApoE4 isoforms in HDX-MS analysis were the deuterium uptakes (**Figures 4B** and **Supplementary Figure 8**) observed around position 112, including helix H3 (residues 79-123) and helix H4 (residues 134-136). Moreover, some differences were observed for the CTD, suggesting that C112R substitution has a long-range impact on the entire ApoE molecule and its tetramerization. The sequence coverage was complete for the NTD of the ApoE3_M_ detected by mass spectrometry after enzymatic digestion, as opposed to the ApoE_WT_ (**Figure 4A**). This observation strongly supports our hypothesis that monomerization by five-point mutations in the CTD interrupts this tight packing.

**Figure 4.**
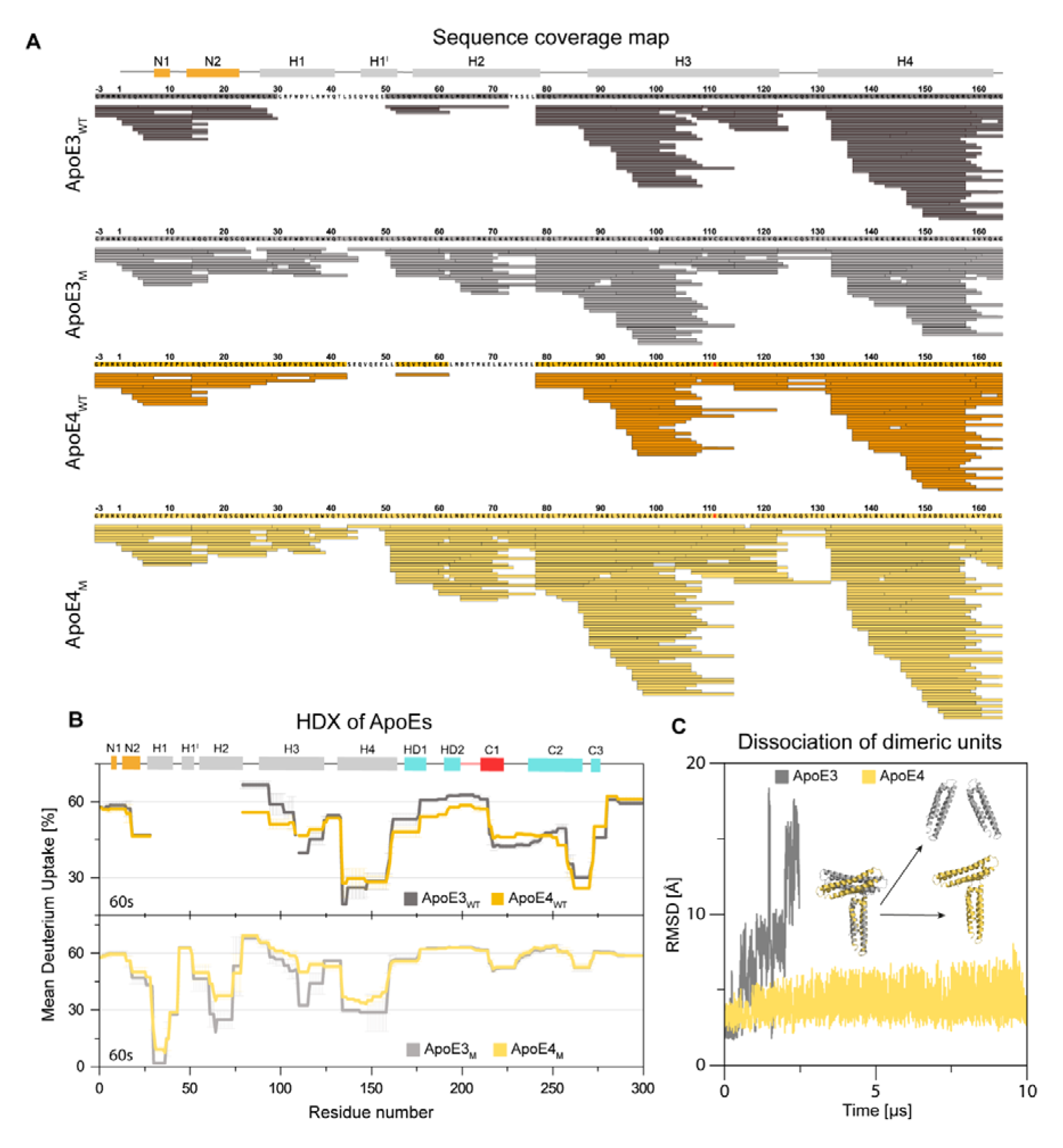
Comparison of structural properties of ApoE isoforms at different self-association states. **(A)** LC-MS/MS analysis of NTD of ApoE_WT_ and ApoE_M_ proteins. **(B)** Comparison of ApoE_WT_ (the upper panel) and ApoE_M_ (the lower panel) mean deuterium uptake after 60 sec. ApoE3 and ApoE4 isoforms are shown in grey and yellow, respectively. The data were determined in 3 time intervals (60s, 120s and 600s) and error bars represent standard error of mean (SEM). **(C)** Dissociation of ApoE3 (grey) and ApoE4 (yellow) dimeric units during MD simulations.

Similarly to the ApoE_WT_, the ApoE_M_ isoforms differ in helix H3 around C112 (residues 110-123), as well as neighboring helices H2 (residues 61-78) and H4 (residues 134- 158) (**Figure 4B**). However, a systematic change in solvation over time was observed in a different part of NTD (**Supplementary Figure 9**). Conversely, no difference in deuterium uptake was observed in CTD between ApoE3_M_ and ApoE4_M_. A significant increase in solvation of CTD of the ApoE_M_ compared to CTD of the ApoE_WT_ suggests that tetramers are not formed due to the loss of interactions in this region upon monomerization. Altogether, the results of these experiments provide strong evidence that the increased aggregation propensity of ApoE4 compared to the other isoforms is driven by the structural changes induced by the C112R substitution to the self-associating interface.

Using the adaptive sampling MD, we observed that the V-shaped dimeric unit of ApoE4 is more resilient to dissociation than the T-shaped dimeric unit of ApoE3 (**Figure 4C**). While the ApoE3 dimeric unit dissociated after ∼2.5 µs, the dissociation of the ApoE4 dimeric unit did not occur even after ∼10 µs. We also found significantly stronger binding energy in the dimeric unit of ApoE4 than in ApoE3, explaining its resilience to dissociation. These interactions were formed primarily by electrostatic contacts (**Supplementary Table 8** and **Supplementary Note 1**). Altogether, these results suggest that the difference in the binding angle and strength between the two interacting domains (chains A and B) could be related to their different aggregation propensity.

### Drug candidate tramiprosate and its metabolite SPA induce ApoE3-like conformational behavior in ApoE4

Drug candidate tramiprosate and the metabolite SPA have shown a positive effect in patients with AD and ApoE ε4/ε4 genotypes but not on patients with other ApoE genotypes [14]. We carried out *in vitro* aggregation experiments at 37°C and pH 7.4 with both ApoE3 and ApoE4 in the presence of tramiprosate and SPA. The signal from light scattering decreased in the presence of both SPA (**Figure 5A**) and tramiprosate (**Supplementary Figure 10)**, suggesting the reduction of ApoE4 aggregation. This finding is consistent with molecular dynamics simulations, in which the stable ApoE4 V-shaped dimeric unit dissociated in the presence of SPA after 2.5 µs, which is within the time scale of ApoE3 dissociation (**Supplementary Figure 11**). To understand such behavior, we analyzed the interactions at the self-association interface. We found that the binding energy between the two interacting domains dropped in the presence of SPA. SPA interacted with some of the charged residues involved in keeping the NTDs of the two chains closely together, i.e., R38, R145 and E45, E49, decreasing the stability of the ApoE4 V-shaped dimeric unit and subsequently leading to its dissociation (**Supplementary Table 8** and **Supplementary Note 2**).

**Figure 5.**
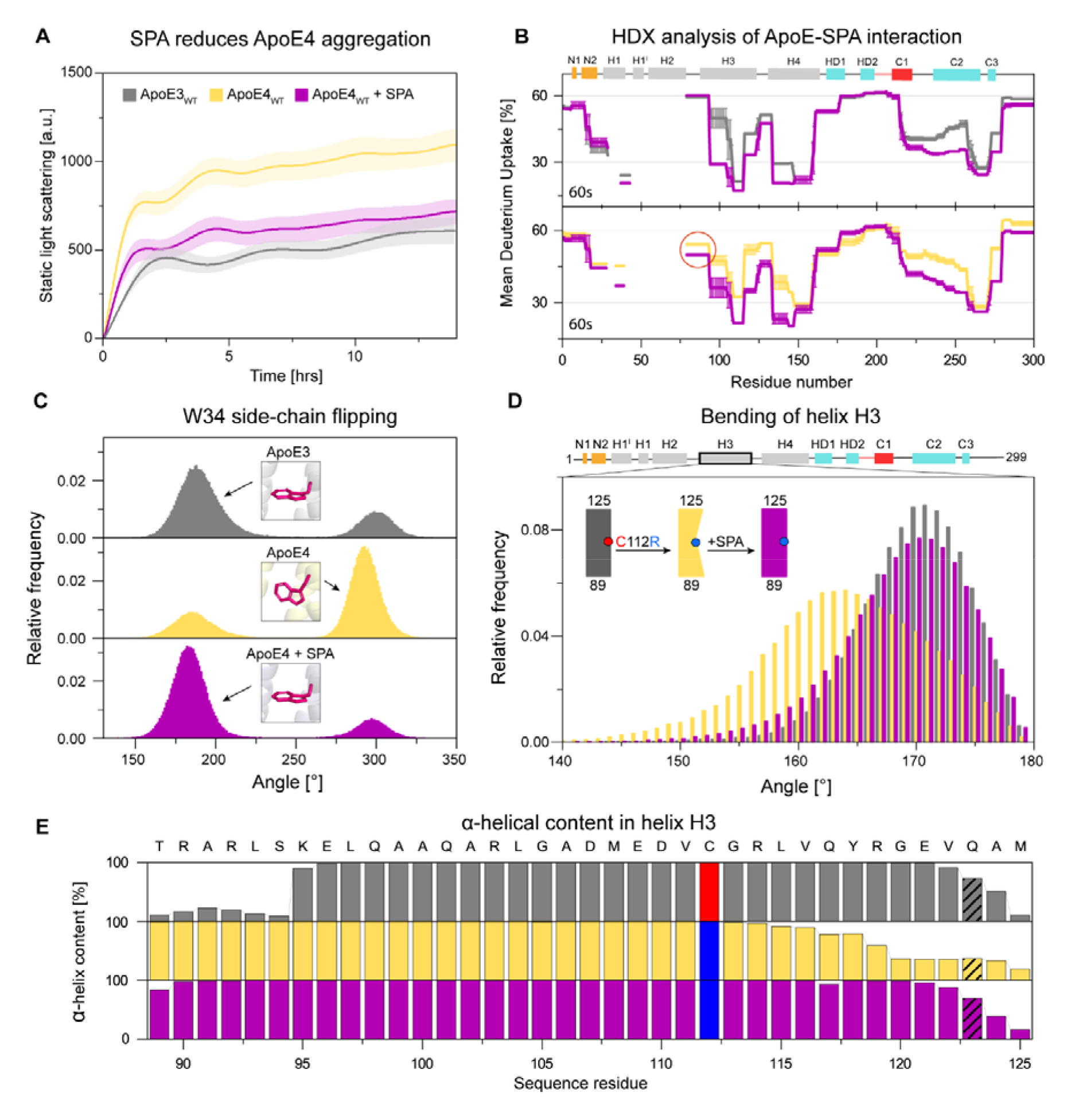
Modulation of ApoE4 behavior toward ApoE3-like upon its interaction with SPA. **(A)** Reduction of ApoE4 aggregation (yellow) by SPA (purple) to the level of ApoE3 aggregation (grey). The data were collected in Tris buffer at 37 °C and pH 7.4, representing averages of 3 replicates with standard deviation shown as error band. **(B)** Comparison of ApoE3 (grey) and ApoE4 (yellow) mean deuterium uptake after 60 sec in the presence of SPA (purple). The red circle highlights the interaction of ApoE4 with SPA in the region corresponding to the loop connecting helices H2 and H3. The data were determined in 3 time intervals (60s, 120s and 600s) and error bars represent standard error of mean (SEM). **(C)** Comparison of histogram distribution of the dihedral angle *χ*_1_ of the W34 in ApoE3_M_ (grey), ApoE4_M_ (yellow), and ApoE4_M_-SPA (purple). **(D)** Comparison of histogram distribution of the angle between the C_αα_ atoms of residues 95-106-117 of the helix in ApoE3_M_ (grey), ApoE4_M_ (yellow), and ApoE4_M_-SPA (purple). **(E)** Comparison of the α-helical content of helix H3 in ApoE3_M_ (grey), ApoE4_M_ (yellow), and ApoE4_M_-SPA (purple) calculated by adaptive simulations. The critical residues in positions 112 and 123 discussed in the text are labelled.

These findings are further supported by the HDX results (**Figure 5B** and **Supplementary Figures 12-14**) which indicate that the interaction of SPA with ApoE is non-specific. SPA suppressed solvation in both isoforms equally, mainly in helices H3 (residues 94-123), H4 (residues 134-146), and the region spanning from helices C1 to C2 (residues 219- 257) in CTD. Interestingly, SPA interacted with the loop connecting helices H2 and H3 in ApoE4_WT_ but not in ApoE3_WT_. In the case of ApoE_M_ isoforms with or without SPA, no changes in deuteration were observed in the CTD parts of the proteins (**Supplementary Figure 15**). The bimodal isotopic pattern observed in hydrogen exchange mass spectra of ApoE3_M_ free and ApoE4_M_ in the presence of SPA suggested that some peptides from the 15-109 protein segment underwent cooperative unfolding with EX1 kinetics signatures (**Supplementary Figure 16**). Such EX1 kinetics were previously described for ApoEs free monomolecular protein mutants [79]. In our case, we produced two peptide populations: (i) unfolded without H/D exchange, with the same isotopic envelope as the non-deuterated samples, and (ii) folded, with an isotopic envelope that showed ongoing H/D exchange. The results from EX1 kinetics deduced that upon interaction with SPA, ApoE4_M_ behaved similarly to free ApoE3_M_. All the other peptides exhibited EX2 kinetics.

In the next step, we complemented data from HDX-MS by performing adaptive sampling MD simulations of the full-length ApoE_M_ isoforms. We constructed Markov state models from these simulations using the RMSD of the protein Cα atoms for the ApoE3_M,_ ApoE4_M_, ApoE3_M_+SPA and ApoE4_M_+SPA systems **(Supplementary Figure 17)**. The distribution of the RMSD values varied extremely. This was primarily due to the high fluctuations of the CTD, while the NTD mainly remained stable (**Supplementary Note 3**). We, therefore, analyzed these MD simulations in more detail, with a focus on the NTD areas. Primarily, we compared two critical differences between the ApoE3 and ApoE4 structures: (i) the orientation of the W34 side-chain at the self-association interface and (ii) the C112R substitution and its “domino-like” effect on the helix H3 (residues 89-125) conformation and stability. To compare the “flipping” of the W34 side-chain, we calculated the dihedral angle *χ*_1_ in the ApoE3_M,_ ApoE4_M_ and ApoE4_M_+SPA systems (**Figure 5C**). We found that ApoE3_M_ displayed a majority of its population around *χ*_1_ = 180° while ApoE4_M_ was mostly around *χ*_1_ = 300°. Interestingly, SPA strongly shifted the orientation of the W34 side chain in ApoE4_M_ towards the *χ*_1_ = 180°, resembling free ApoE3_M_. The main effect of the C112R substitution on the conformation of the H3 helix was reflected in the bending of the helix (**Figure 5D**). The kink of this H3 helix was calculated as the angle between the C_αα_ atoms of residues 95, 106 and 117, where 180° was a non-bent helix. We found that the H3 helix was ∼7° straighter in ApoE3_M_ than in ApoE4_M_. Moreover, ApoE4_M_ showed a more spread and skewed distribution. As in the previous case, the presence of SPA induced the H3 helix in ApoE4_M_ to become more similar to that of ApoE3_M_.

Finally, we investigated whether the C112R substitution also results in a change in the stability of the H3 helix in terms of α-helical content (**Figure 5E**). The results showed partial unfolding at the N-terminus of the H3 helix (residues 89-94) in the case of ApoE3, while no change was observed in ApoE4. Conversely, we observed a significant decrease in α-helical content at the end of the H3 helix (residues 120-124) in ApoE4. This finding indicates a possible destabilization of Q123, leading to a loss of interaction between Q123 and W39/T42 in the ApoE4 V-shaped dimeric unit. The presence of SPA in the ApoE4 simulation stabilized this region which is reflected in the increased content of helical structures at the end of the H3 helix where it resembled its ApoE3 counterpart. Overall, SPA induces an ApoE3-like conformation upon ApoE4, making it less aggregation-prone.

### Treatment of ApoE **ε**4/**ε**4 cerebral organoids by tramiprosate increased the levels of AD-related proteins and cholesteryl esters

Lastly, we aimed to study the effects of tramiprosate in the biologically relevant system of cerebral organoids. To perform this experiment, we used a previously generated and characterized isogenic pair of induced pluripotent stem cell lines carrying the ApoE ε4/ε4 (sAD-E4) and ApoE ε3/ε3 (sAD-E3) genotypes [72]. These two cell lines were differentiated into cerebral organoids and, upon maturation, continuously treated with 100 µM tramiprosate for 50 days (**Supplementary Figure 18**). Specifically, the treatment with tramiprosate began on day 50 of the culturing process and continued until day 100, as samples were harvested. Two independent batches of organoids were generated from each cell line and used for all subsequent analyses to demonstrate reproducibility.

First, we verified that generated organoids showed a standard morphology upon differentiation. As shown in **Figure 6A**, organoids from the sAD-E3 cell line formed typical spheroids with smooth edges (visible in brightfield) with numerous structures visible on immunohistological sections (as shown by cell nuclei counterstained with Hoechst). Our immunohistochemical analysis identified regions positive for markers of neural progenitor cells (PAX6), immature and mature neurons (DCX, TUJ, MAP2), astrocytes (S100b, GFAP), microglia (IBA1), as well as the presence of ApoE. The expression of these markers thus confirmed the successful differentiation of cerebral organoids *in vitro*. Levels of these cell markers were later quantified in individual organoids by mass spectrometry protein assays.

**Figure 6.**
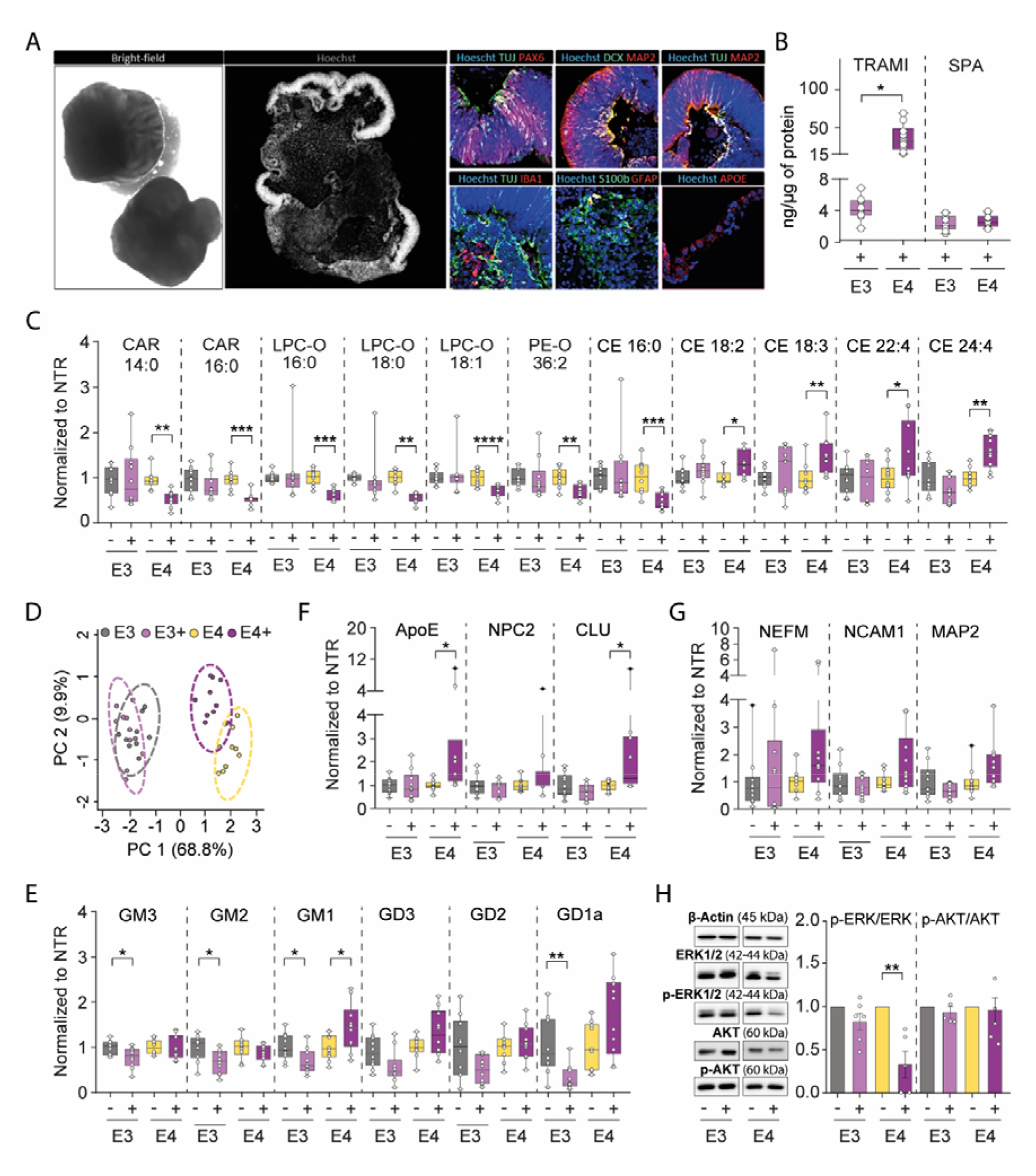
Cerebral organoids derived from ApoE3 ε3/ε3 and ApoE4 ε4/ε4 cerebral organoids show differences upon treatment with tramiprosate in metabolism, cholesterol storage and MAP kinase signaling pathway. (**A**) Morphology of cerebral organoids carrying ApoE3 ε3/ε3 and ApoE4 ε4/ε4 alleles. (**B**) Levels of tramiprosate (TRAMI) and 3- sulfopropanoic acid (SPA) in untreated (-) and treated (+) ApoE e3/e3 and e4/e4 organoids (N=10). (**C**) Levels of carnitines (CAR), ether-linked lysophospholipid (LPC-O) and phospholipid species (PE-O), and cholesteryl esters (CE) in untreated (-) and treated (+) ApoE e3/e3 and e4/e4 organoids (N=10). (**D)** Principal component analysis (PCA) of 64 dysregulated lipid species. (**E**) Levels of major mono-sialo-glycosphingolipid (GM1-3) and di-sialo-glycosphingolipid (GD1-3) species in untreated (-) and treated (+) ApoE e3/e3 and e4/e4 organoids (N=10). (**F** and **G**) Levels of proteins involved in lipid transport (ApoE, NPC2, CLU), and neuronal protein markers (NEFM, NCAM1, MAP2) in untreated (-) and treated (+) ApoE e3/e3 and e4/e4 organoids (N=10). (**H**) Western blot showing that ERK, but not AKT signaling pathway is downregulated upon treatment selectively in ApoE4 organoids. Targeted LC-MS-based proteomics and lipidomics analysis of cerebral organoids untreated (-) and treated (+) with tramiprosate (N=10). Biological outliers have been presented as black dots (●). Statistical analysis was performed using the Kruskal-Wallis test, comparing the cerebral organoids treated with tramiprosate to the untreated, *p≤0.05, **p≤0.01, ***p≤0.001, ****p≤0.0001 (N=10). A comprehensive list of abbreviations and nomenclature systems for lipids and proteins can be found in **Supplementary Table 10**.

Upon the verification of organoid maturity, we initiated continuous treatment with tramiprosate. After 50 days, no cytotoxicity or morphological changes were observed. Thus, both untreated and treated organoids were subjected to a comprehensive mass spectrometry characterization of metabolites, lipids and proteins. Metabolomic analysis revealed that tramiprosate was present in ApoE ε3/ε3 cerebral organoids in lower concentration than in ApoE ε4/ε4 (**Figure 6B**). The reason for this lower concentration (n=10, **p=0.01) of tramiprosate in ApoE ε3/ε3 organoids is unknown. We speculate that this difference could be related to uptake due to different biomembrane compositions rather than metabolic conversion. Importantly, cerebral organoids metabolized added tramiprosate into SPA in agreement with clinical observations [12].

Since ApoE plays a central role in lipid metabolism [80], we aimed to analyse the effect of tramiprosate on the level of lipids and gangliosides. We studied 310 lipid species belonging to 22 different lipid classes. The tramiprosate effect on the level of total lipid classes is shown in **Supplementary Figure 19**. Tramiprosate did not show a significant effect in ApoE ε3/ε3 organoids while it led to a downregulation of carnitines (CAR), lysophosphatidylcholines (LPC-O) and oxidized phosphatidylethanols (PE-O), as well as an upregulation of ceramide (CER) in ApoE ε4/ε4 (**Figure 6C** and **Supplementary Figure 19**). Further lipid species level investigation revealed dysregulation of 64 lipid species (**Supplementary Table 9**) following the tramiprosate treatment. Principal component analysis (PCA) of 64 lipid species shows greater separation of treated *versus* non-treated ApoE ε4/ε4 organoids when compared to ApoE ε3/ε3 treated *versus* non-treated organoids (**Figure 6D**). Several species of cholesteryl ester with polyunsaturated fatty acids (CE18:2, CE18:3, CE22:4 and CE24:4) were upregulated in ApoE ε4/ε4, while CE 16:0 was downregulated. Cholesteryl ester significantly improves the transport of cholesterol, as it can be packed into lipoprotein particles [81]. Long-chain carnitine species (CAR12:0, CAR14:0, CAR15:0, CAR16:0, CAR18:0 and CAR18:1) were downregulated in ApoE ε4/ε4 upon the tramiprosate treatment. This could be due to the increased utilization of carnitines by beta-oxidation [82]. We also saw an ApoE4-specific downregulation of several species of ether lysophospholipid (LPC-O) and ether phospholipids (PE-O and PC-O). Ether phospholipids are known to be pro-inflammatory, previously reported to be upregulated in AD patients. A clinical study revealed that tramiprosate reduces the level of pro-inflammatory cytokines, which corresponds with our observation of the downregulation of ether-linked phospholipids [83,84].

The lipidomic analysis further quantified six gangliosides (**Figure 6E**). A significant decline in gangliosides (GM3, GM2, GM1, GD1a; n=10, *p<0.05) was observed in the ApoE ε3/ε3. On the contrary, an increase in levels of gangliosides was observed in the ApoE ε4/ε4 upon treatment with tramiprosate. These trends mimic the changes in ceramide levels (especially Cer36:1), a precursor of these major glycosphingolipids.

We then performed the targeted proteomic mass spectrometry analysis of 32 proteins (**Supplementary Table 4**), which revealed that the treatment with tramiprosate slightly increased the level of AD-related lipid transport, Aβ amyloid binding proteins (ApoE, NPC2 and CLU) and neuronal morphogenesis proteins (NEFM, NCAM1 and MAP2) in ApoE ε4/ε4 cerebral organoids (**Figures 6F** and **6G**). Conversely, tramiprosate did not show an effect on these proteins in ApoE ε3/ε3 cerebral organoids. This suggests that signaling via ApoE ε4/ε4 stimulates neurogenesis [85].

Moreover, tramiprosate showed a similar effect on the expression of PPIA (n=10, *p=0.03) and HNRNPAB protein (n=10, **p=0.002) that was increased only in ApoE ε4/ε4 cerebral organoids (**Supplementary Figure 16**). The results on HNRNPAB might indicate a positive effect of tramiprosate on cholinergic neurons, as it was previously reported that the destruction of cholinergic neurons causes a decrease of HNRNPAB levels as it occurs in AD [86]. It has been previously shown that ApoE3 and ApoE4 differentially activate multiple neuronal pathways and regulate synaptogenesis via MAP kinase signaling [85]. We opted to determine the activation of ERK1/2 and AKT signaling molecules upon tramiprosate treatment using the Western blot (**Figure 6H**). We found that the phosphorylation of key signal transduction protein of MAP kinase signaling (ERK1/2) was significantly downregulated only in cerebral organoids derived from ApoE ε4/ε4, but not ApoE ε3/ε3 (n=3, **p=0.001. No significant differences in phosphorylation of AKT upon treatment with tramiprosate were detected. Since it was previously shown that ApoE ε4/ε4 stimulates MAP-kinase signaling to a greater extent than ApoE ε3/ε3 [85], our data suggest that tramiprosate has the potential to reduce P-ERK1/2 levels selectively in ApoE ε4/ε4, mimicking those of ApoE ε3/ε3.

In summary, the data obtained from the treatment of cerebral organoids by tramiprosate revealed that: (i) it was endogenously metabolized into SPA in cerebral organoids, (ii) it increased the concentration of cholesteryl ester and decreases the level of carnitines specifically in ApoE ε4/ε4 organoids, (iii) it increased the level of AD-related proteins involved in lipid transport and stimulated neurogenesis, and (iv) ApoE ε4/ε4 organoids treated with tramiprosate showed a specific decrease of P-ERK1/2 signaling levels.

## Discussion

ApoE plays a crucial role in AD pathogenesis because it affects multiple essential pathways, including the Aβ-amyloid and lipid metabolism [87]. The “ApoE Cascade Hypothesis” proposed by Bu and co-workers [88] states that disruption of ApoE-mediated cellular lipid homeostasis by misfolding and oligomerization of ApoE initiates a pathogenic cascade that contributes to AD-related cellular dysfunction. It is widely accepted that the isoform and lipidation status of ApoE influences the ApoE aggregation-driven pathology [89–91]. Liao and colleagues suggested that a primary mechanism for ApoE-mediated plaque formation is a result of lipid-free ApoE aggregation, as targeting ApoE aggregates by therapeutic antibodies reduced Aβ pathology [92]. Although the lipid-free ApoE species should be relatively rare [93–96], it has been demonstrated that the presence of lipid-free ApoE increases amyloid deposition [97,98].

Here, we propose that the ApoE T-shaped/V-shaped dimer represents a basic unit of lipid-free ApoE aggregates. We based our hypothesis on the packing pattern observed in the crystal lattices of ApoE proteins. Interestingly, while all three ApoE isoforms share the same self-association interface, the pathological ApoE4 isoform differs from the ApoE2 and ApoE3 isoforms by the angle between the two interacting NTDs. We demonstrate that this angular difference is a consequence of a “domino-like effect” of the C112R substitution, starting with the loss of the R61-E109 interaction, leading to destabilization of the H3 helix and re-orientation of Q123. This structural interpretation is further supported by computer simulations, which showed destabilization and bending of the H3 helix in ApoE4. Altered intradomain interaction and H3 bending as a consequence of C112R substitution and R61 rearrangement in ApoE4 have been previously reported [99–101]. These conformational changes make ApoE4 more susceptible to chemical and thermal denaturation than ApoE3 and ApoE2 [102,103]. Furthermore, our experimental data confirmed an increased tendency of ApoE4 to aggregate, compared to ApoE3. Computer simulations of T-shaped and V-shaped dimeric unit dissociation revealed that the ApoE4 unit is more resilient to dissociation than the ApoE3 unit. This is consistent with previous observations that ApoE4, as opposed to ApoE3, can form filamentous aggregates and that this process involves NTDs [38,104]. Therefore, V-shaped dimeric unit model provides a rationale for the increased aggregation propensity of ApoE4.

Only 4 of the 21 ApoE structures experimentally determined do not fit our hypothesis. The two ApoE3 structures (PDB IDs: 1OR2 and 1OR3) have a V-shaped packing, and on the other hand, the two ApoE4 structures (PDB IDs: 1LE4 and 1B68) have a T-shaped packing. A possible explanation for the second case is that these ApoE4 proteins were purified in the presence of heparin, whose binding site was identified by the authors to overlap with the self-association interface partially [105]. In the case of ApoE3 structures, we showed that they represent a kind of intermediate state between V-shaped and T-shaped packing caused by the different conformation of the loop connecting the H2 and H3 helices. The high flexibility of this loop leads to the bending of the H2 and H3 helices. This bending has already been reported in these ApoE3 structures [106]. Noteworthy that the structure (PDB ID: 1OR2) is an orthorhombic crystal form whose cell dimensions do not match any other ApoE structure (**Supplementary Table 7**). The structure (PDB ID: 1OR3) is the only trigonal crystal form of ApoE3 with almost the exact cell dimensions and packing as our trigonal crystal form of ApoE4 (PDB ID: 8AX8). Interestingly, conformational alterations in these unique structures led to V-shaped packing, considered “pathological” in our proposal. On the other hand, we found that SPA interacts with ApoE4 but not with ApoE3 in this region.

Enzymatic cleavage of ApoE revealed that the region corresponding to the self-association interface could not be detected by mass spectrometry, which is consistent with previous studies [107,108]. It is one of seven highly conserved regions of ApoE and the only one to which no biological function could previously be assigned [109]. To explore whether this self-association interface represents a biologically relevant interface (**Supplementary Figure 21** and **Note 4**), we created a structural model of a full-length ApoE3 dimer by superposition of ApoE3 T-shaped dimeric unit with the ApoE3_M_. The model contained a visible clash between residues 1-23 and residues 204-223 near the self-association interface. Interestingly, regions corresponding to residues 12-20 and 204-210 have been previously identified as a major source of the distinction between tetrameric ApoE3 and ApoE4 isoforms [79]. Based on this model, the region corresponding to residues 204-223 (helix C1) was the only part of the CTD of chain A that was in direct contact with chain B. Interestingly, this model also suggests a possible interaction between helix C1 from chain A and the end of helix H3 containing Q123 from chain B. This region was more disordered in ApoE4 [79] and may also contribute to the different behavior of ApoE4 and ApoE3 isoforms.

In contrast, we showed that the position of the N1 and N2 helices in the NMR structure of ApoE3_M_ is induced by the five “monomerizing” point mutations at the end of the CTD [31]. This newly acquired conformation prevents the self-association of ApoE due to the clash with helix C1 (residues 204-223) of the second ApoE molecule. Aggregation of ApoE_M_ upon removal of the helices N1 and N2 (residues 1-23) despite five “monomerizing” mutations validates this proposal. The displacement of helices N1 and N2 explains these mutations’ “monomerizing” effect in the ApoE_M_. Further suppression of aggregation by introducing four additional spatially distant mutations localized at the self-association interface confirmed the involvement of this interface in ApoE aggregation.

Structural analysis can accelerate rational drug design by providing high-resolution molecular targets [110]. We have investigated the interaction between ApoE isoforms and tramiprosate/SPA at the molecular and cellular levels. At the molecular level, we showed that SPA acts via non-specific interactions (**Figure 5B**) and displays an anti-aggregation activity in ApoE4 (**Figure 5A**). Interestingly, SPA modulates the structural features of ApoE4, e.g., the conformation of helix H3 and the orientation of W34 towards resembling ApoE3. It has been previously shown that small-molecule structure correctors can modify the aberrant conformation of ApoE4 and abolish its detrimental effects in cultured neurons [111]. Petros and co-workers identified a small molecule which binds to the cavity formed by the rotation of W34 in ApoE4, stabilizes its structure and suppresses neuroinflammation in the cell culture [29]. Interestingly, the crystal structure of the complex of ApoE4 with corrector molecule (PDB ID 6NCN) showed that the bound ligand interacts with W34 and changes its orientation to that observed in ApoE3 structures. A closer inspection of this structure revealed that the re-orientation of W34 caused ApoE4 crystallization through the T-shaped dimeric unit, which supports our observations.

Oral drug candidate ALZ-801, a prodrug of tramiprosate, recently entered phase 3 clinical trials to treat ApoE ε4/ε4 patients with mild or moderate AD [112]. The proposed mode of action for tramiprosate and SPA is a direct interaction with Aβ-amyloids and suppressing their oligomerization [11]. Here we demonstrate that tramiprosate and its metabolite SPA also modify the aggregation propensity of ApoE4. HDX-MS experiments and computer simulations carried out in the presence of SPA revealed non-specific interactions of these small polar molecules with charged residues at the surface of ApoE4, inducing ApoE3-like conformational behavior. Moreover, surface charges are partially neutralized by these non-specific interactions, which impact ApoE oligomerization and aggregation.

At the cellular level, the data collected with cerebral organoids demonstrate that tramiprosate and its metabolite SPA influence lipid transport/Aβ binding proteins and cholesterol homeostasis. Specifically, the concentration of lipid-storage molecules cholesteryl esters was increased in ApoE ε4/ε4 cerebral organoids, while it remained unchanged in ApoE ε3/ε3 organoids treated by tramiprosate. Interestingly, a recent study showed that facilitating cholesterol transport increases myelination and improves cognitive function in ApoE4 mice [113]. In addition, cholesteryl esters were recently identified as an upstream regulator of pTau proteostasis and Aβ-amyloid secretion [114]. Additionally, we also found that tramiprosate specifically downmodulates MAP kinase signaling (via downregulation of P-ERK1/2) only in ApoE ε4/ε4 but not in ApoE ε3/ε3 cerebral organoids (n = 3, P = 0.001). Combining previous clinical observations [14] with the data presented in this study implies that the drug candidate tramiprosate and its metabolite SPA can suppress ApoE4 aggregation besides interacting with Aβ-amyloids in ApoE ε4/ε4 patients.

## Conclusions

In summary, the proposed new self-association interface at play in modulating ApoE dysfunction as induced by the “domino-like” effects of C112R mutation supports the “ApoE Cascade Hypothesis” [88]. Understanding the structural basis of ApoE aggregation and constructing truncated mutants with accelerated aggregation propensity will stimulate the development of specific and potent corrector molecules. Corrector molecules could target ApoE4 produced in brains and the periphery (without a need to cross a brain-blood-barrier), based on the recent report by Liu and co-workers [115]. These corrector molecules can potentially treat AD, and other ApoE4-related disorders, like Parkinson’s disease and Lewy body dementia [116], and modulate the course of ageing [117].

### Abbreviations

Aβ: amyloid beta
AD: Alzheimer’s disease
AKT: protein kinase B
ApoE: apolipoprotein E
CAR: carnitines
CE: cholesteryl esters
CLU: clusterin
CTD: C-terminal domain
DCX: doublecortin
ERK1/2: extracellular signal-regulated kinase
GFAP: glial fibrillary acidic protein
GD: di-sialo-glycosphingolipid
GM: mono-sialo-glycosphingolipids
HD: hinge domain
HDX-MS: hydrogen-deuterium mass spectrometry
HNRNPAB: heterogenous nuclear ribonucleoprotein A/B
IBA1: ionized calcium-binding adapter molecule 1
LPC-O: alkyl ether-linked lysophosphatidylcholines
MAP2: microtubule-associated protein 2
MD: molecular dynamics
NCAM1: neural cell adhesion molecule 1
NEFM: neurofilament medium polypeptide
NMR: nuclear magnetic resonance
NPC2: NPC intracellular cholesterol transporter 2
NTD: N-terminal domain
MAP2: microtubule-associated protein 2
PAX6: paired box protein Pax-6
PCA: principal component analysis
PC-O: alkyl ether-linked phosphatidylcholines
PE-O: alkyl ether-linked phosphatidylethanolamines
PPIA: alkyl ether-linked phosphatidylethanolamines
S100b: S100 calcium-binding protein B
SPA: 3-sulfopropanoic acid
SRM: selected reaction monitoring
TEM: transmission electron microscopy
TUJ: tubulin beta-3 chain.

## Declarations

### Ethical Approval and Consent to participate

Not applicable.

### Consent for publication

All authors have approved of the consents of this manuscript and provided consent for publication.

### Availability of supporting data

The online version contains supplementary material available at…

## Authors’ contributions

Michal Nemergut – manuscript writing, data interpretation, protein preparation, structural analysis, biophysical analysis, mutagenesis; Sergio Marques – computer simulations; Lukas Uhrik – HDX-MS analysis; Tereza Vanova – experimental design and culture of cerebral organoids, data analysis, manuscript writing; Marketa Nezvedova – proteomic analysis; Darshak Chandulal Gadara – lipidomics analysis; Durga Jha – lipidomics analysis; Jan Tulis – protein preparation; Veronika Blechova – mutagenesis and DNA cloning; Joan Iglesias-Planas – data analysis; Antonin Kunka – biophysical measurements; Anthony Legrand – biophysical measurements; Hana Hornakova – electron microscopy; Veronika Pospisilova - confocal microscopy; Jiri Sedmik - culture of cerebral organoids; Jan Raska - western blot analysis; Zbynek Prokop – data interpretation; Jiri Damborsky – concept, data interpretation, coordination; Dasa Bohaciakova – cerebral organoids experiments, data interpretation; Zdenek Spacil – omics analysis, data interpretation; Lenka Hernychova – HDX-MS analysis, data interpretation; David Bednar – computer modelling, data interpretation; Martin Marek – crystallization and structure determinations, data interpretation, supervision of research. All authors read and approved the final manuscript.

## Acknowledgements

The authors are thankful to Dr Stanislav Mazurienko (Loschmidt Laboratories, Masaryk University, Brno) for his help with data analysis and to members of the Swiss Light Source (SLS) synchrotron for using their beamline facilities and help during data collection. We acknowledge CF Biomolecular Interactions and Crystallography of CIISB, Instruct-CZ Centre, supported by MEYS CR (LM2023042) and European Regional Development Fund-Project “UP CIISB” (No. CZ.02.1.01/0.0/0.0/18_046/0015974).

## Authors’ information

Not applicable.

## References

1. Tomaskova H, Kuhnova J, Cimler R, Dolezal O, Kuca K. Prediction of population with Alzheimer’s disease in the European Union using a system dynamics model. Neuropsychiatr Dis Treat. 2016;12:1589–98.

2. 2022 Alzheimer’s disease facts and figures. Alzheimers Dement J Alzheimers Assoc. 2022;18:700– 89.

3. Gustavsson A, Norton N, Fast T, Frölich L, Georges J, Holzapfel D, et al. Global estimates on the number of persons across the Alzheimer’s disease continuum. Alzheimers Dement [Internet]. [cited 2022 Jun 9];n/a. Available from: https://onlinelibrary.wiley.com/doi/abs/10.1002/alz.12694

4. Cummings J, Bauzon J, Lee G. Who funds Alzheimer’s disease drug development? Alzheimers Dement Transl Res Clin Interv. 2021;7:e12185.

5. Huang L-K, Chao S-P, Hu C-J. Clinical trials of new drugs for Alzheimer disease. J Biomed Sci. 2020;27:18.

6. Knopman DS, Jones DT, Greicius MD. Failure to demonstrate efficacy of aducanumab: An analysis of the EMERGE and ENGAGE trials as reported by Biogen, December 2019. Alzheimers Dement. 2021;17:696–701.

7. Cummings J, Lee G, Nahed P, Kambar MEZN, Zhong K, Fonseca J, et al. Alzheimer’s disease drug development pipeline: 2022. Alzheimers Dement Transl Res Clin Interv. 2022;8:e12295.

8. Huang Y, Mucke L. Alzheimer mechanisms and therapeutic strategies. Cell. 2012;148:1204–22.

9. Hey JA, Yu JY, Versavel M, Abushakra S, Kocis P, Power A, et al. Clinical Pharmacokinetics and Safety of ALZ-801, a Novel Prodrug of Tramiprosate in Development for the Treatment of Alzheimer’s Disease. Clin Pharmacokinet. 2018;57:315–33.

10. Gervais F, Paquette J, Morissette C, Krzywkowski P, Yu M, Azzi M, et al. Targeting soluble Abeta peptide with Tramiprosate for the treatment of brain amyloidosis. Neurobiol Aging. 2007;28:537–47.

11. Kocis P, Tolar M, Yu J, Sinko W, Ray S, Blennow K, et al. Elucidating the Aβ42 Anti-Aggregation Mechanism of Action of Tramiprosate in Alzheimer’s Disease: Integrating Molecular Analytical Methods, Pharmacokinetic and Clinical Data. CNS Drugs. 2017;31:495–509.

12. Hey JA, Kocis P, Hort J, Abushakra S, Power A, Vyhnálek M, et al. Discovery and Identification of an Endogenous Metabolite of Tramiprosate and Its Prodrug ALZ-801 that Inhibits Beta Amyloid Oligomer Formation in the Human Brain. CNS Drugs. 2018;32:849–61.

13. Abushakra S, Porsteinsson A, Vellas B, Cummings J, Gauthier S, Hey JA, et al. Clinical Benefits of Tramiprosate in Alzheimer’s Disease Are Associated with Higher Number of APOE4 Alleles: The “APOE4 Gene-Dose Effect.” J Prev Alzheimers Dis. 2016;3:219–28.

14. Abushakra S, Porsteinsson A, Scheltens P, Sadowsky C, Vellas B, Cummings J, et al. Clinical Effects of Tramiprosate in APOE4/4 Homozygous Patients with Mild Alzheimer’s Disease Suggest Disease Modification Potential. J Prev Alzheimers Dis. 2017;4:149–56.

15. Abushakra S, Porsteinsson AP, Sabbagh M, Bracoud L, Schaerer J, Power A, et al. APOE ε4/ε4 homozygotes with early Alzheimer’s disease show accelerated hippocampal atrophy and cortical thinning that correlates with cognitive decline. Alzheimers Dement N Y N. 2020;6:e12117.

16. Corder EH, Saunders AM, Strittmatter WJ, Schmechel DE, Gaskell PC, Small GW, et al. Gene dose of apolipoprotein E type 4 allele and the risk of Alzheimer’s disease in late onset families. Science. 1993;261:921–3.

17. Sadigh-Eteghad S, Talebi M, Farhoudi M. Association of apolipoprotein E epsilon 4 allele with sporadic late onset Alzheimer’s disease. A meta-analysis. Neurosci Riyadh Saudi Arab. 2012;17:321– 6.

17. Korologou-Linden R, Bhatta L, Brumpton BM, Howe LD, Millard LAC, Kolaric K, et al. The causes and consequences of Alzheimer’s disease: phenome-wide evidence from Mendelian randomization. Nat Commun. Nature Publishing Group; 2022;13:4726.

19. Mahley RW, Weisgraber KH, Huang Y. Apolipoprotein E: structure determines function, from atherosclerosis to Alzheimer’s disease to AIDS. J Lipid Res. 2009;50:S183–8.

20. Holtzman DM, Bales KR, Tenkova T, Fagan AM, Parsadanian M, Sartorius LJ, et al. Apolipoprotein E isoform-dependent amyloid deposition and neuritic degeneration in a mouse model of Alzheimer’s disease. Proc Natl Acad Sci U S A. 2000;97:2892–7.

21. Verghese PB, Castellano JM, Garai K, Wang Y, Jiang H, Shah A, et al. ApoE influences amyloid-β (Aβ) clearance despite minimal apoE/Aβ association in physiological conditions. Proc Natl Acad Sci U S A. 2013;110:E1807–16.

22. Strittmatter WJ, Roses AD. Apolipoprotein E and Alzheimer’s disease. Annu Rev Neurosci. 1996;19:53–77.

23. Weisgraber KH, Rall SC, Mahley RW. Human E apoprotein heterogeneity. Cysteine-arginine interchanges in the amino acid sequence of the apo-E isoforms. J Biol Chem. 1981;256:9077–83.

24. Safieh M, Korczyn AD, Michaelson DM. ApoE4: an emerging therapeutic target for Alzheimer’s disease. BMC Med. 2019;17:64.

24. Yang A, Kantor B, Chiba-Falek O. APOE: The New Frontier in the Development of a Therapeutic Target towards Precision Medicine in Late-Onset Alzheimer’s. Int J Mol Sci. Multidisciplinary Digital Publishing Institute; 2021;22:1244.

26. Kim J, Eltorai AEM, Jiang H, Liao F, Verghese PB, Kim J, et al. Anti-apoE immunotherapy inhibits amyloid accumulation in a transgenic mouse model of Aβ amyloidosis. J Exp Med. 2012;209:2149–56.

27. Huynh T-PV, Liao F, Francis CM, Robinson GO, Serrano JR, Jiang H, et al. Age-Dependent Effects of apoE Reduction Using Antisense Oligonucleotides in a Model of β-amyloidosis. Neuron. 2017;96:1013–1023.e4.

28. Chen H-K, Liu Z, Meyer-Franke A, Brodbeck J, Miranda RD, McGuire JG, et al. Small Molecule Structure Correctors Abolish Detrimental Effects of Apolipoprotein E4 in Cultured Neurons. J Biol Chem. 2012;287:5253–66.

29. Petros AM, Korepanova A, Jakob CG, Qiu W, Panchal SC, Wang J, et al. Fragment-Based Discovery of an Apolipoprotein E4 (apoE4) Stabilizer. J Med Chem. American Chemical Society; 2019;62:4120– 30.

30. Garai K, Frieden C. The association−dissociation behavior of the ApoE proteins: kinetic and equilibrium studies. Biochemistry. 2010;49:9533–41.

31. Zhang Y, Vasudevan S, Sojitrawala R, Zhao W, Cui C, Xu C, et al. A monomeric, biologically active, full-length human apolipoprotein E. Biochemistry. 2007;46:10722–32.

32. Chen J, Li Q, Wang J. Topology of human apolipoprotein E3 uniquely regulates its diverse biological functions. Proc Natl Acad Sci. Proceedings of the National Academy of Sciences; 2011;108:14813–8.

33. Perugini MA, Schuck P, Howlett GJ. Self-association of human apolipoprotein E3 and E4 in the presence and absence of phospholipid. J Biol Chem. 2000;275:36758–65.

34. Chou C-Y, Lin Y-L, Huang Y-C, Sheu S-Y, Lin T-H, Tsay H-J, et al. Structural Variation in Human Apolipoprotein E3 and E4: Secondary Structure, Tertiary Structure, and Size Distribution. Biophys J. 2005;88:455–66.

35. Hubin E, Verghese PB, van Nuland N, Broersen K. Apolipoprotein E associated with reconstituted high-density lipoprotein-like particles is protected from aggregation. FEBS Lett. 2019;593:1144–53.

36. Namba Y, Tomonaga M, Kawasaki H, Otomo E, Ikeda K. Apolipoprotein E immunoreactivity in cerebral amyloid deposits and neurofibrillary tangles in Alzheimer’s disease and kuru plaque amyloid in Creutzfeldt-Jakob disease. Brain Res. 1991;541:163–6.

37. Gal J, Katsumata Y, Zhu H, Srinivasan S, Chen J, Johnson LA, et al. Apolipoprotein E Proteinopathy Is a Major Dementia-Associated Pathologic Biomarker in Individuals with or without the APOE Epsilon 4 Allele. Am J Pathol. Elsevier; 2022;192:564–78.

38. Hatters DM, Zhong N, Rutenber E, Weisgraber KH. Amino-terminal domain stability mediates apolipoprotein E aggregation into neurotoxic fibrils. J Mol Biol. 2006;361:932–44.

39. Sarkar G, Sommer SS. The “megaprimer” method of site-directed mutagenesis. BioTechniques. 1990;8:404–7.

40. Kabsch W. XDS. Acta Crystallogr D Biol Crystallogr. International Union of Crystallography; 2010;66:125–32.

41. Evans PR, Murshudov GN. How good are my data and what is the resolution? Acta Crystallogr D Biol Crystallogr. International Union of Crystallography; 2013;69:1204–14.

42. McCoy AJ, Grosse-Kunstleve RW, Adams PD, Winn MD, Storoni LC, Read RJ. Phaser crystallographic software. J Appl Crystallogr. International Union of Crystallography; 2007;40:658–74.

43. Adams P, Afonine P, Bunkóczi G, Chen V, Echols N, Headd J, et al. The Phenix Software for Automated Determination of Macromolecular Structures. Methods San Diego Calif. 2011;55:94–106.

44. Emsley P, Lohkamp B, Scott WG, Cowtan K. Features and development of Coot. Acta Crystallogr D Biol Crystallogr. International Union of Crystallography; 2010;66:486–501.

45. Kavan D, Man P. MSTools—Web based application for visualization and presentation of HXMS data. Int J Mass Spectrom. 2011;302:53–8.

46. Perez-Riverol Y, Csordas A, Bai J, Bernal-Llinares M, Hewapathirana S, Kundu DJ, et al. The PRIDE database and related tools and resources in 2019: improving support for quantification data. Nucleic Acids Res. 2019;47:D442–50.

47. Schuck P. Size-distribution analysis of macromolecules by sedimentation velocity ultracentrifugation and lamm equation modeling. Biophys J. 2000;78:1606–19.

48. Brautigam CA. Calculations and Publication-Quality Illustrations for Analytical Ultracentrifugation Data. Methods Enzymol. 2015;562:109–33.

48. 49. Hanwell MD, Curtis DE, Lonie DC, Vandermeersch T, Zurek E, Hutchison GR. Avogadro: an advanced semantic chemical editor, visualization, and analysis platform. J Cheminformatics. 2012;4:17.

50. Rappe AK, Casewit CJ, Colwell KS, Goddard WA, Skiff WM. UFF, a full periodic table force field for molecular mechanics and molecular dynamics simulations. J Am Chem Soc. 1992;114:10024–35.

51. Frisch MJ, Trucks GW, Schlegel HB, Scuseria GE, Robb MA, Cheeseman JR, et al. Gaussian 09, Revision E.01. Wallingford, CT: Gaussian, Inc.; 2009.

52. Case DA, Babin V, Berryman JT, Betz RM, Cai Q, Cerutti S, et al. AMBER 14. San Francisco: University of California; 2014.

53. Zoete V, Cuendet MA, Grosdidier A, Michielin O. SwissParam: a fast force field generation tool for small organic molecules. J Comput Chem. 2011;32:2359–68.

54. Rose PW, Bi C, Bluhm WF, Christie CH, Dimitropoulos D, Dutta S, et al. The RCSB Protein Data Bank: new resources for research and education. Nucleic Acids Res. 2013;41:D475–82.

55. Waterhouse A, Bertoni M, Bienert S, Studer G, Tauriello G, Gumienny R, et al. SWISS-MODEL: Homology modelling of protein structures and complexes. Nucleic Acids Res. 2018;46:W296–303.

56. Doerr S, Harvey MJ, Noé F, De Fabritiis G. HTMD: High-Throughput Molecular Dynamics for Molecular Discovery. J Chem Theory Comput [Internet]. 2016 [cited 2016 Aug 30];12:1845–52. Available from: http://pubs.acs.org/doi/abs/10.1021/acs.jctc.6b00049

57. Bas DC, Rogers DM, Jensen JH. Very fast prediction and rationalization of pKa values for protein– ligand complexes. Proteins Struct Funct Bioinforma. 2008;73:765–83.

58. Jorgensen WL, Chandrasekhar J, Madura JD, Impey RW, Klein ML. Comparison of simple potential functions for simulating liquid water. J Chem Phys. 1983;79:926–35.

59. Huang J, Rauscher S, Nawrocki G, Ran T, Feig M, de Groot BL, et al. CHARMM36m: an improved force field for folded and intrinsically disordered proteins. Nat Methods. 2017;14:71–3.

60. Frieden C, Wang H, Ho CMW. A mechanism for lipid binding to apoE and the role of intrinsically disordered regions coupled to domain-domain interactions. Proc Natl Acad Sci U S A. 2017;114:6292– 7.

61. Feenstra K. Anton, Hess Berk, Berendsen Herman J. C. Improving efficiency of large time-scale molecular dynamics simulations of hydrogen-rich systems. J Comput Chem. 1999;20:786–98.

62. Harvey MJ, De Fabritiis G. An Implementation of the Smooth Particle Mesh Ewald Method on GPU Hardware. J Chem Theory Comput. 2009;5:2371–7.

63. Harvey MJ, Giupponi G, Fabritiis GD. ACEMD: Accelerating Biomolecular Dynamics in the Microsecond Time Scale. J Chem Theory Comput. 2009;5:1632–9.

64. Hopkins CW, Le Grand S, Walker RC, Roitberg AE. Long-Time-Step Molecular Dynamics through Hydrogen Mass Repartitioning. J Chem Theory Comput. 2015;11:1864–74.

65. Naritomi Y, Fuchigami S. Slow dynamics in protein fluctuations revealed by time-structure based independent component analysis: the case of domain motions. J Chem Phys. 2011;134:065101.

66. Doerr S, Harvey MJ, Noé F, De Fabritiis G. HTMD: High-Throughput Molecular Dynamics for Molecular Discovery. J Chem Theory Comput. 2016;12:1845–52.

66. Swails J. ParmEd [Internet]. 2010 [cited 2018 Mar 8]. Available from: https://github.com/ParmEd/ParmEd

68. Roe DR, Cheatham TE. PTRAJ and CPPTRAJ: Software for Processing and Analysis of Molecular Dynamics Trajectory Data. J Chem Theory Comput. 2013;9:3084–95.

69. Abraham MJ, Murtola T, Schulz R, Páll S, Smith JC, Hess B, et al. GROMACS: High performance molecular simulations through multi-level parallelism from laptops to supercomputers. SoftwareX. 2015;1–2:19–25.

70. Kabsch W, Sander C. Dictionary of protein secondary structure: pattern recognition of hydrogen-bonded and geometrical features. Biopolymers. 1983;22:2577–637.

71. Aqvist J, Medina C, Samuelsson JE. A new method for predicting binding affinity in computer-aided drug design. Protein Eng. 1994;7:385–91.

72. Lin Y-T, Seo J, Gao F, Feldman HM, Wen H-L, Penney J, et al. APOE4 Causes Widespread Molecular and Cellular Alterations Associated with Alzheimer’s Disease Phenotypes in Human iPSC-Derived Brain Cell Types. Neuron. 2018;98:1141–1154.e7.

73. Fedorova V, Pospisilova V, Vanova T, Amruz Cerna K, Abaffy P, Sedmik J, et al. Glioblastoma and cerebral organoids: development and analysis of an in vitro model for glioblastoma migration. Mol Oncol [Internet]. [cited 2023 Mar 14];n/a. Available from: https://onlinelibrary.wiley.com/doi/abs/10.1002/1878-0261.13389

74. Nezvedová M, Jha D, Váňová T, Gadara D, Klímová H, Raška J, et al. Single Cerebral Organoid Mass Spectrometry of Cell-Specific Protein and Glycosphingolipid Traits. Anal Chem. American Chemical Society; 2023;95:3160–7.

75. Fedorova V, Vanova T, Elrefae L, Pospisil J, Petrasova M, Kolajova V, et al. Differentiation of neural rosettes from human pluripotent stem cells in vitro is sequentially regulated on a molecular level and accomplished by the mechanism reminiscent of secondary neurulation. Stem Cell Res. 2019;40:101563.

76. Miranda AM, Bravo FV, Chan RB, Sousa N, Di Paolo G, Oliveira TG. Differential lipid composition and regulation along the hippocampal longitudinal axis. Transl Psychiatry. 2019;9:144.

77. Huynh K, Barlow CK, Jayawardana KS, Weir JM, Mellett NA, Cinel M, et al. High-Throughput Plasma Lipidomics: Detailed Mapping of the Associations with Cardiometabolic Risk Factors. Cell Chem Biol. 2019;26:71–84.e4.

78. Xuan Q, Hu C, Yu D, Wang L, Zhou Y, Zhao X, et al. Development of a High Coverage Pseudotargeted Lipidomics Method Based on Ultra-High Performance Liquid Chromatography–Mass Spectrometry. Anal Chem. American Chemical Society; 2018;90:7608–16.

79. Chetty PS, Mayne L, Lund-Katz S, Englander SW, Phillips MC. Helical structure, stability, and dynamics in human apolipoprotein E3 and E4 by hydrogen exchange and mass spectrometry. Proc Natl Acad Sci. Proceedings of the National Academy of Sciences; 2017;114:968–73.

80. Lindner K, Beckenbauer K, van Ek LC, Titeca K, de Leeuw SM, Awwad K, et al. Isoform-and cell-state-specific lipidation of ApoE in astrocytes. Cell Rep. 2022;38:110435.

81. Proitsi P, Kim M, Whiley L, Pritchard M, Leung R, Soininen H, et al. Plasma lipidomics analysis finds long chain cholesteryl esters to be associated with Alzheimer’s disease. Transl Psychiatry. 2015;5:e494.

81. MahmoudianDehkordi S, Ahmed AT, Bhattacharyya S, Han X, Baillie RA, Arnold M, et al. Alterations in acylcarnitines, amines, and lipids inform about the mechanism of action of citalopram/escitalopram in major depression.

83. Bossu P, Salani F, Ciaramella A, Sacchinelli E, Mosca A, Banaj N, et al. Anti-inflammatory Effects of Homotaurine in Patients With Amnestic Mild Cognitive Impairment. Front Aging Neurosci. Frontiers Media SA; 2018;10:1–8.

84. Donovan EL, Pettine SM, Hickey MS, Hamilton KL, Miller BF. Lipidomic analysis of human plasma reveals ether-linked lipids that are elevated in morbidly obese humans compared to lean. Diabetol Metab Syndr. BioMed Central; 2013;5:1–13.

85. Huang Y-WA, Zhou B, Nabet AM, Wernig M, Südhof TC. Differential Signaling Mediated by ApoE2, ApoE3, and ApoE4 in Human Neurons Parallels Alzheimer’s Disease Risk. J Neurosci. Society for Neuroscience; 2019;39:7408–27.

86. Berson A, Barbash S, Shaltiel G, Goll Y, Hanin G, Greenberg DS, et al. Cholinergic-associated loss of hnRNP-A/B in Alzheimer’s disease impairs cortical splicing and cognitive function in mice. EMBO Mol Med. John Wiley & Sons, Ltd; 2012;4:730–42.

87. Uddin MdS, Kabir MdT, Al Mamun A, Abdel-Daim MM, Barreto GE, Ashraf GM. APOE and Alzheimer’s Disease: Evidence Mounts that Targeting APOE4 may Combat Alzheimer’s Pathogenesis. Mol Neurobiol. 2019;56:2450–65.

88. Martens YA, Zhao N, Liu C-C, Kanekiyo T, Yang AJ, Goate AM, et al. ApoE Cascade Hypothesis in the pathogenesis of Alzheimer’s disease and related dementias. Neuron. 2022;110:1304–17.

89. Fitz NF, Cronican AA, Saleem M, Fauq AH, Chapman R, Lefterov I, et al. Abca1 deficiency affects Alzheimer’s disease-like phenotype in human ApoE4 but not in ApoE3-targeted replacement mice. J Neurosci Off J Soc Neurosci. 2012;32:13125–36.

90. Rawat V, Wang S, Sima J, Bar R, Liraz O, Gundimeda U, et al. ApoE4 Alters ABCA1 Membrane Trafficking in Astrocytes. J Neurosci. Society for Neuroscience; 2019;39:9611–22.

91. Lanfranco MF, Ng CA, Rebeck GW. ApoE Lipidation as a Therapeutic Target in Alzheimer’s Disease. Int J Mol Sci. 2020;21:6336.

92. Liao F, Li A, Xiong M, Bien-Ly N, Jiang H, Zhang Y, et al. Targeting of nonlipidated, aggregated apoE with antibodies inhibits amyloid accumulation. J Clin Invest. 2018;128:2144–55.

93. Huang Y, von Eckardstein A, Wu S, Maeda N, Assmann G. A plasma lipoprotein containing only apolipoprotein E and with gamma mobility on electrophoresis releases cholesterol from cells. Proc Natl Acad Sci U S A. 1994;91:1834–8.

94. Burgess JW, Gould DR, Marcel YL. The HepG2 extracellular matrix contains separate heparinase-and lipid-releasable pools of ApoE. Implications for hepatic lipoprotein metabolism. J Biol Chem. 1998;273:5645–54.

95. DeMattos RB, Curtiss LK, Williams DL. A Minimally Lipidated Form of Cell-derived Apolipoprotein E Exhibits Isoform-specific Stimulation of Neurite Outgrowth in the Absence of Exogenous Lipids or Lipoproteins*. J Biol Chem. 1998;273:4206–12.

96. LaDu MJ, Stine WB, Narita M, Getz GS, Reardon CA, Bu G. Self-assembly of HEK cell-secreted ApoE particles resembles ApoE enrichment of lipoproteins as a ligand for the LDL receptor-related protein. Biochemistry. 2006;45:381–90.

97. Koldamova R, Staufenbiel M, Lefterov I. Lack of ABCA1 considerably decreases brain ApoE level and increases amyloid deposition in APP23 mice. J Biol Chem. 2005;280:43224–35.

98. Wahrle SE, Jiang H, Parsadanian M, Hartman RE, Bales KR, Paul SM, et al. Deletion of Abca1 increases Abeta deposition in the PDAPP transgenic mouse model of Alzheimer disease. J Biol Chem. 2005;280:43236–42.

99. Dong LM, Wilson C, Wardell MR, Simmons T, Mahley RW, Weisgraber KH, et al. Human apolipoprotein E. Role of arginine 61 in mediating the lipoprotein preferences of the E3 and E4 isoforms. J Biol Chem. 1994;269:22358–65.

100. Dong LM, Weisgraber KH. Human apolipoprotein E4 domain interaction. Arginine 61 and glutamic acid 255 interact to direct the preference for very low density lipoproteins. J Biol Chem. 1996;271:19053–7.

101. Ray A, Ahalawat N, Mondal J. Atomistic Insights into Structural Differences between E3 and E4 Isoforms of Apolipoprotein E. Biophys J. 2017;113:2682–94.

102. Morrow JA, Segall ML, Lund-Katz S, Phillips MC, Knapp M, Rupp B, et al. Differences in stability among the human apolipoprotein E isoforms determined by the amino-terminal domain. Biochemistry. 2000;39:11657–66.

103. Acharya P, Segall ML, Zaiou M, Morrow J, Weisgraber KH, Phillips MC, et al. Comparison of the stabilities and unfolding pathways of human apolipoprotein E isoforms by differential scanning calorimetry and circular dichroism. Biochim Biophys Acta BBA - Mol Cell Biol Lipids. 2002;1584:9–19.

104. Raulin A-C, Kraft L, Al-Hilaly YK, Xue W-F, McGeehan JE, Atack JR, et al. The Molecular Basis for Apolipoprotein E4 as the Major Risk Factor for Late-Onset Alzheimer’s Disease. J Mol Biol. 2019;431:2248–65.

105. Dong J, Peters-Libeu CA, Weisgraber KH, Segelke BW, Rupp B, Capila I, et al. Interaction of the N- terminal domain of apolipoprotein E4 with heparin. Biochemistry. 2001;40:2826–34.

106. Segelke BW, Forstner M, Knapp M, Trakhanov SD, Parkin S, Newhouse YM, et al. Conformational flexibility in the apolipoprotein E amino-terminal domain structure determined from three new crystal forms: implications for lipid binding. Protein Sci Publ Protein Soc. 2000;9:886–97.

107. Huang RY-C, Garai K, Frieden C, Gross ML. Hydrogen/Deuterium Exchange and Electron-Transfer Dissociation Mass Spectrometry Determine the Interface and Dynamics of Apolipoprotein E Oligomerization. Biochemistry. 2011;50:9273–82.

108. Gau B, Garai K, Frieden C, Gross ML. Mass Spectrometry-Based Protein Footprinting Characterizes the Structures of Oligomeric Apolipoprotein E2, E3, and E4. Biochemistry. American Chemical Society; 2011;50:8117–26.

109. Frieden C. ApoE: the role of conserved residues in defining function. Protein Sci Publ Protein Soc. 2015;24:138–44.

110. Yamada H, Tamada T, Kosaka M, Miyata K, Fujiki S, Tano M, et al. ‘Crystal lattice engineering,’ an approach to engineer protein crystal contacts by creating intermolecular symmetry: Crystallization and structure determination of a mutant human RNase 1 with a hydrophobic interface of leucines. Protein Sci Publ Protein Soc. 2007;16:1389–97.

111. Wang C, Najm R, Xu Q, Jeong D-E, Walker D, Balestra ME, et al. Gain of toxic apolipoprotein E4 effects in human iPSC-derived neurons is ameliorated by a small-molecule structure corrector. Nat Med. 2018;24:647–57.

112. Alzheon Inc. A Phase 3, Multicenter, Randomized, Double-blind, Placebo-controlled Study of the Efficacy, Safety and Biomarker Effects of ALZ-801 in Subjects With Early Alzheimer’s Disease and APOE4/4 Genotype [Internet]. clinicaltrials.gov; 2022 Jun. Report No.: NCT04770220. Available from: https://clinicaltrials.gov/ct2/show/NCT04770220

113. Blanchard JW, Akay LA, Davila-Velderrain J, von Maydell D, Mathys H, Davidson SM, et al. APOE4 impairs myelination via cholesterol dysregulation in oligodendrocytes. Nature. Nature Publishing Group; 2022;1–11.

114. van der Kant R, Langness VF, Herrera CM, Williams DA, Fong LK, Leestemaker Y, et al. Cholesterol Metabolism Is a Druggable Axis that Independently Regulates Tau and Amyloid-β in iPSC- Derived Alzheimer’s Disease Neurons. Cell Stem Cell. 2019;24:363–375.e9.

115. Liu C-C, Zhao J, Fu Y, Inoue Y, Ren Y, Chen Y, et al. Peripheral apoE4 enhances Alzheimer’s pathology and impairs cognition by compromising cerebrovascular function. Nat Neurosci. Nature Publishing Group; 2022;25:1020–33.

116. Yamazaki Y, Zhao N, Caulfield TR, Liu C-C, Bu G. Apolipoprotein E and Alzheimer disease: pathobiology and targeting strategies. Nat Rev Neurol. 2019;15:501–18.

117. Wang L, Dou Z. Apolipoprotein E regulates chromatin stability and senescence. Nat Aging. Nature Publishing Group; 2022;2:282–4.

