## Supplementary Information File for "Domino-like Effect of C112R Mutation on ApoE4 Aggregation and Its Reduction by Alzheimer’s Disease Drug Candidate": Nemergut-etal-2023-SI-R1-no-labels.docx

^1^ Loschmidt Laboratories, Department of Experimental Biology, Faculty of Science, Masaryk University, Kamenice 5, 625 00 Brno, Czech Republic; ^2^ RECETOX, Faculty of Science, Masaryk University, Kamenice 5, 625 00 Brno, Czech Republic; ^3^ International Clinical Research Center, St. Anne's University Hospital Brno, Pekarska 53, 656 91 Brno, Czech Republic; ^4^ Center for Interdisciplinary Biosciences, Technology and Innovation Park, P. J. Safarik University in Kosice, Trieda SNP 1, 04011 Kosice, Slovakia; ^5^ Research Centre for Applied Molecular Oncology, Masaryk Memorial Cancer Institute, Zluty kopec 7, 656 53 Brno, Czech Republic; ^6^ Department of Histology and Embryology, Faculty of Medicine, Kamenice 5, 625 00 Brno, Czech Republic

### **SUPPLEMENTARY** TABLES

**Supplementary Table 1:** Primers used in PCR reactions.

| ApoE + W34A | |
| --- | --- |
| forward | 5'-GCACTGGGTCGTTTTGCGGATTATCTGCGTTGG-3' |
| reverse | 5'-GCTAGTTATTGCTCAGCGG-3' |
| ApoE + R38A/E45A/E49A/R145A | |
| forward | 5'-GGTCGTTTTTGGGATTATCTGGCGTGGGTTCAGACCCTGAGCGCGCAGGTTCA  AGCGGAACTGCTGAGCAGCCAGGTT-3' |
| reverse | 5'-ATCGGCATCACGCAGCAGACGTTTCGCCAGTTTACGCAGATGGCTTGCCAG-3' |
| ApoE_M_-AP | |
|  | 5'-GGAGGACATATGGGTCAGCGTTGGGAATTAGC-3' |
|  | 5'-TCCTCCGGATCCTTAATGATTATCGCTCGG-3' |

**Supplementary Table 2:** Lipid and metabolite standards and their concentration (ng/mL).

| **Standards** | **Conc. (ng/mL)** |
| --- | --- |
| 15:0-18:1(d7) PC | 640 |
| 15:0-18:1(d7) PE | 20 |
| 15:0-18:1(d7) PG | 120 |
| 17:0-14:1 PS | 20 |
| 15:0-18:1(d7) PI | 40 |
| 18:1(d7) Lyso PC | 100 |
| 18:1(d7) Lyso PE | 20 |
| 18:1(d7) Chol Ester | 1400 |
| 15:0-18:1(d7) DAG | 40 |
| 15:0-18:1(d7)-15:0 TAG | 220 |
| d18:1-18:1(d9) SM | 120 |
| C16 (d3) Carnitine | 0.33 |
| d18:1/12:0 Cer | 25 |
| d18:1/12:0 Lac Cer | 25 |
| d18:1/12:0 Gla Cer | 25 |
| (d7) Cholesterol | 12500 |

**Supplementary Table 3:** SRM transitions used for the quantitation of gangliosides (d18:1/18:0) based on optimal sensitivity and selectivity.

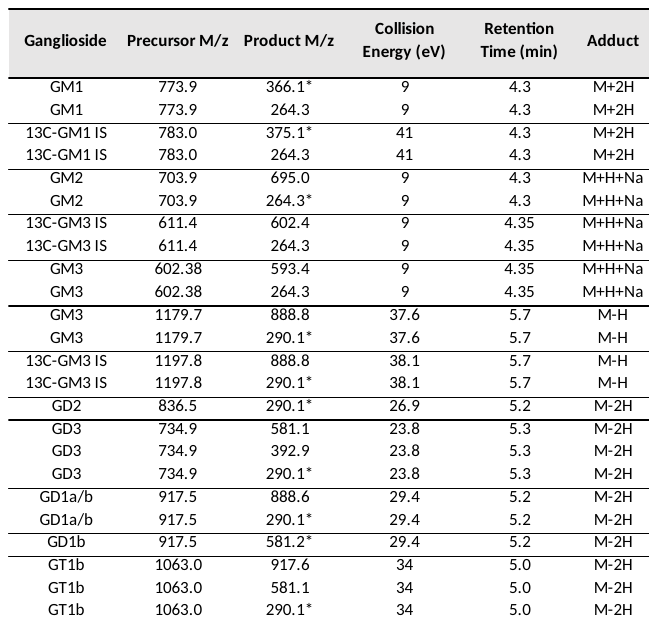

*MRM transitions used for the quantitation of the GSs (d18:1/18:0) based on optimal sensitivity and selectivity.

**Supplementary Table 4.** List of proteins quantified in cerebral organoids.

| Gene name | Full name | Protein number |
| --- | --- | --- |
| *ACTB* | Beta-actin (Actin, cytoplasmic 1) | P60709 |
| *ApoE* | Apolipoprotein E | P02649 |
| *CCT6A* | T-complex protein 1 subunit zeta | P40227* |
| *CD44* | CD44 antigen | P16070* |
| *CDH2* | Cadherin-2 (Neural cadherin) | P19022 |
| *CLU* | Clusterin (APOJ) | P10909 |
| *CTSB* | Cathepsin B (APP secretase) | P07858 |
| *DCX* | Neuronal migration protein doublecortin | O43602* |
| *FABP7* | Fatty acid-binding protein, brain | O15540 |
| *GAPDH* | Glyceraldehyde-3-phosphate dehydrogenase | P04406 |
| *GLUL* | Glutamine synthetase | P15104 |
| *HNRNPAB* | Heterogenous nuclear ribonucleoprotein A/B | Q99729 |
| *LDHB* | L-lactate dehydrogenase B chain | P07195 |
| *MAP2* | Microtubule-associtaed protein 2 | P11137 |
| *MAPT* | Microtubule-associated protein tau | P10636 |
| *NCAM1* | Neural cell adhesion molecule 1 | P13591 |
| *NEFL* | Neurofilament light polypeptide | P07196* |
| *NEFM* | Neurofilament medium polypeptide | P07197 |
| *NES* | Nestin | P48681 |
| *NPC2* | NPC intracellular cholesterol transporter 2 | P61916 |
| *PARK7* | Parkinson disease protein 7 (DJ-1) | Q99497 |
| *PAX6* | Paired box protein Pax-6 | P26367* |
| *PFN1* | Profilin-1 | P07737 |
| *PPIA* | Peptidyl-prolyl cis-trans isomerase A | P62937 |
| *PSMD7* | 26S proteasome non-ATPase regulatory subunit 7 | P51665 |
| *S100B* | S100 calcium-binding protein B (Protein S100-B, S100b) | P04271 |
| *STMN1* | Stathmin | P16949 |
| *SYN1* | Synapsin-1 (Brain protein 4.1) | P17600* |
| *TPI1* | Triosephosphate isomerase | P60174 |
| *TTR* | Transthyretin (Prealbumin) | P02766* |
| *TUBB3* | Tubulin beta-3 chain (TUJ) | Q13509* |
| *VIM* | Vimentin | P08670 |

* Proteins not detected in each sample of a single cerebral organoid due to low abundances.

| **Supplementary Table 5:** Transitions list of monitored unique peptides of analyzed proteins. | | | | | | | | | |
| --- | --- | --- | --- | --- | --- | --- | --- | --- | --- |
| **Protein number** | **Gene name** | **Peptide Sequence** | **Precursor Adduct** | **Precursor Mz [Da]** | **Product Adduct** | **Product Mz [Da]** | **Fragment Ion** | **Retention Time [min]** | **Collision Energy [eV]** |
| P11137 | *MAP2* | LINQPLPDLK | [M+2H] | 575.8 | [M+H] | 682.4 | y6* | 11.5 | 18.9 |
| P11137 | *MAP2* | LINQPLPDLK | [M+2H] | 575.8 | [M+H] | 472.3 | y4 | 11.5 | 18.9 |
| P11137 | *MAP2* | LINQPLPDLK | [M+2H] | 575.8 | [M+H] | 260.2 | y2 | 11.5 | 18.9 |
| P11137 | *MAP2* | LINQPLPDLK | [M+2H] | 579.9 | [M+H] | 690.4 | y6* | 11.5 | 18.9 |
| P11137 | *MAP2* | LINQPLPDLK | [M+2H] | 579.9 | [M+H] | 480.3 | y4 | 11.5 | 18.9 |
| P11137 | *MAP2* | LINQPLPDLK | [M+2H] | 579.9 | [M+H] | 268.2 | y2 | 11.5 | 18.9 |
| P10636 | *MAPT* | IGSLDNITHVPGGGNK | [M+3H] | 526.9 | [M+H] | 529.3 | y6 | 8.6 | 14.2 |
| P10636 | *MAPT* | IGSLDNITHVPGGGNK | [M+3H] | 526.9 | [M+2H] | 733.4 | y15* | 8.6 | 14.2 |
| P10636 | *MAPT* | IGSLDNITHVPGGGNK | [M+3H] | 526.9 | [M+2H] | 704.9 | y14 | 8.6 | 14.2 |
| P10636 | *MAPT* | IGSLDNITHVPGGGNK | [M+3H] | 529.6 | [M+H] | 537.3 | y6 | 8.6 | 14.2 |
| P10636 | *MAPT* | IGSLDNITHVPGGGNK | [M+3H] | 529.6 | [M+2H] | 737.4 | y15* | 8.6 | 14.2 |
| P10636 | *MAPT* | IGSLDNITHVPGGGNK | [M+3H] | 529.6 | [M+2H] | 708.9 | y14 | 8.6 | 14.2 |
| P04406 | *GAPDH* | VGVNGFGR | [M+2H] | 403.2 | [M+H] | 706.4 | y7* | 4.9 | 13.5 |
| P04406 | *GAPDH* | VGVNGFGR | [M+2H] | 403.2 | [M+H] | 649.3 | y6 | 4.9 | 13.5 |
| P04406 | *GAPDH* | VGVNGFGR | [M+2H] | 403.2 | [M+H] | 550.3 | y5 | 4.9 | 13.5 |
| P04406 | *GAPDH* | VGVNGFGR | [M+2H] | 403.2 | [M+H] | 436.2 | y4 | 4.9 | 13.5 |
| P04406 | *GAPDH* | VGVNGFGR | [M+2H] | 408.2 | [M+H] | 716.4 | y7* | 4.9 | 13.5 |
| P04406 | *GAPDH* | VGVNGFGR | [M+2H] | 408.2 | [M+H] | 659.3 | y6 | 4.9 | 13.5 |
| P04406 | *GAPDH* | VGVNGFGR | [M+2H] | 408.2 | [M+H] | 560.3 | y5 | 4.9 | 13.5 |
| P04406 | *GAPDH* | VGVNGFGR | [M+2H] | 408.2 | [M+H] | 446.2 | y4 | 4.9 | 13.5 |
| P07197 | *NEFM* | SIELESVR | [M+2H] | 466.8 | [M+H] | 732.4 | y6* | 7.1 | 15.5 |
| P07197 | *NEFM* | SIELESVR | [M+2H] | 466.8 | [M+H] | 603.3 | y5 | 7.1 | 15.5 |
| P07197 | *NEFM* | SIELESVR | [M+2H] | 466.8 | [M+H] | 490.3 | y4 | 7.1 | 15.5 |
| P07197 | *NEFM* | SIELESVR | [M+2H] | 471.8 | [M+H] | 742.4 | y6* | 7.1 | 15.5 |
| P07197 | *NEFM* | SIELESVR | [M+2H] | 471.8 | [M+H] | 613.4 | y5 | 7.1 | 15.5 |
| P07197 | *NEFM* | SIELESVR | [M+2H] | 471.8 | [M+H] | 500.3 | y4 | 7.1 | 15.5 |
| P02649 | *ApoE* | LAVYQAGAR | [M+2H] | 474.8 | [M+H] | 764.4 | y7* | 4.3 | 15.7 |
| P02649 | *ApoE* | LAVYQAGAR | [M+2H] | 474.8 | [M+H] | 665.3 | y6 | 4.3 | 15.7 |
| P02649 | *ApoE* | LAVYQAGAR | [M+2H] | 474.8 | [M+H] | 374.2 | y4 | 4.3 | 15.7 |
| P02649 | *ApoE* | LAVYQAGAR | [M+2H] | 479.8 | [M+H] | 774.4 | y7* | 4.3 | 15.7 |
| P02649 | *ApoE* | LAVYQAGAR | [M+2H] | 479.8 | [M+H] | 675.3 | y6 | 4.3 | 15.7 |
| P02649 | *ApoE* | LAVYQAGAR | [M+2H] | 479.8 | [M+H] | 384.2 | y4 | 4.3 | 15.7 |
| P61916 | *NPC2* | SGINC[+57]PIQK | [M+2H] | 508.8 | [M+H] | 759.4 | y6* | 3.2 | 16.8 |
| P61916 | *NPC2* | SGINC[+57]PIQK | [M+2H] | 508.8 | [M+H] | 645.3 | y5 | 3.2 | 16.8 |
| P61916 | *NPC2* | SGINC[+57]PIQK | [M+2H] | 512.8 | [M+H] | 767.4 | y6* | 3.2 | 16.8 |
| P61916 | *NPC2* | SGINC[+57]PIQK | [M+2H] | 512.8 | [M+H] | 653.4 | y5 | 3.2 | 16.8 |
| P13591 | *NCAM1* | ALSSEWKPEIR | [M+3H] | 439.2 | [M+H] | 514.3 | y4 | 8.3 | 11 |
| P13591 | *NCAM1* | ALSSEWKPEIR | [M+3H] | 439.2 | [M+2H] | 566.3 | y9* | 8.3 | 11 |
| P13591 | *NCAM1* | ALSSEWKPEIR | [M+3H] | 439.2 | [M+2H] | 522.8 | y8 | 8.3 | 11 |
| P13591 | *NCAM1* | ALSSEWKPEIR | [M+3H] | 442.6 | [M+H] | 524.3 | y4 | 8.3 | 11 |
| P13591 | *NCAM1* | ALSSEWKPEIR | [M+3H] | 442.6 | [M+2H] | 571.3 | y9* | 8.3 | 11 |
| P13591 | *NCAM1* | ALSSEWKPEIR | [M+3H] | 442.6 | [M+2H] | 527.8 | y8 | 8.3 | 11 |
| P10909 | *CLU* | ELDESLQVAER | [M+2H] | 644.8 | [M+H] | 802.4 | y7* | 8.9 | 21 |
| P10909 | *CLU* | ELDESLQVAER | [M+2H] | 644.8 | [M+H] | 715.4 | y6 | 8.9 | 21 |
| P10909 | *CLU* | ELDESLQVAER | [M+2H] | 644.8 | [M+H] | 602.3 | y5 | 8.9 | 21 |
| P10909 | *CLU* | ELDESLQVAER | [M+2H] | 649.8 | [M+H] | 812.5 | y7* | 8.9 | 21 |
| P10909 | *CLU* | ELDESLQVAER | [M+2H] | 649.8 | [M+H] | 725.4 | y6 | 8.9 | 21 |
| P10909 | *CLU* | ELDESLQVAER | [M+2H] | 649.8 | [M+H] | 612.3 | y5 | 8.9 | 21 |
| Transition used for relative quantification are highlighted with asterisks. | | | | | | | | | |

**Supplementary Table 6.** Crystallographic data collection and refinement statistics.*

|  | ApoE4-NTD | ApoE4-NTD | ApoE4-NTD | ApoE4-NTD |
| --- | --- | --- | --- | --- |
| **Data collection** |  |  |  |  |
| Wavelength (Å) | 1 | 1 | 1 | 1 |
| Space group | *P*2_1_2_1_2_1_ | *P*3_1_21 | *P*2_1_2_1_2_1_ | *P*2_1_2_1_2_1_ |
| Cell dimensions |  |  |  |  |
| a, b, c (Å) | 45.593, 53.175, 73.92 | 47.693, 47.693, 104.773 | 45.529, 53.085, 73.171 | 45.567, 53.025, 72.577 |
| α, β, γ (°) | 90, 90, 90 | 90, 90, 120 | 90, 90, 90 | 90, 90, 90 |
| Resolution (Å) | 36.96 - 1.549 (1.604 - 1.549) | 34.92 - 1.551 (1.606 - 1.551) | 38.66 - 1.75 (1.813 - 1.75) | 36.29 - 1.9 (1.966 - 1.9) |
| Total reflections | 348,050 (33,729) | 197,703 (8,741) | 237,062 (21,866) | 149,445 (3,275) |
| Unique reflections | 26,821 (2,643) | 20,689 (2,007) | 18,473 (1,799) | 13,462 (890) |
| Rmerge | 0.064 (1.669) | 0.066 (1.429) | 0.099 (2.04) | 0.04 (0.749) |
| I/σI | 24.55 (1.57) | 17.3 (1.7) | 17.3 (1.21) | 35.4 (1.6) |
| Completeness (%) | 99.98 (99.96) | 99.8 (97.1) | 99.84 (99.34) | 93.23 (63.6) |
| Multiplicity | 13 (12.8) | 9.5 (8.8) | 12.8 (12.2) | 11.1 (3.7) |
| CC (1/2) | 0.999 (0.596) | 0.999 (0.7) | 0.999 (0.597) | 1 (0.722) |
| Wilson B-factor | 22.74 | 22.27 | 28.18 | 33.95 |
| **Refinement** |  |  |  |  |
| Resolution (Å) | 36.96 - 1.549 (1.604 - 1.549) | 34.92 - 1.551 (1.606 - 1.551) | 38.66 - 1.75 (1.813 - 1.75) | 36.29 - 1.9 (1.966 - 1.9) |
| No. reflections | 26,820 (2,643) | 20,680 (2,007) | 18,473 (1,793) | 13,462 (887) |
| Rwork / Rfree (%) | 18.43 / 19.51 | 19.84 / 23.1 | 19.03 / 21.5 | 21.22 / 23.1 |
| No. atoms |  |  |  |  |
| Protein | 1,237 | 1,155 | 1,196 | 1,198 |
| Ligand | - | - | 6 | - |
| Water | 166 | 105 | 134 | 64 |
| B-factors | 31.58 | 32.3 | 37.77 | 45.71 |
| Protein | 30.58 | 31.77 | 37.09 | 45.84 |
| Ligand | - | - | 78.2 | - |
| Water | 39.04 | 38.17 | 42.07 | 43.21 |
| R.m.s. deviations |  |  |  |  |
| Bond lengths (Å) | 0.008 | 0.012 | 0.015 | 0.004 |
| Bond angles (°) | 0.96 | 1.42 | 1.23 | 0.7 |
| Ramachandran favored (%) | 97.16 | 98.47 | 95.74 | 95.74 |
| Ramachandran allowed (%) | 2.84 | 1.53 | 4.26 | 3.55 |
| Ramachandran outliers (%) | 0.00 | 0.00 | 0.00 | 0.71 |
| PDB ID code | 8AX9 | 8AX8 | 8CE0 | 8CDY |

*Values in parentheses are for the highest-resolution shell.

**Supplementary Table 7.** Comparison of crystal packing and unit cell parameters of all previously solved ApoE structures.

| **Protein** | **PDB code** | **Packing** | **Space group** | **Crystal form** | **Length (Å)** | | | **Angle (°)** | | |
| --- | --- | --- | --- | --- | --- | --- | --- | --- | --- | --- |
| ApoE2 | 1LE2 | T-shaped | P2_1_2_1_2_1_ | Ortho-1 | 41.06 | 53.94 | 83.91 | 90 | 90 | 90 |
| ApoE2 | 1NFO | T-shaped | P2_1_2_1_2_1_ | Ortho-1 | 40.61 | 53.67 | 84.80 | 90 | 90 | 90 |
| ApoE3 | 1LPE | T-shaped | P2_1_2_1_2_1_ | Ortho-1 | 40.65 | 53.96 | 85.43 | 90 | 90 | 90 |
| ApoE3 | 1NFN | T-shaped | P2_1_2_1_2_1_ | Ortho-1 | 40.70 | 53.36 | 84.51 | 90 | 90 | 90 |
| ApoE3 | 1BZ4 | T-shaped | P2_1_2_1_2_1_ | Ortho-1 | 40.78 | 53.2 | 84.78 | 90 | 90 | 90 |
| ApoE3 | 1EA8 | T-shaped | P2_1_2_1_2_1_ | Ortho-1 | 40.80 | 53.47 | 84.46 | 90 | 90 | 90 |
| ApoE3 | 1H7I | T-shaped | P2_1_2_1_2_1_ | Ortho-1 | 40.70 | 52.96 | 84.42 | 90 | 90 | 90 |
| ApoE3 | 6V7M | T-shaped | P2_1_2_1_2_1_ | Ortho-1 | 41.33 | 54.45 | 86.67 | 90 | 90 | 90 |
| ApoE3 | 1OR2 | V-shaped | P2_1_2_1_2_1_ | Ortho-2 | 47.68 | 55.59 | 63.59 | 90 | 90 | 90 |
| ApoE3 | 1OR3 | V-shaped | P3_1_21 | Trigonal | 47.37 | 47.37 | 104.54 | 90 | 90 | 120 |
| ApoE4 | 1B68 | T-shaped | P2_1_2_1_2_1_ | Ortho-1 | 40.21 | 53.21 | 84.76 | 90 | 90 | 90 |
| ApoE4 | 1LE4 | T-shaped | P2_1_2_1_2_1_ | Ortho-1 | 40.71 | 53.33 | 85.28 | 90 | 90 | 90 |
| ApoE4 | 1GS9 | V-shaped | P2_1_2_1_2_1_ | Ortho-3 | 45.51 | 53.09 | 73.37 | 90 | 90 | 90 |
| ApoE4 | 8AX9 | V-shaped | P2_1_2_1_2_1_ | Ortho-3 | 45.59 | 53.18 | 73.92 | 90 | 90 | 90 |
| ApoE4 | 8AX8 | V-shaped | P3_1_21 | Trigonal | 47.69 | 47.69 | 104.77 | 90 | 90 | 120 |
| ApoE4 | 8CDY | V-shaped | P2_1_2_1_2_1_ | Ortho-3 | 45.53 | 53.09 | 73.17 | 90 | 90 | 90 |
| ApoE4 | 8CE0 | V-shaped | P2_1_2_1_2_1_ | Ortho-3 | 45.57 | 53.03 | 72.58 | 90 | 90 | 90 |
| ApoE4+C | 6NCN | T-shaped | P2_1_2_1_2_1_ | Ortho-1 | 41.07 | 52.79 | 84.54 | 90 | 90 | 90 |
| ApoE4+C | 6NCO | T-shaped | P2_1_2_1_2_1_ | Ortho-1 | 40.96 | 52.87 | 86.44 | 90 | 90 | 90 |
| ApoEM | 7FCR | T-shaped | P2_1_2_1_2_1_ | Ortho-1 | 40.64 | 53.21 | 85.99 | 90 | 90 | 90 |
| ApoEM | 7FCS | T-shaped | P2_1_2_1_2_1_ | Ortho-1 | 40.87 | 53.49 | 85.78 | 90 | 90 | 90 |

**Supplementary Table 8.** Total interaction energies of the two chains in the dimers of ApoE3 and ApoE4, without and with SPA, and interface residues with the highest interactions.

| **System** | **Total LIE^a^**  **(kcal.mol^-1^)** | **Chain A residues^b^** | **Chain B residues^b^** |
| --- | --- | --- | --- |
| **ApoE3** | *E*_elec_ = -338 ± 84  *E*_vdW_ = -37 ± 7 | R145, R38, R142, D35, K146, R32 | E49, E45, T130, R134, E50 |
| **ApoE4** | *E*_elec_ = -396 ± 68  *E*_vdW_ = -41 ± 7 | R145, D35, R38, R142, R25, R32, K146 | E45, E49, R134, E131, T130 |
| **ApoE3+ SPA** | *E*_elec_ = -250 ± 69  *E*_vdW_ = -39 ± 7 | R145, R38, R142, R32, D35 | E49, E45, E50 |
| **ApoE4+ SPA** | *E*_elec_ = -322 ± 73  *E*_vdW_ = -37 ± 8 | R145, R38, D35, K146, R142, R32 | E49, E45, E130, R134, E131 |

^a^Total mean energies between the two monomeric units: *E*_elec_ is the electrostatic component and *E*_vdW_ is the van der Waals component of the linear interaction energy (LIE); ^b^The residues are listed in the order of the electrostatic interaction (more negative energy first). Results calculated for the first 500 ns of the respective concatenated adaptive MDs; the variability corresponds to the standard deviation of the mean over all the snapshots used.

**Supplementary Table 9.** Summary list of 64 dysregulated lipid species.

| **Lipid Species** | **E3 CO (treated vs non-treated)** | | **E4 CO (treated vs non-treated)** | |
| --- | --- | --- | --- | --- |
|  | **Fold Change** | **p-value** | **Fold Change** | **p-value** |
| CAR(12:0) | 1.23 | 0.6690 | 0.53 | 0.00250 |
| CAR(14:0) | 1.05 | 0.3680 | 0.58 | 0.00130 |
| CAR(15:0 ) | 1.27 | 0.6666 | 0.63 | 0.01140 |
| CAR(16:0) | 0.86 | 0.2801 | 0.58 | 0.00060 |
| CAR(18:0) | 0.68 | 0.0364 | 0.68 | 0.01560 |
| CAR(18:1) | 1.17 | 0.6620 | 0.60 | 0.00760 |
| CE(16:0) | 1.14 | 0.6228 | 0.50 | 0.00090 |
| CE(18:2) | 1.16 | 0.2195 | 1.32 | 0.02790 |
| CE(18:3) | 1.18 | 0.2830 | 1.49 | 0.00510 |
| CE(22:4) | 0.94 | 0.8698 | 1.54 | 0.03610 |
| CE(24:4) | 0.71 | 0.0953 | 1.59 | 0.00160 |
| DG(16:0/16:0) | 0.93 | 0.3889 | 0.69 | 0.01170 |
| DG(16:0/18:1) | 0.97 | 0.6925 | 0.76 | 0.00710 |
| DG(18:0/20:4) | 0.72 | 0.0159 | 1.31 | 0.45390 |
| TG(46:0/NL-14:0) | 0.92 | 0.4775 | 0.58 | 0.03610 |
| TG(50:0/NL-18:0) | 0.89 | 0.5435 | 0.47 | 0.02360 |
| TG(52:0/NL-16:0) | 1.05 | 0.8571 | 0.41 | 0.00990 |
| Cer(d16:1/18:0) | 0.82 | 0.3317 | 1.33 | 0.02440 |
| Cer(d18:1/14:0) | 1.02 | 0.7921 | 1.41 | 0.00770 |
| Cer(d18:1/16:0) | 0.90 | 0.7333 | 1.41 | 0.00250 |
| Cer(d18:1/18:0) | 0.70 | 0.0333 | 1.43 | 0.00760 |
| Cer(d18:1/19:0) | 0.94 | 0.8597 | 1.44 | 0.00180 |
| Cer(d18:1/20:0) | 0.71 | 0.0066 | 1.19 | 0.11150 |
| Cer(d18:1/22:0) | 0.76 | 0.0189 | 1.05 | 0.65270 |
| Cer(d18:2/18:0) | 0.85 | 0.3646 | 1.42 | 0.01220 |
| SM(31:1) | 1.68 | 0.0645 | 2.07 | 0.00360 |
| SM(35:2) | 0.91 | 0.6760 | 1.35 | 0.01180 |
| SM(36:2) | 1.15 | 0.1985 | 1.84 | 0.00020 |
| SM(38:3) | 1.26 | 0.0776 | 1.66 | 0.00020 |
| dhCer(d18:0/18:0) | 0.70 | 0.0435 | 0.90 | 0.45970 |
| dhCer(d18:0/20:0) | 0.93 | 0.0432 | 0.72 | 0.02060 |
| dhCer(d18:1/18:0) | 0.69 | 0.0212 | 1.36 | 0.03130 |
| dhCer(d18:1/20:0) | 0.70 | 0.0067 | 1.21 | 0.14180 |
| dhCer(d18:1/22:0) | 0.75 | 0.0148 | 1.02 | 0.83700 |
| HexCer(d18:1/22:0) | 0.91 | 0.3996 | 0.73 | 0.02190 |
| Hex2Cer(d18:1/18:0) | 0.79 | 0.0234 | 0.98 | 0.75760 |
| Hex2Cer(d18:1/24:0) | 1.09 | 0.8521 | 0.73 | 0.03610 |
| LPC(O-16:0) | 1.12 | 0.5585 | 0.64 | 0.00030 |
| LPC(O-18:0) | 0.97 | 0.0770 | 0.55 | <0,0001 |
| LPC(O-18:1) | 1.08 | 0.9283 | 0.71 | 0.00230 |
| PC(30:2) | 1.05 | 0.7674 | 1.46 | 0.01120 |
| PC(31:0) | 0.85 | 0.0305 | 0.85 | 0.05870 |
| PC(32:0) | 0.95 | 0.5781 | 1.18 | 0.04310 |
| PC(32:3) | 1.11 | 0.6949 | 1.60 | 0.00170 |
| PC(34:4) | 1.18 | 0.1453 | 1.45 | 0.00690 |
| PC(35:1) | 0.82 | 0.0514 | 0.66 | 0.00330 |
| PC(35:2) | 0.99 | 0.8622 | 0.82 | 0.03360 |
| PC(38:7) | 1.12 | 0.2458 | 1.26 | 0.02810 |
| PC(O-30:1) | 1.23 | 0.1292 | 1.65 | 0.04990 |
| PC(O-32:1) | 0.96 | 0.6807 | 0.79 | 0.03520 |
| PC(O-34:0) | 0.76 | 0.0020 | 0.71 | 0.00310 |
| PC(O-36:1) | 0.82 | 0.0151 | 0.66 | 0.00010 |
| PC(O-36:2) | 0.99 | 0.8748 | 0.82 | 0.01470 |
| PC(O-38:1) | 1.03 | 0.9539 | 0.75 | 0.02340 |
| PE(34:3) | 1.10 | 0.9334 | 1.54 | 0.01930 |
| PE(O-36:2) | 0.98 | 0.2815 | 0.69 | 0.00320 |
| PE(O-36:3) | 1.02 | 0.4272 | 0.74 | 0.00790 |
| PE(O-36:6) | 1.07 | 0.9846 | 0.81 | 0.03930 |
| PE(O-38:3) | 0.78 | 0.0268 | 0.59 | 0.00080 |
| PE(O-40:6) | 0.75 | 0.0155 | 0.90 | 0.24980 |
| PE(O-40:7) | 1.12 | 0.9513 | 0.76 | 0.02420 |
| PI(32:0) | 0.72 | 0.0378 | 0.59 | 0.00790 |
| PI(32:1) | 0.98 | 0.6973 | 0.73 | 0.04350 |
| PI(34:0) | 0.55 | 0.0027 | 0.83 | 0.24590 |

**Supplementary Table 10.** Abbreviations and nomenclature for analyzed lipids and proteins.

| **Abbreviation** | **Full name** |
| --- | --- |
| CAR | Carnitines |
| CE | Cholesteryl esters |
| DG | Diglycerides |
| TG | Triglycerides |
| Cer | Ceramides |
| SM | Sphingomyelins |
| dhCer | Dihydroceramides |
| HexCer | Hexosylceramides |
| Hex2Cer | Dihexosylceramides |
| Hex3Cer | Trihexosylceramides |
| LPC | Lysophosphatidylcholines |
| LPC-O | Alkyl ether-linked lysophosphatidylcholines |
| LPE | Lysophosphatidylethanolamines |
| PC | Phosphatidylcholines |
| PC-O | Alkyl ether-linked phosphatidylcholines |
| PC-P | Alkenyl ether-linked phosphatidylcholines |
| PE | Phosphatidylethanolamines |
| PE-O | Alkyl ether-linked phosphatidylethanolamines |
| PG | Phosphatidylgylcerols |
| PI | Phosphatidylinositols |
| PS | Phosphatidylserines |
| ApoE | Apolipoprotein E |
| NPC2 | Niemann-Pick proteins type C2 |
| CLU | Clusterin |
| NEFM | Neurofilament medium |
| NCAM1 | Neural cell adhesion molecule 1 |
| MAP2 | Microtubule-associated protein 2 |

Ceramide nomenclature is depicted as Lipid class (Sphingoid backbone/Fatty acid chain), with the numerals in the parenthesis providing information on the chain length and degree of saturation. Ganglioside nomenclature is based on Svennerholm system where G represents ganglioside, M – monosialo, D – disialo, and the numerals indicate the sequence of migration of gangliosides on thin-layer chromatograms.

**Supplementary Table 11.** Kinetic and equilibrium parameters obtained from the MSM analysis of the different ApoE_M_ systems concerning the unfolding of the CTD.*

|  | ApoE3_M_ | ApoE4_M_ | ApoE3_M_ + SPA | ApoE4_M_ + SPA |
| --- | --- | --- | --- | --- |
| *f*_fol_ (%) | 10.8% | 16.1% | 13.2% | 11.2% |
| RMSD_fol_ (Å) | 7.6 ± 1.5 | 9.8 ± 3.1 | 9.8 ± 0.7 | 7.6 ± 1.2 |
| *k*_unf_ (s^-1^) | 1.74 x10^7^ | 1.68 x10^6^ | 6.70 x10^6^ | 9.93 x10^6^ |
| *k*_unf_/*k*_fol_ | 150.2 | 6.54 | 20.3 | 23.9 |
| *K*_unf_ | 8.26 | 5.21 | 6.58 | 7.94 |

**f*_fol_ is the fraction of the folded state, RMSD_fol_ is the average RMSD and ± SD of the folded state, *k*_unf_ is the rate of the unfolding, *k*_fol_ if the rating of folding, and *K*_unf_ is the equilibrium constant for the unfolding process.

### SUPPLEMENTARY FIGURES

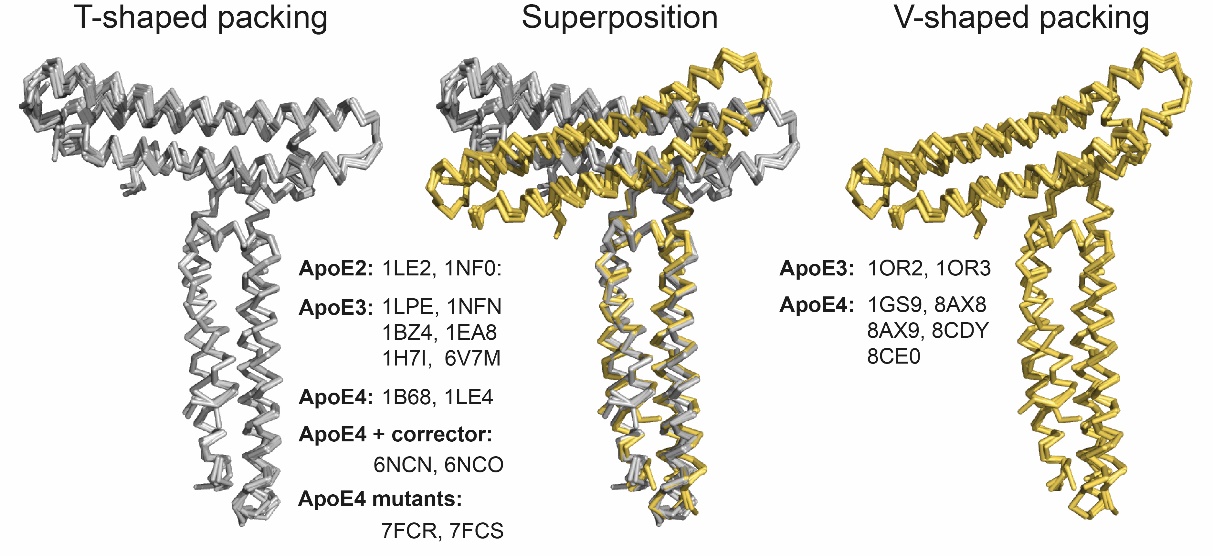

**Supplementary Figure 1.** Superposition of T-shaped and V-shaped molecular building dimeric unit of all previously solved ApoE structures.

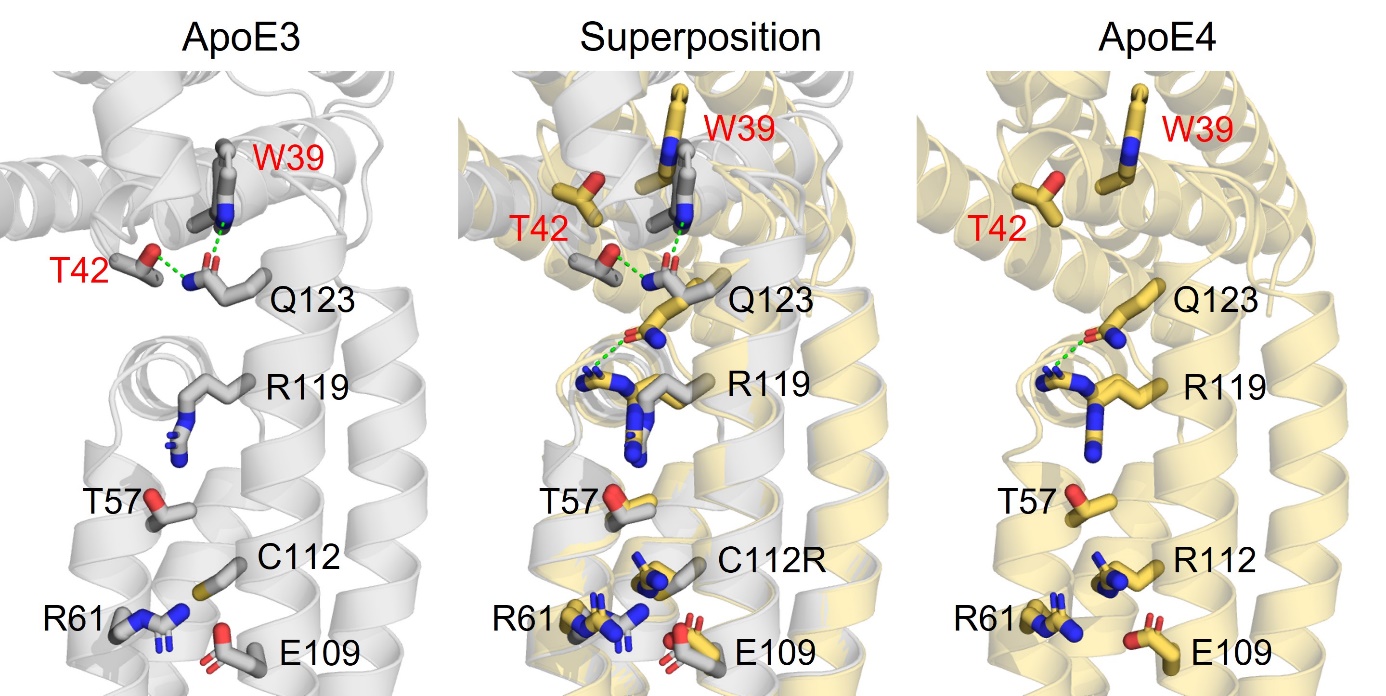

**Supplementary Figure 2.** “Domino-like” effect of C112R substitution leading from small local changes up to a change in the tilt angle between chain A and B.

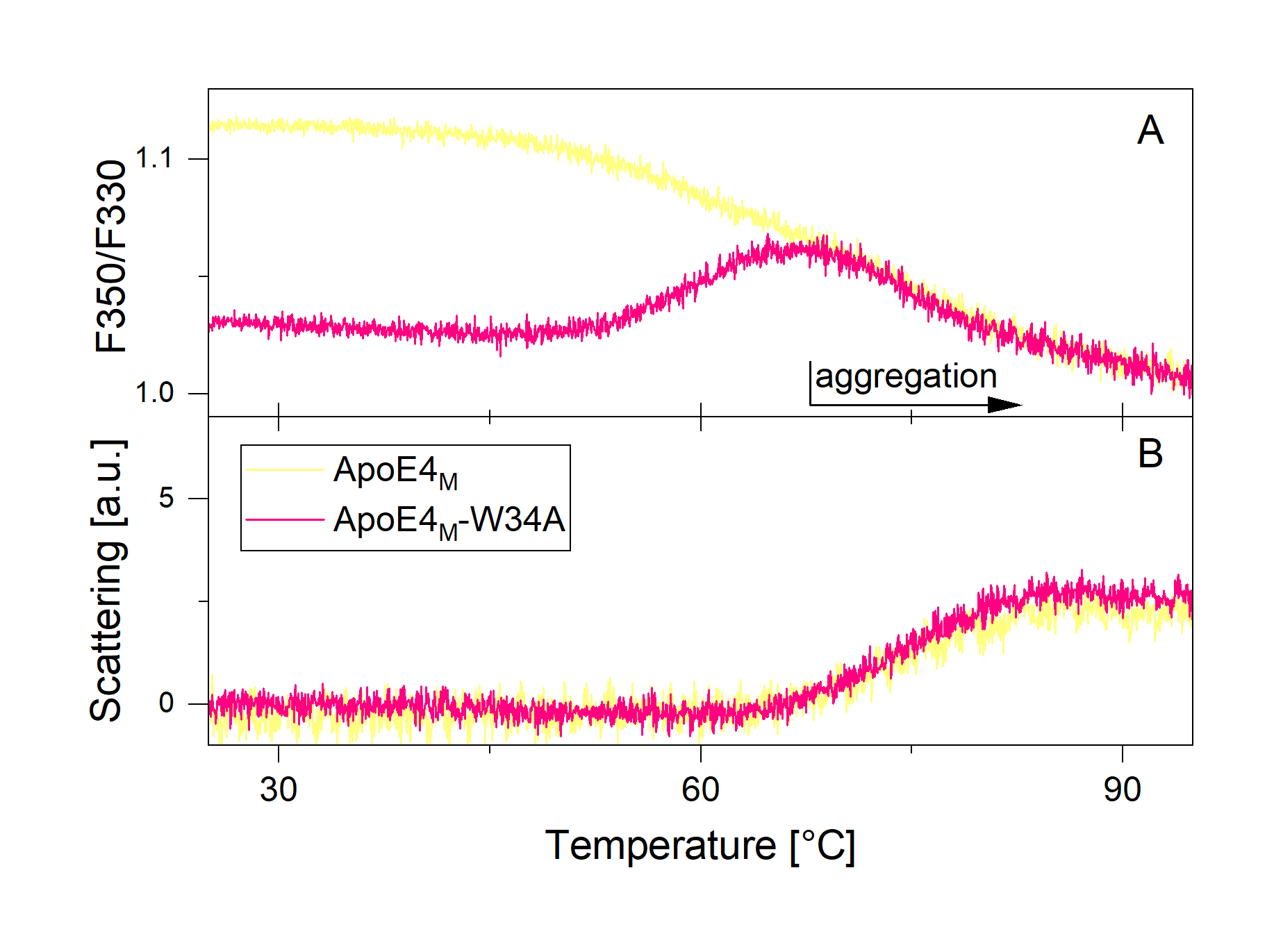

**Supplementary Figure 3.** Temperature denaturation of ApoE4_M_ (yellow) and ApoE4_M_-W34A (pink) shown as (**A**) the fluorescence intensity ratio at 330 nm and 350 nm and (**B**) light scattering measured as the attenuation of the back-reflected light intensity passing through the sample as a function of temperature. The decrease in Trp fluorescence intensity ratio (F350/F330) with increasing temperature could be explained by the covering of Trp residues during denaturation. The introduction of the W34A mutation that leads to signal inversion suggests that W34 could be responsible for the signal decrease in ApoE4_M_. The signal decrease above ~65 °C is due to sample aggregation, which is reflected as an increase in the scattered light signal.

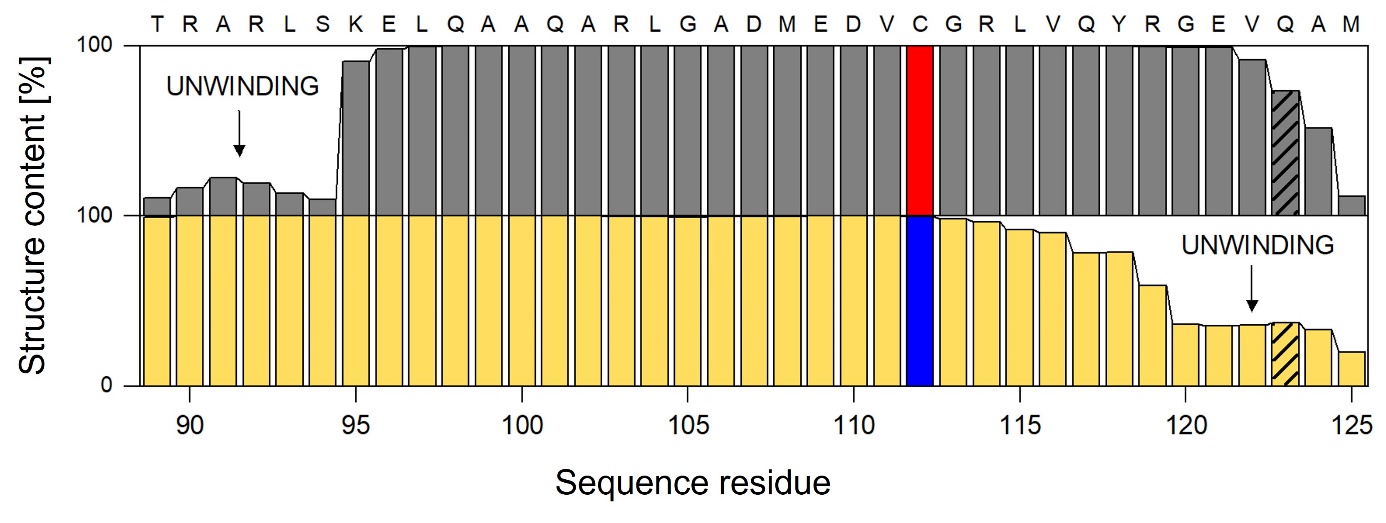

**Supplementary Figure 4.** Comparison of the α-helical content of helix H3 in ApoE3_M_ (grey) and ApoE4_M_ (yellow) calculated from the adaptive simulations. The critical residues in positions 112 and 123 discussed in the text are labelled.

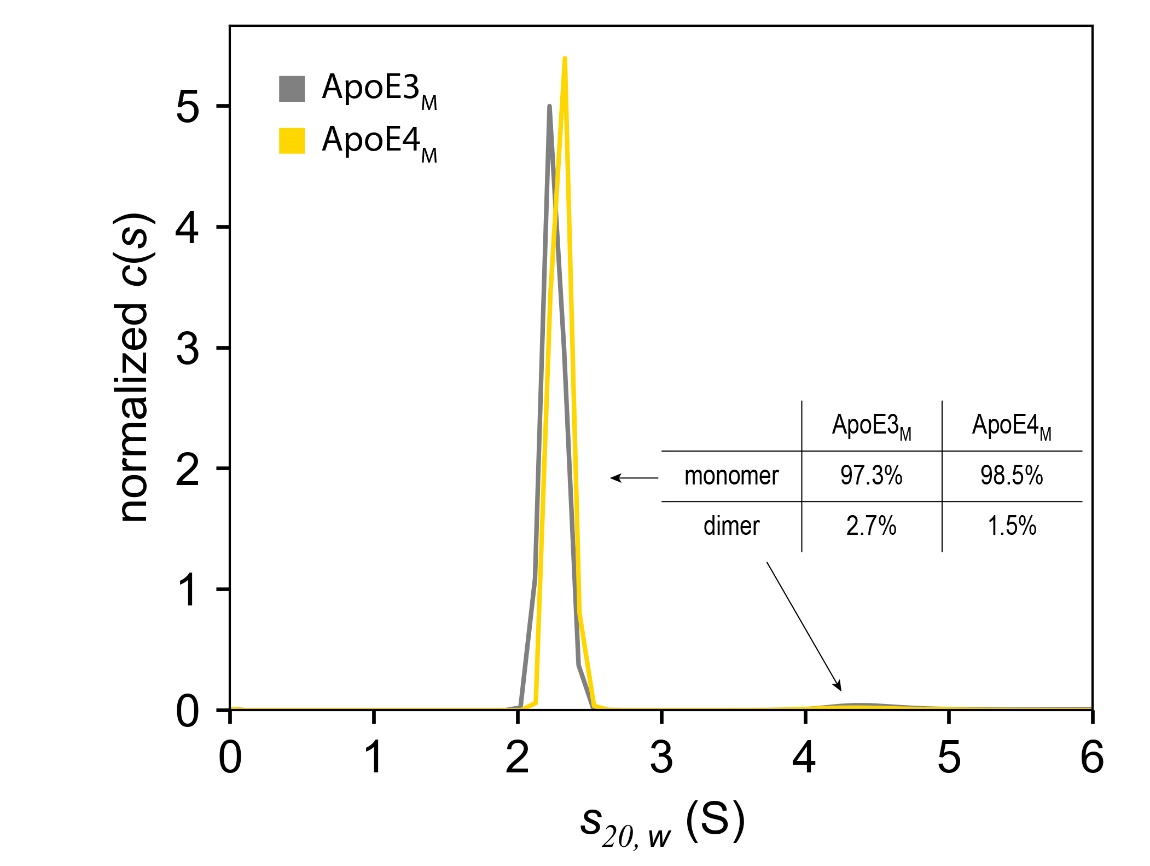

**Supplementary Figure 5.** Continuous sedimentation coefficient distribution analysis of ApoE3_M_ (grey) and ApoE4_M_ (yellow) at 0.8 mg/ml. The data were analyzed with the continuous c(s) distribution model implemented in the program Sedfit.

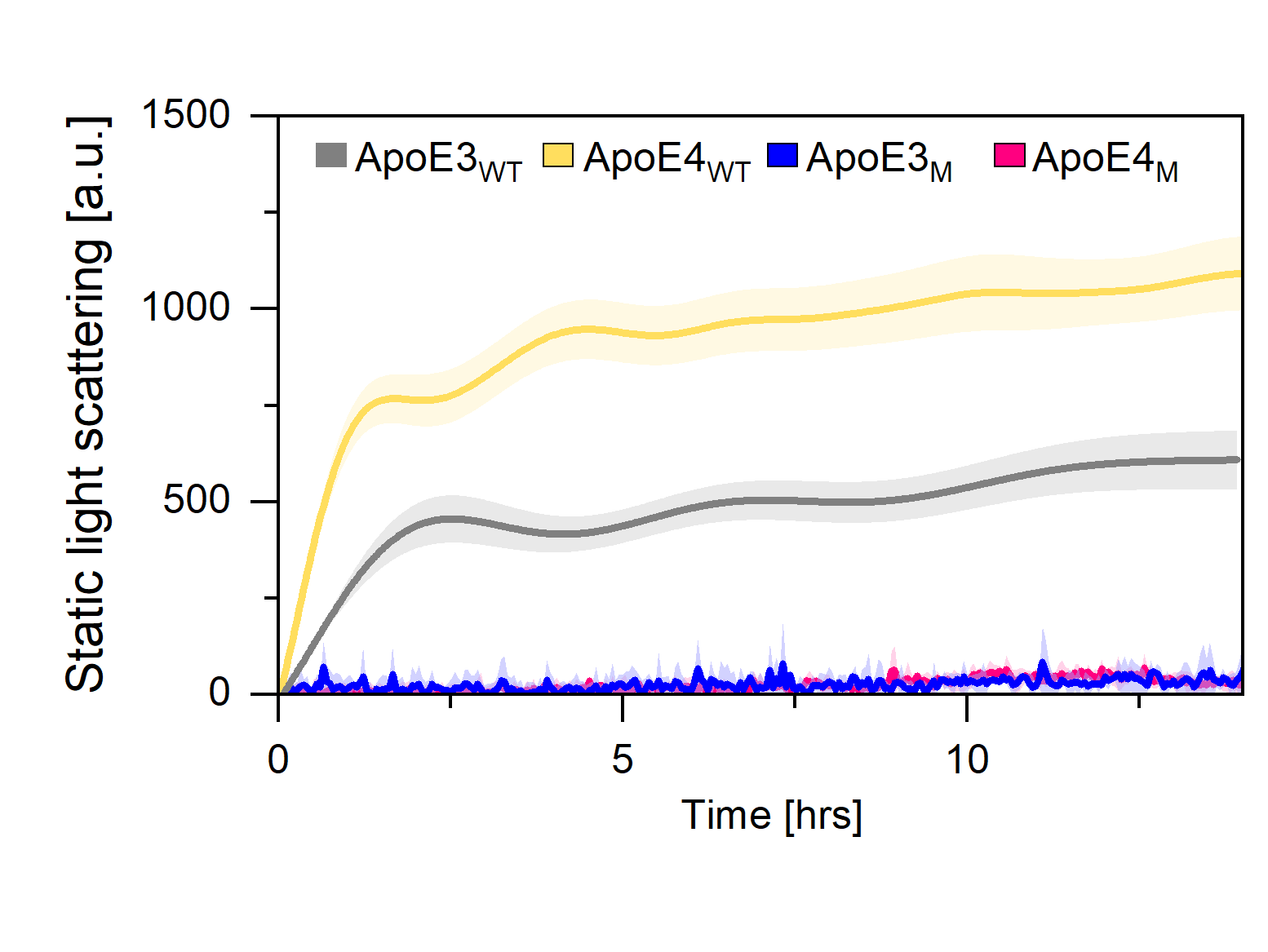

**Supplementary Figure 6.** Isothermal aggregation of ApoE3_WT_ (grey), ApoE4_WT_ (yellow), monomeric ApoE3_M_ (blue), and monomeric ApoE4_M_ (pink) demonstrates that five monomerizing mutations (F257A/W264R/V269A/L279Q/V287E) inhibit ApoE aggregation.

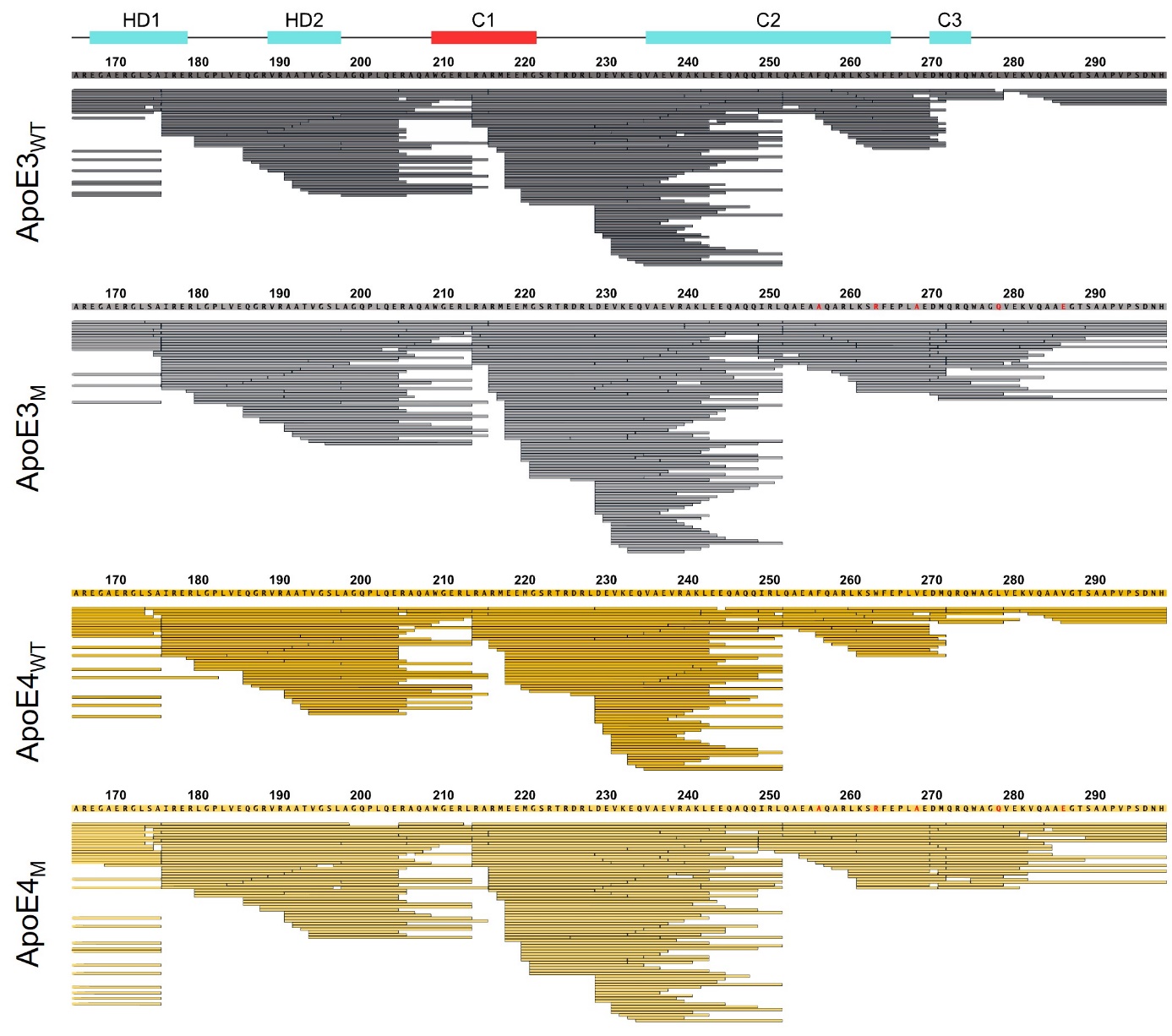

**Supplementary Figure 7.** LC-MS/MS analysis of CTD of ApoE_WT_ and ApoE_M_ proteins.

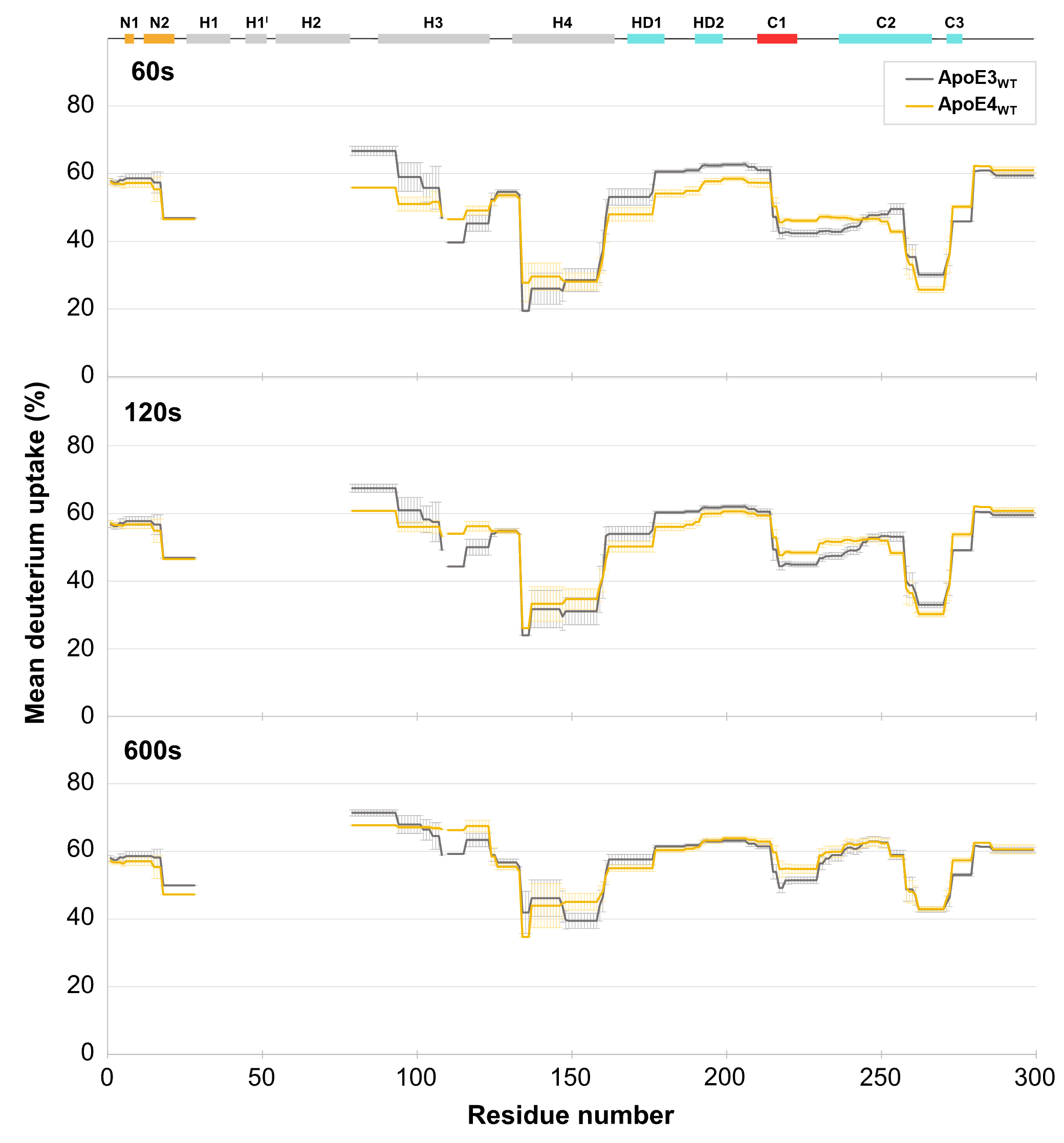

**Supplementary Figure 8.** Comparison of mean deuterium uptake of ApoE3_WT_ (grey) and ApoE4_WT_ (yellow) at 60, 120, and 600 second time points.

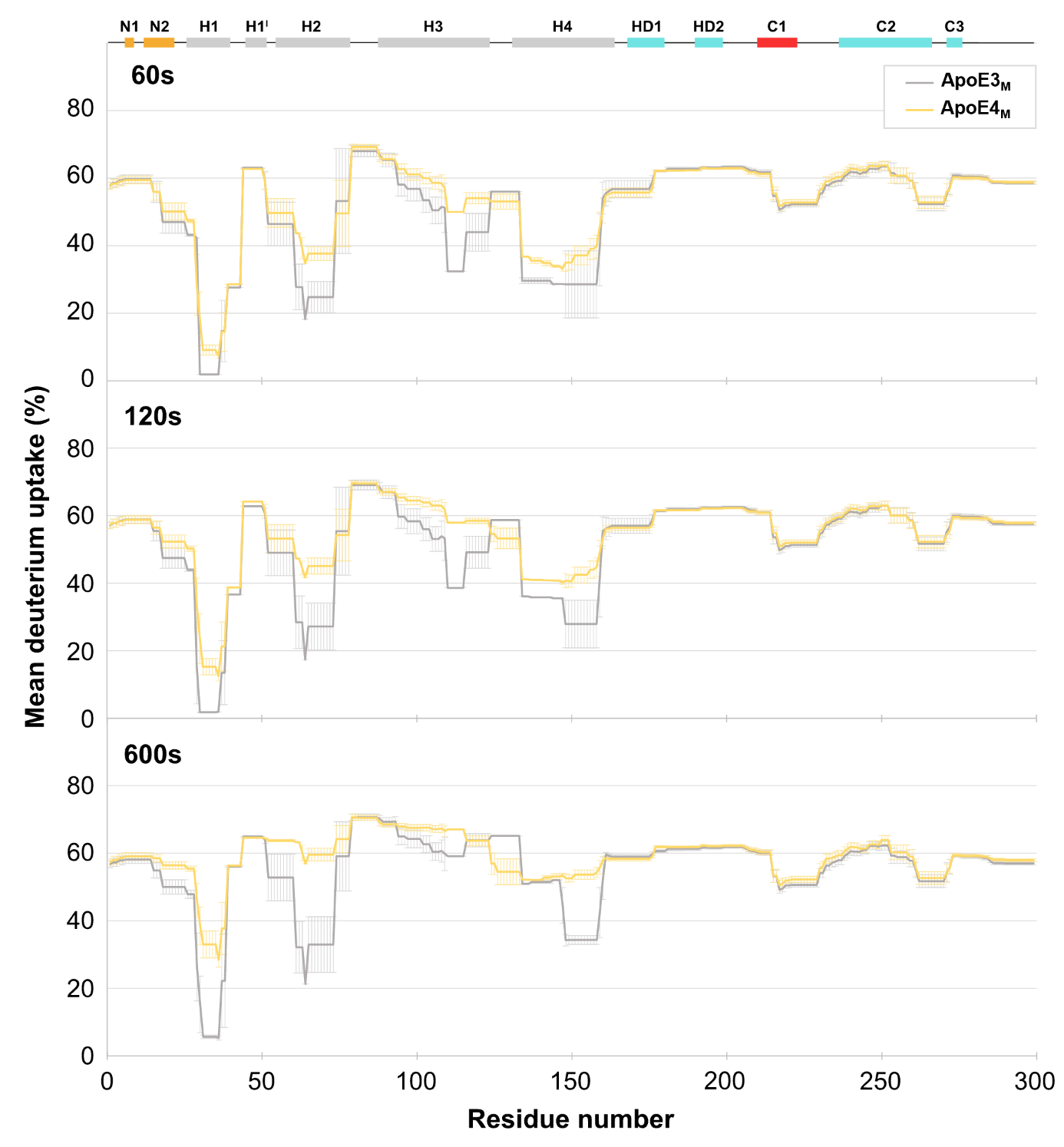

**Supplementary Figure 9.** Comparison of mean deuterium uptake of ApoE3_M_ (grey) and ApoE4_M_ (yellow) at 60, 120, and 600 second time points.

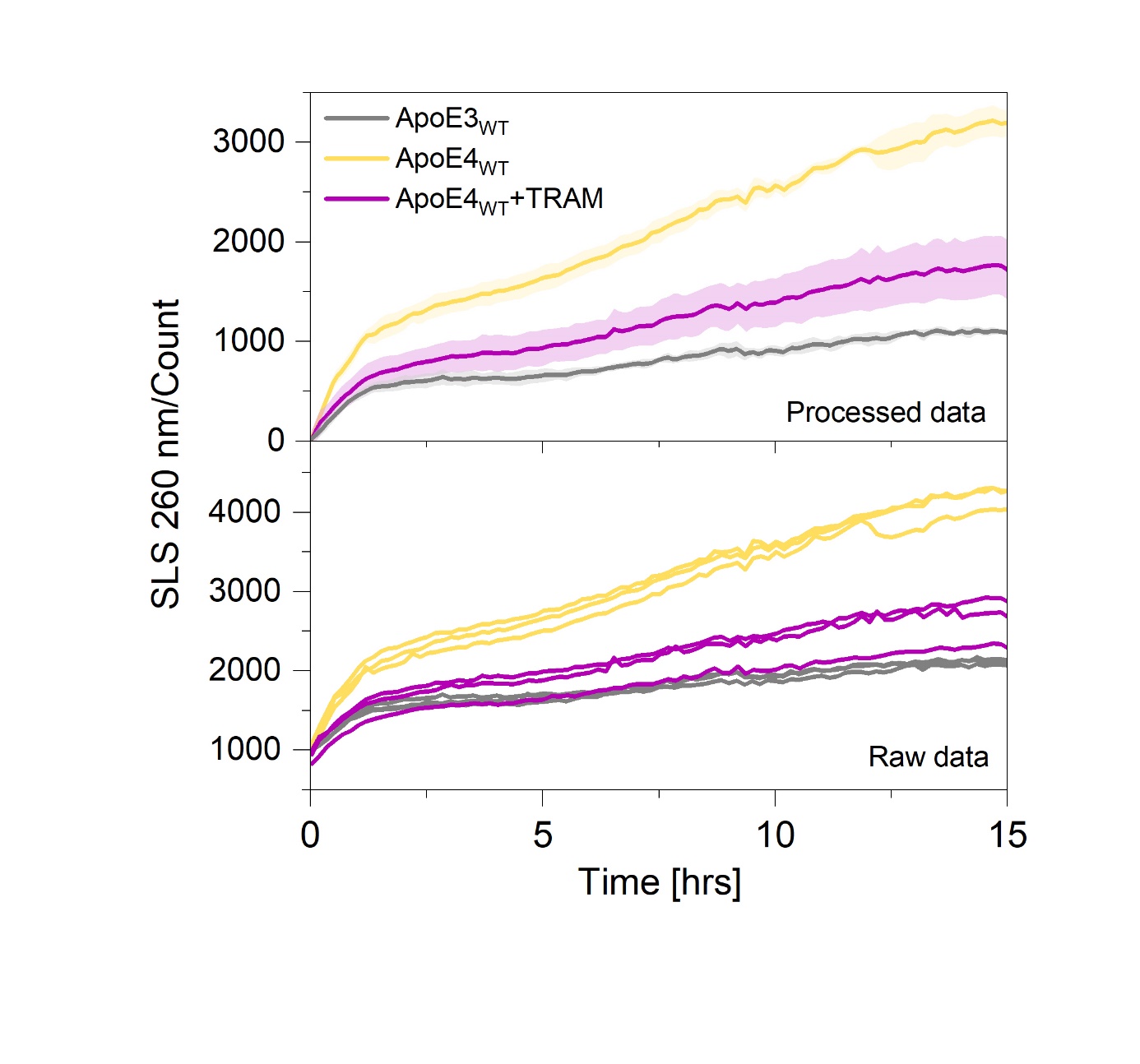

**Supplementary Figure 10.** Reduction of ApoE4 aggregation (yellow) by tramiprosate (TRAM, purple) closer to the level of ApoE3 aggregation (grey). The data were collected in Tris buffer pH 7.4 at 37 ^o^C, representing averages of 3 replicates (lower panel) with standard deviation shown as error band.

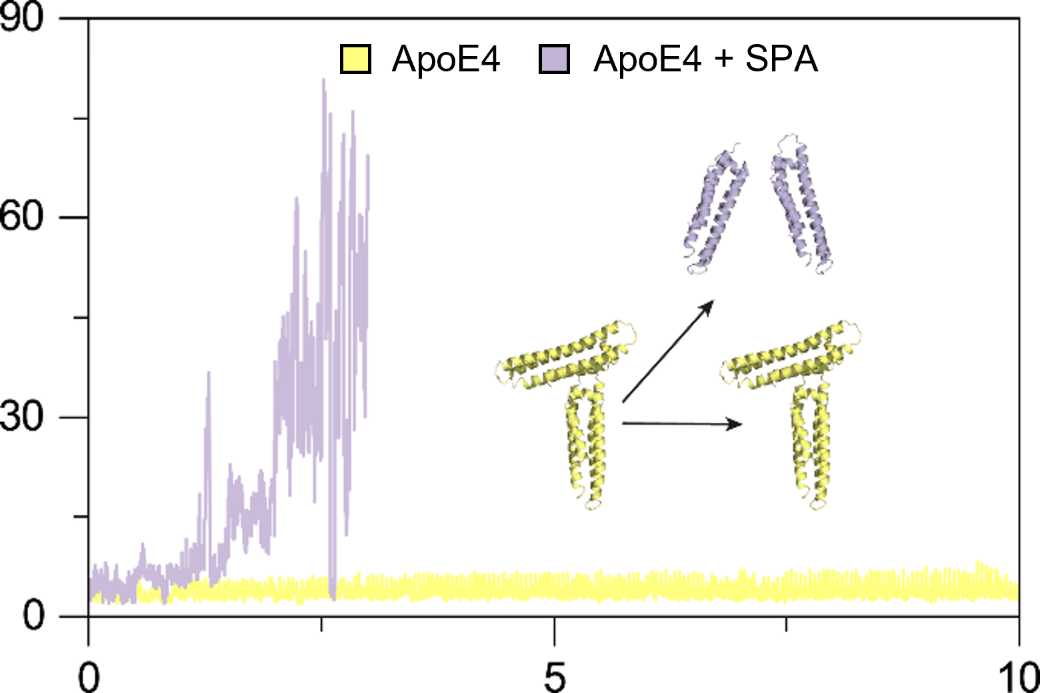

**Supplementary Figure 11.** ApoE4 V-shaped dimeric unit (yellow) and its dissociation in the presence of SPA (purple) were observed in replicated molecular dynamics simulations.

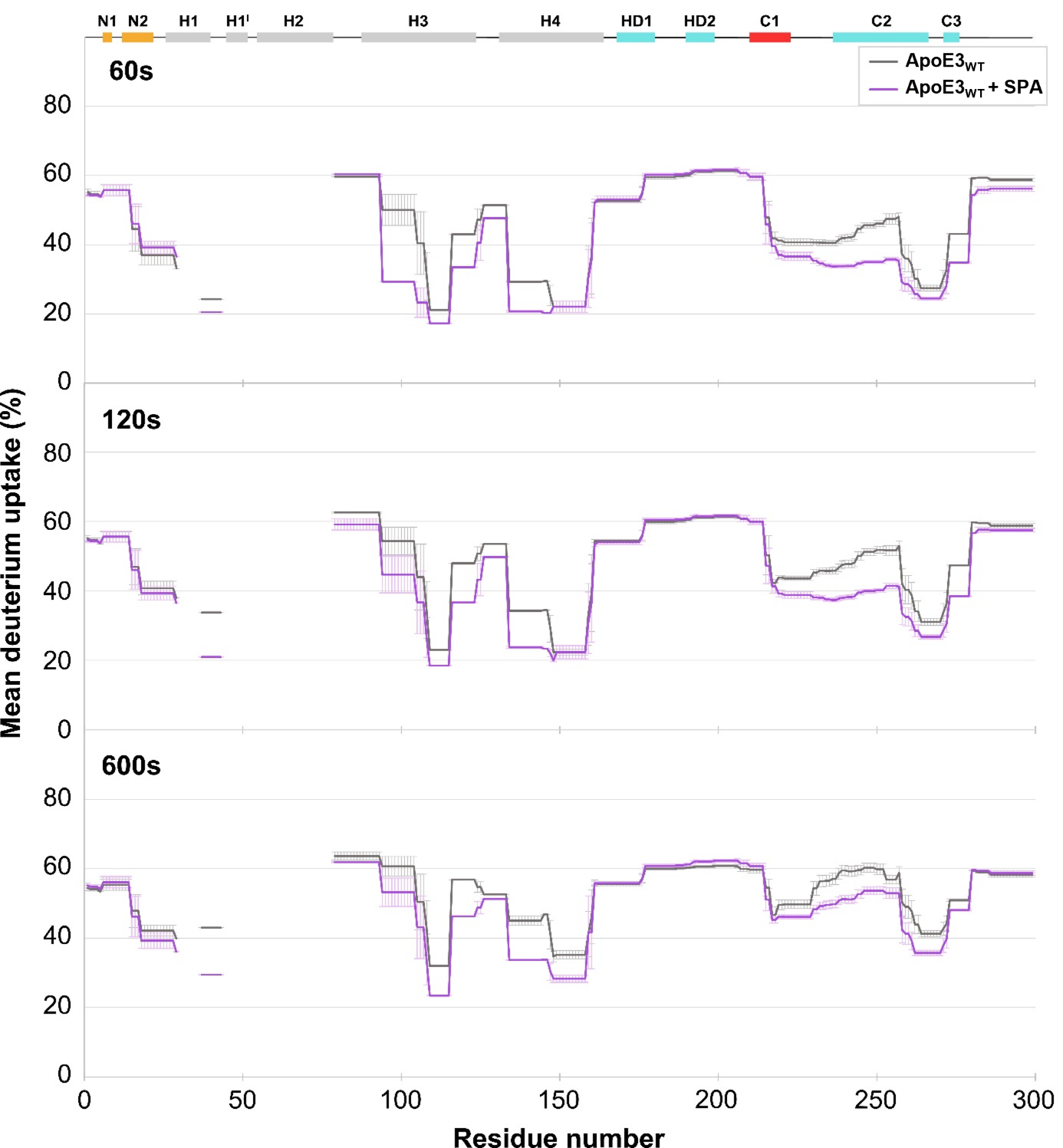

**Supplementary Figure 12**. Comparison of mean deuterium uptake of ApoE3_WT_ without SPA (yellow) and with SPA (purple) at 60, 120, and 600 second time points.

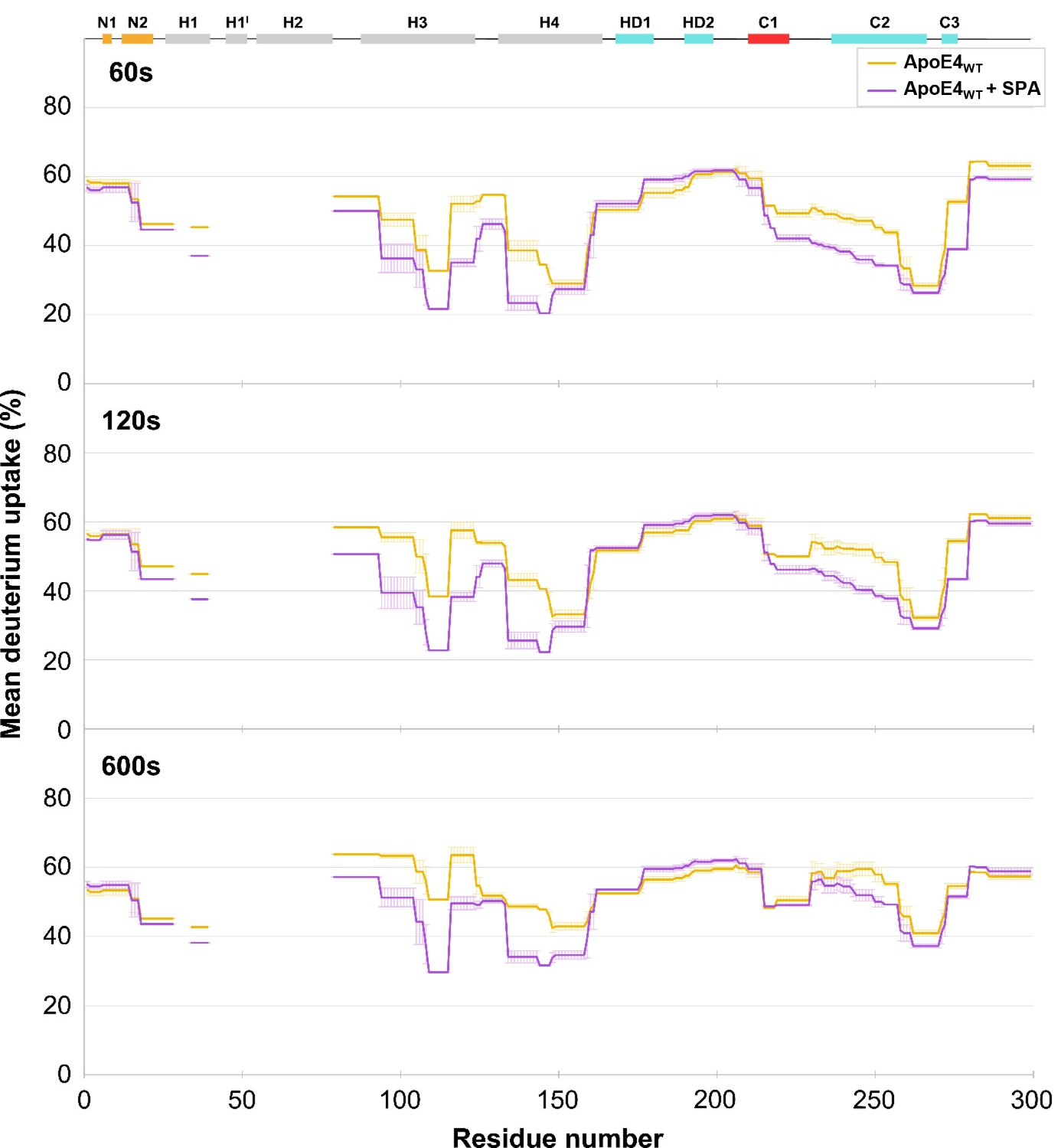

**Supplementary Figure 13.** Comparison of mean deuterium uptake of ApoE4_WT_ without SPA (yellow) and with SPA (purple) at 60, 120, and 600 second time points.

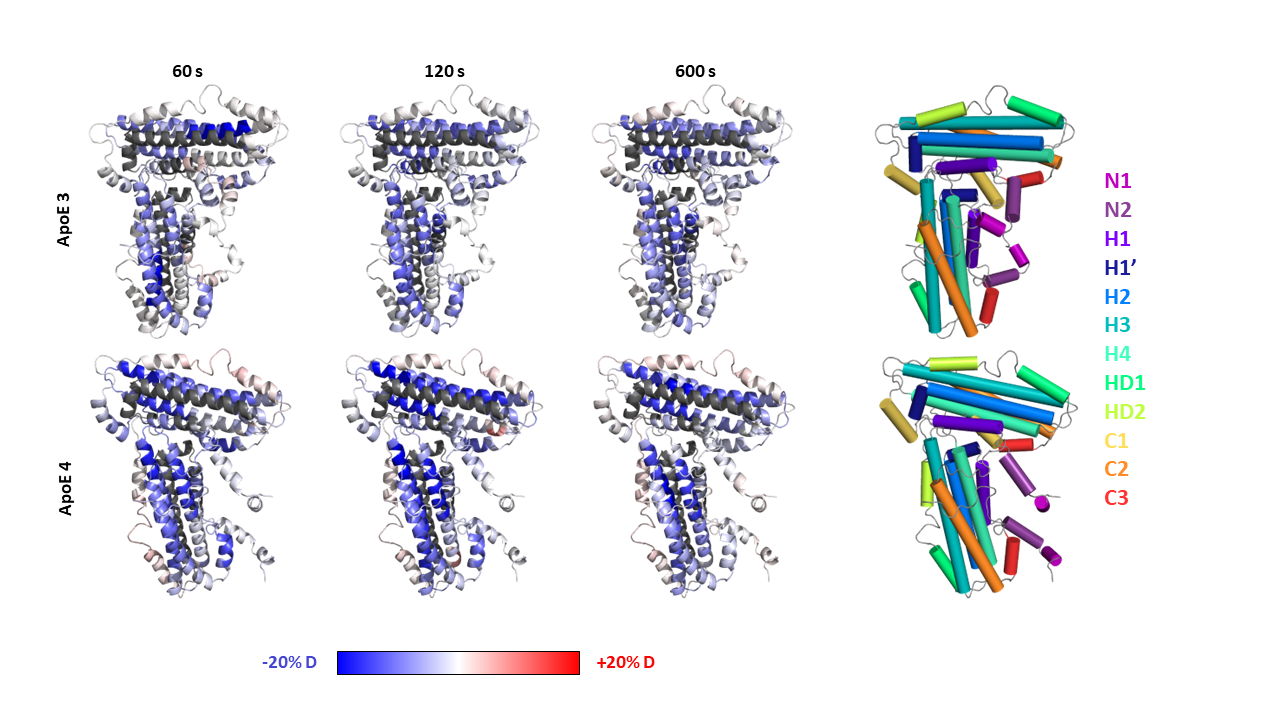

**Supplementary Figure 14.** SPA effect on ApoE_WT_ deuteration measured by HDX-MS. The difference in relative deuterium uptake as measured in the presence and absence of SPA is mapped to the constructed models (Methods) of wild-type ApoE3 (upper row) and ApoE4 (lower row), spanning ±20 relative units (legend at the bottom). Different times of the hydrogen/deuterium exchange experiment are shown in columns (60, 120 and 600 seconds). A color-coded reference to the secondary structural elements of ApoE is given on the right to facilitate the location of each structural element in the protein sequence. The main effect of SPA is shielding the protein structure (blue-shift), although this occurs to different extents in ApoE3 and ApoE4. Notably, the shielding effect in helix H3 is biased in ApoE3 (intense in the N-terminus, right; mild in the C-terminus, left), while it is more even and equally prominent in ApoE4. This difference relates to the differential unwinding observed in ApoE helix H3 (Figure 5E). The exposure resulting from SPA (red-shift) is milder than its shielding effect, and the regions that tend to get more exposed also differ in the two isoforms: N-terminal helix N1 tends to get more exposed in ApoE3 while the hinge domain helices HD1 and HD2 tend to be more exposed in ApoE4.

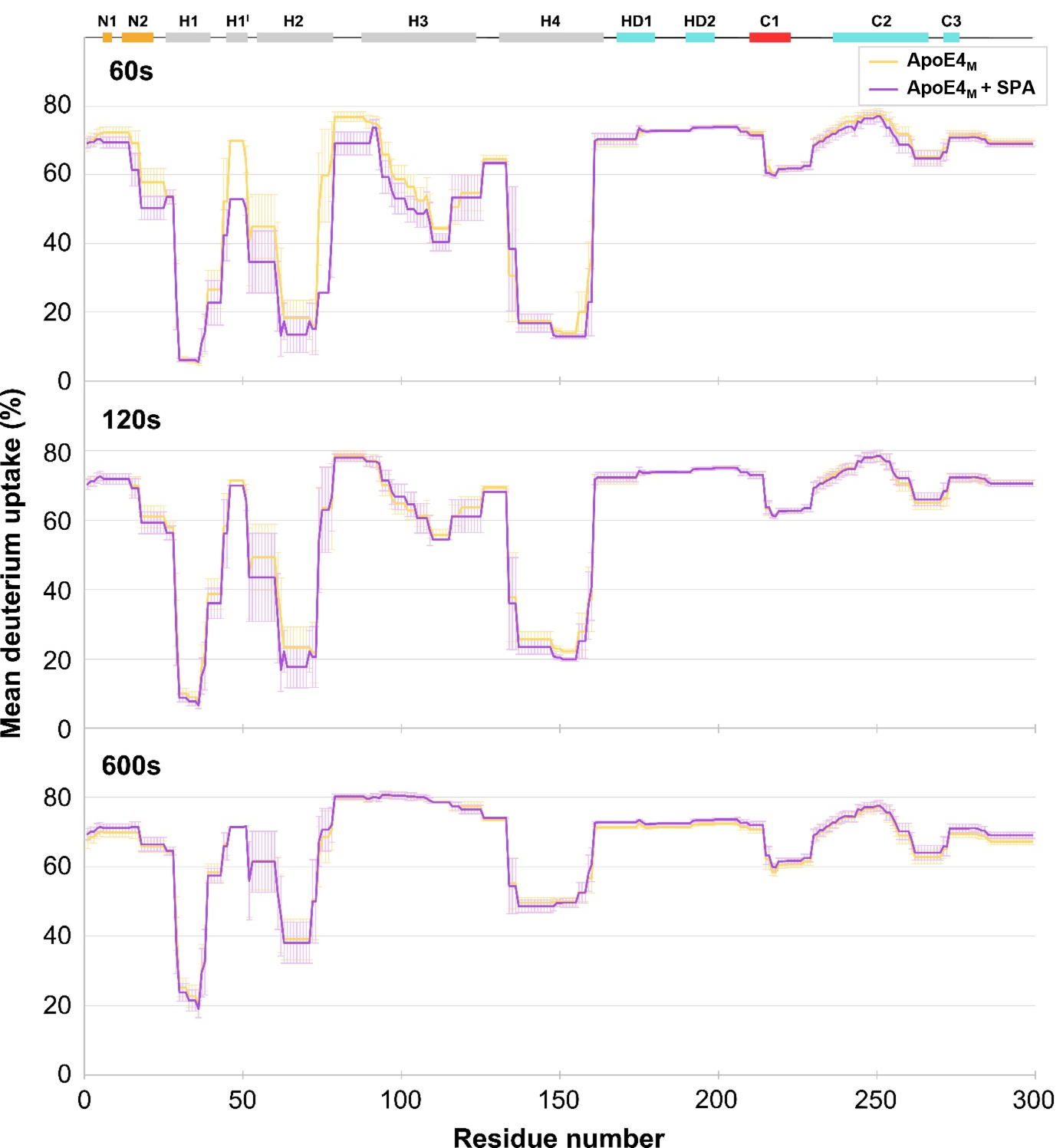

**Supplementary Figure 15.** Comparison of mean deuterium uptake of ApoE4_M_ without SPA (yellow) and with SPA (purple) at 60, 120, and 600 second time points.

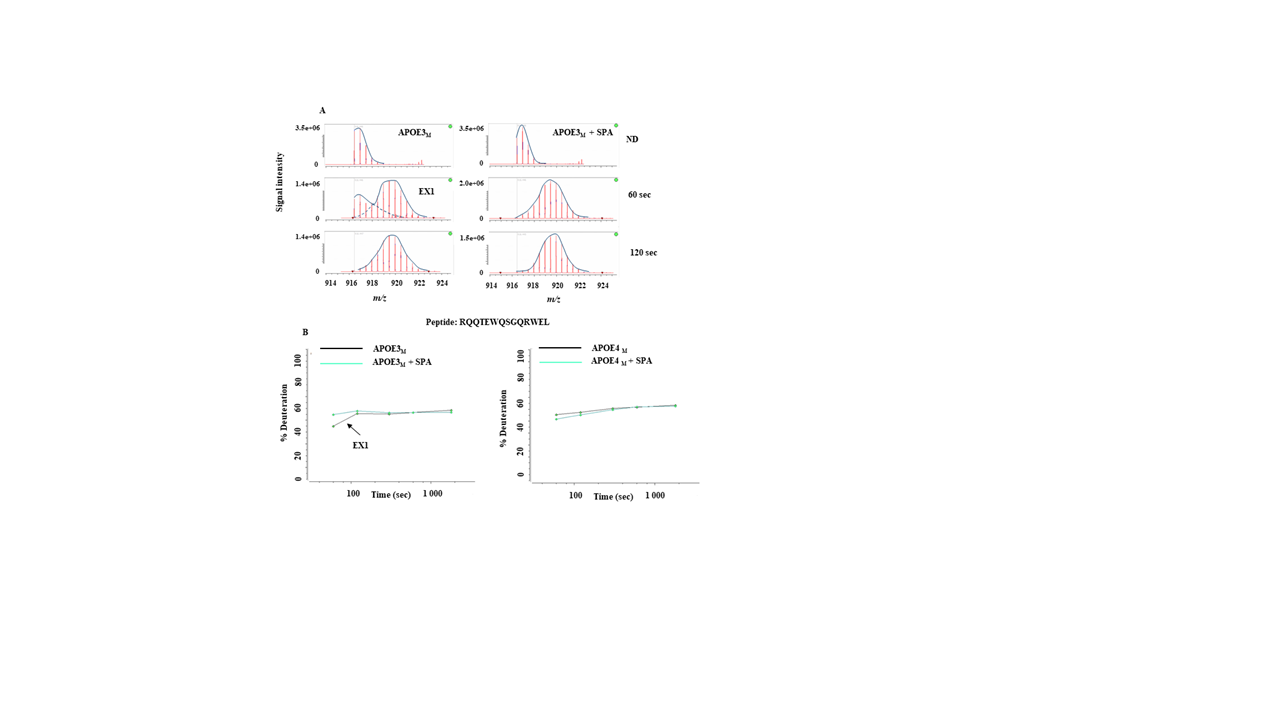

**Supplementary Figure 16.** HDX kinetics of ApoE3_M_ and ApoE4_M_ free or in the presence of 3-SPA. (A) MS spectra of ApoE3_M_ peptide free and in interaction with 3-SPA as non-deuterated (ND) and deuterated at 60 or 120 second time points. (B) Deuterium uptake of ApoE3_M_ and ApoE4_M_ (black line) and ApoE3_M_-SPA and ApoE4_M_-SPA (green line) at time course (60, 120, and 600 seconds).

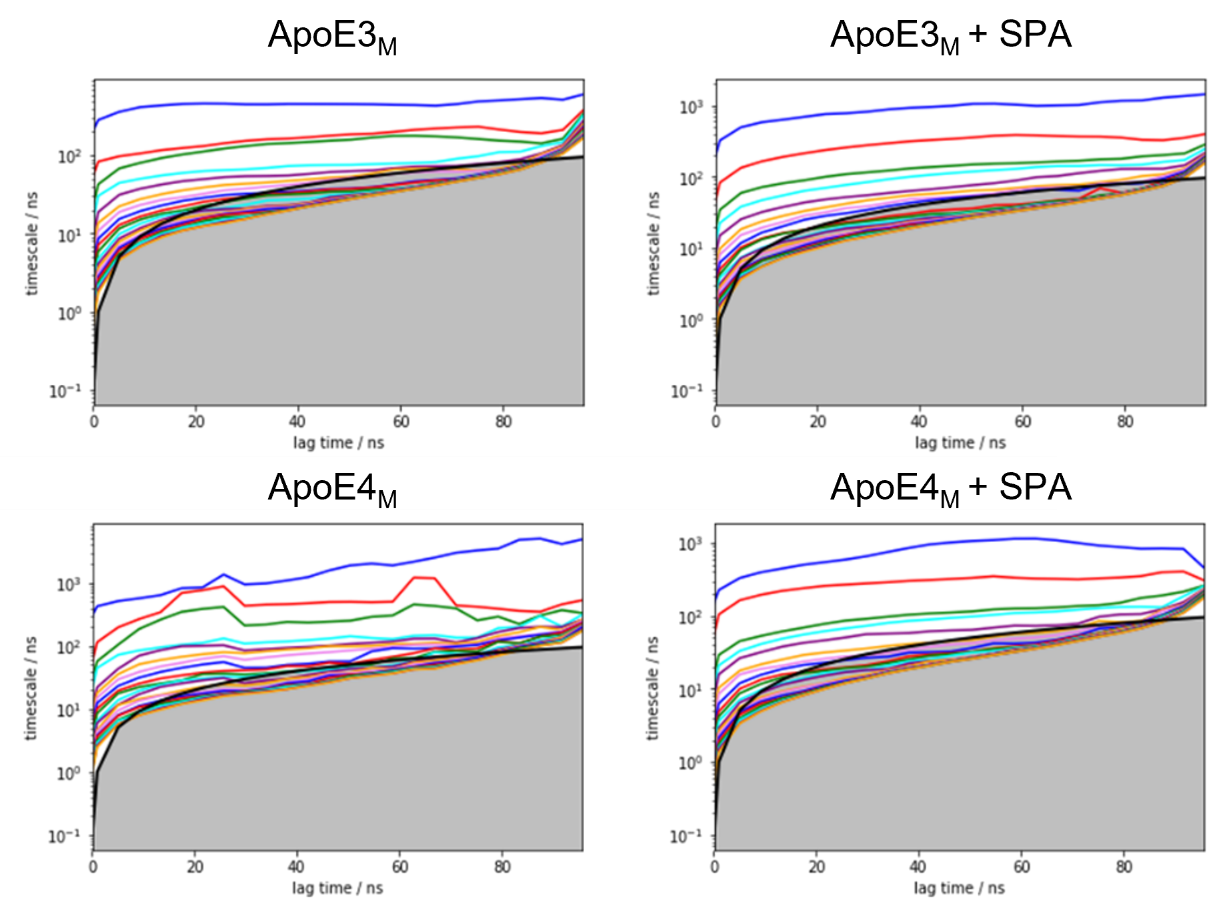

**Supplementary Figure 17.** Implied timescale plots for the MSMs of the simulations of ApoE3_M_ (top) and ApoE4_M_ (bottom), for the free systems (left) or with SPA (right).

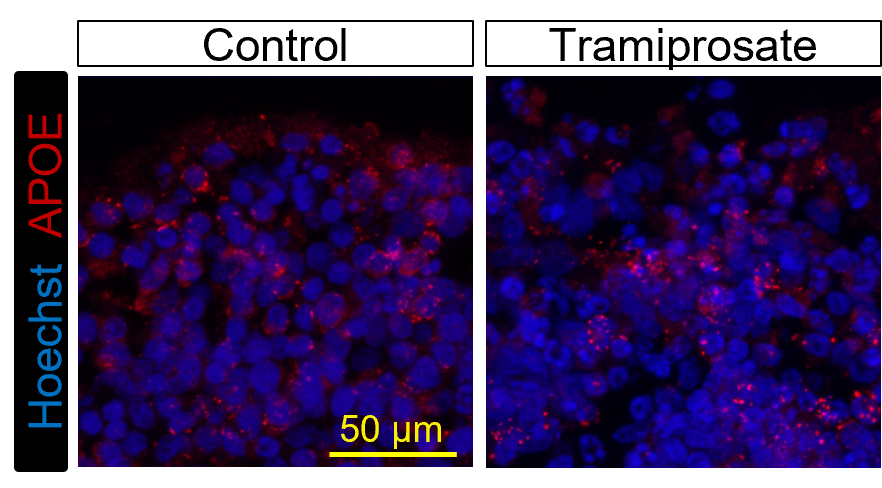

**Supplementary Figure 18**: Representative ApoE staining of control and tramiprosate-treated organoids.

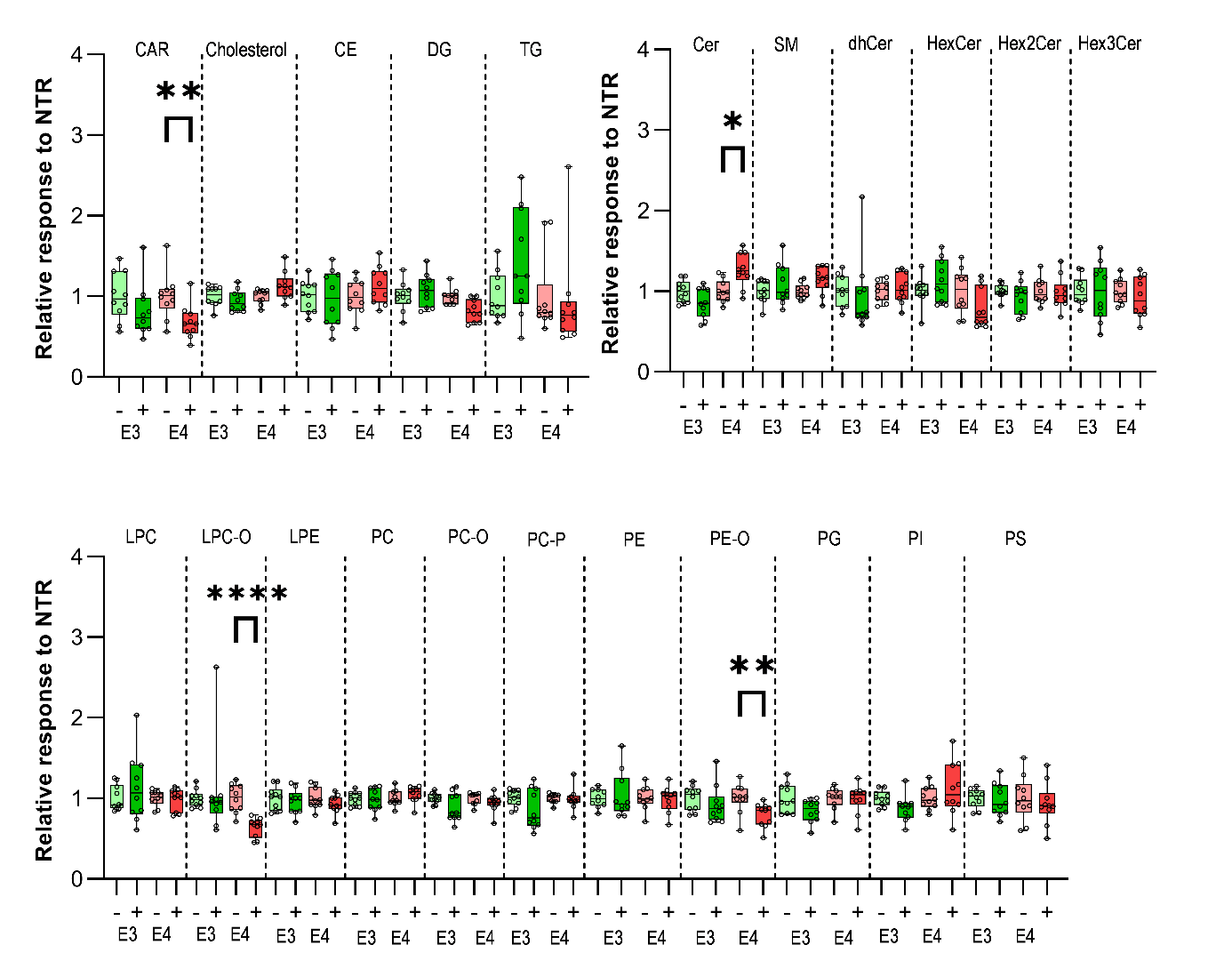

**Supplementary Figure 19**. Assessment of 22 total lipid classes upon tramiprosate treatment of ApoE ε3/ε3 (E3) and ApoE ε4/ε4 (E4) cerebral organoid. Statistical analysis was performed using the Kruskal-Wallis test, comparing the cerebral organoids treated with tramiprosate to the non-treated, *p≤0.05, **p≤0.01, ***p≤0.001, ****p≤0.0001.

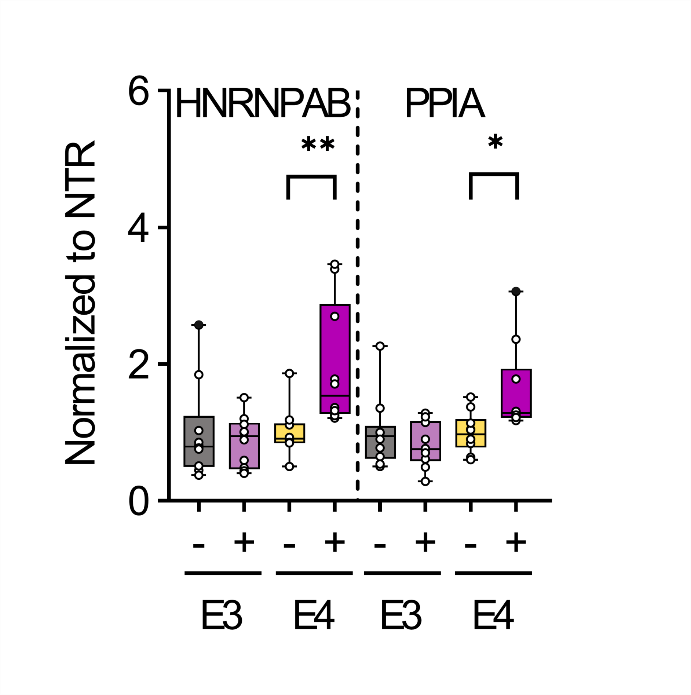

**Supplementary Figure 20**. HNRNPAB and PPIA protein levels in iPSC-derived ApoE ε3/ε3 (E3) and ApoE ε4/ε4 (E4) cerebral organoids non-treated control (-) and treated (+) with tramiprosate (n=10). HNRNPAB and PPIA were upregulated only in tramiprosate-treated E4 organoids relative to the levels in non-treated organoids. Outliers are presented as black dots (●). Statistics was performed using Kruskal-Wallis and uncorrected Dunn´s test (GraphPad), to compare non-treated and tramiprosate-treated cerebral organoids (n =10), *p≤0.05, **p≤0.001.

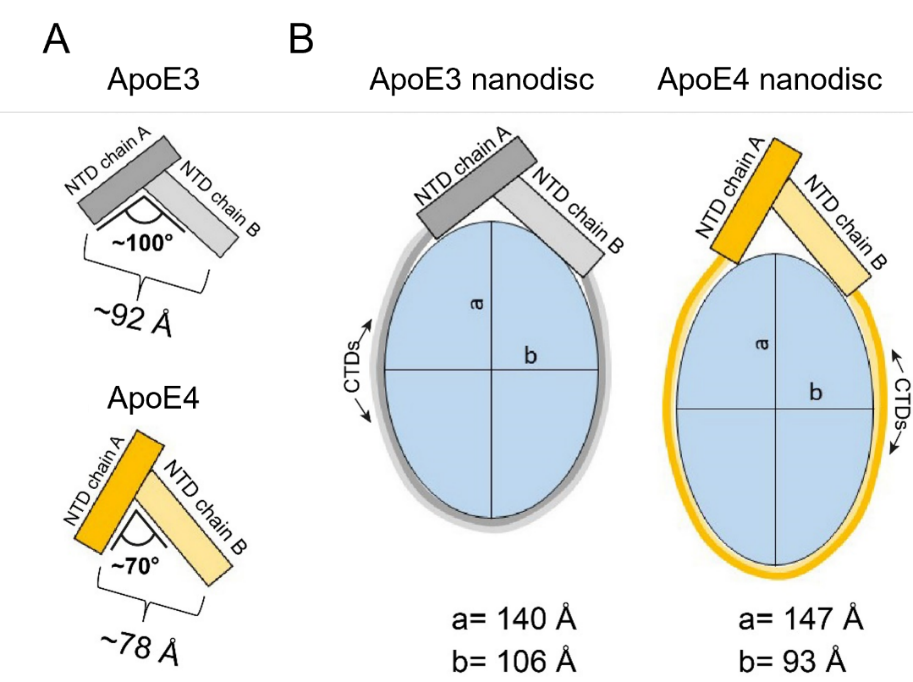

**Supplementary Figure 21**: Possible arrangement of ApoE nanodiscs based on ApoE T-shaped dimeric units. (A) Schematic representation of ApoE3 and ApoE4 T-shaped dimeric units. (B) Schematic representation of ApoE nanodiscs involving T-shaped dimeric units. The dimensions of the ApoE nanodiscs are taken from the work Waldie and co-authors [1].

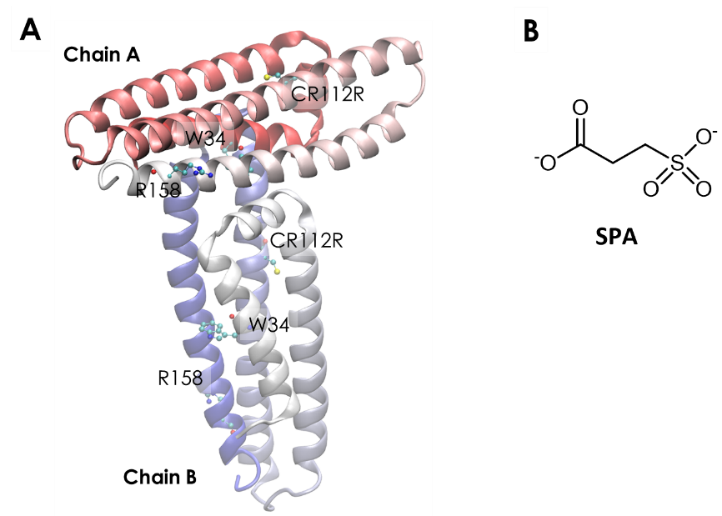

**Supplementary Figure 22.** A) Crystallographic structure of the T-shaped ApoE3 dimeric unit (PDB ID 1BZ4); B) chemical structure of 3-sulfopropanoic acid (SPA) in the dominant protonation state at the physiological pH (7.4). Chain A (horizontal) and chain B (vertical) are identified, as well as relevant residues which are shown as ball-and-sticks: the single-point mutation from ApoE3 to ApoE4, C112R, the mutation between ApoE2 and ApoE3/4, R158, and the dimer interface residue W34.

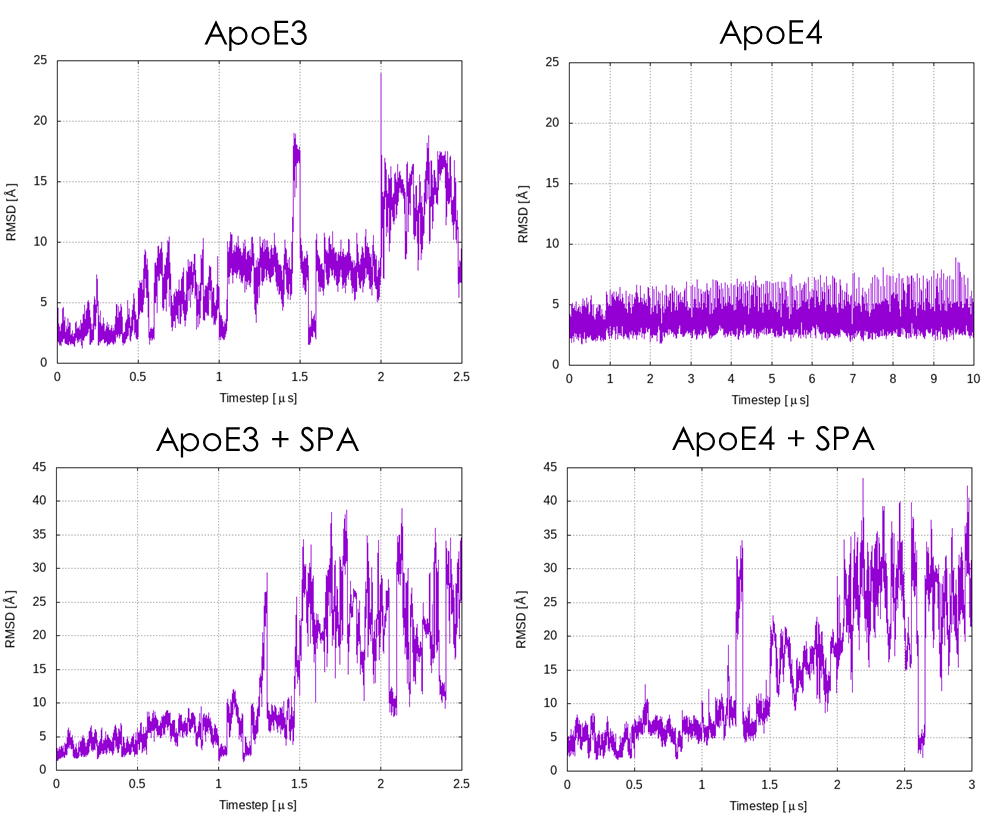

**Supplementary Figure 23.** Time variation of the RMSD of the backbone atoms of the T-shaped dimeric unit with respect to the respective initial structures, during the adaptive MDs. We can consider that the dimer is dissociated for RMSD values above ca. 15 Å.

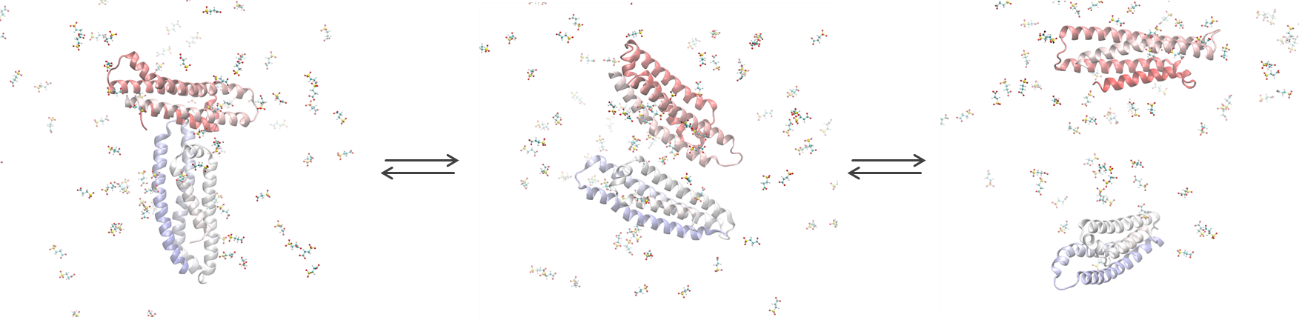

**Supplementary Figure 24.** Dissociation of the ApoE3 dimer in the presence of SPA. The initially well-associated T-shaped dimer (left) changed into the side-by-side interacting dimer (center; RMSD ca. 15-20 Å), and eventually led the distant monomeric units solvated by water and SPA (right; RMSD ca. 30-35 Å). These representative snapshots were obtained from the adaptive simulations of ApoE3 in the presence of SPA.

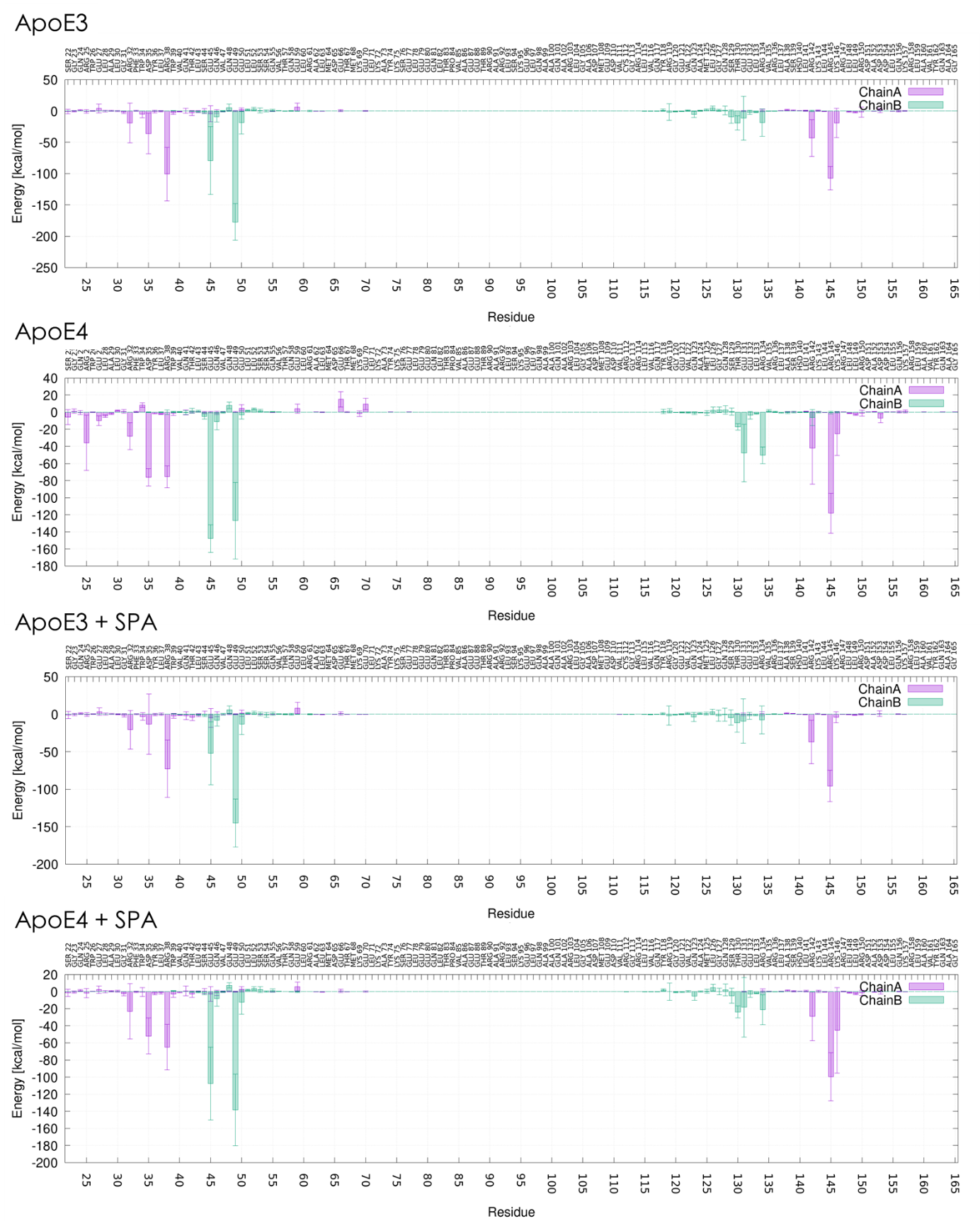

**Supplementary Figure 25.** Interactions of the different residues in each chain of the ApoE dimers with the opposite monomeric chain. Electrostatic component of the linear interaction energy (LIE), calculated for the first 500 ns of the concatenated adaptive MDs with the dimers of ApoE3 and ApoE4, without and with SPA. The error bars represent the standard deviations from the mean interaction energies.

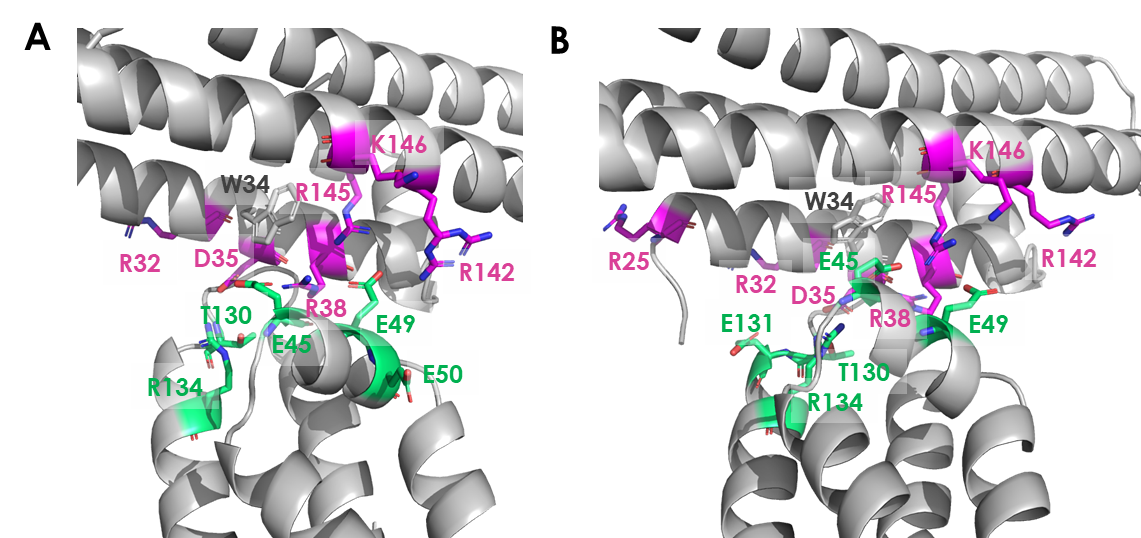

**Supplementary Figure 26.** Residues with the highest interaction energies at the dimer interface for A) ApoE3 and B) ApoE4. The residues in chain A are represented as magenta sticks, and in chain B as green sticks. W34 is also shown for reference.

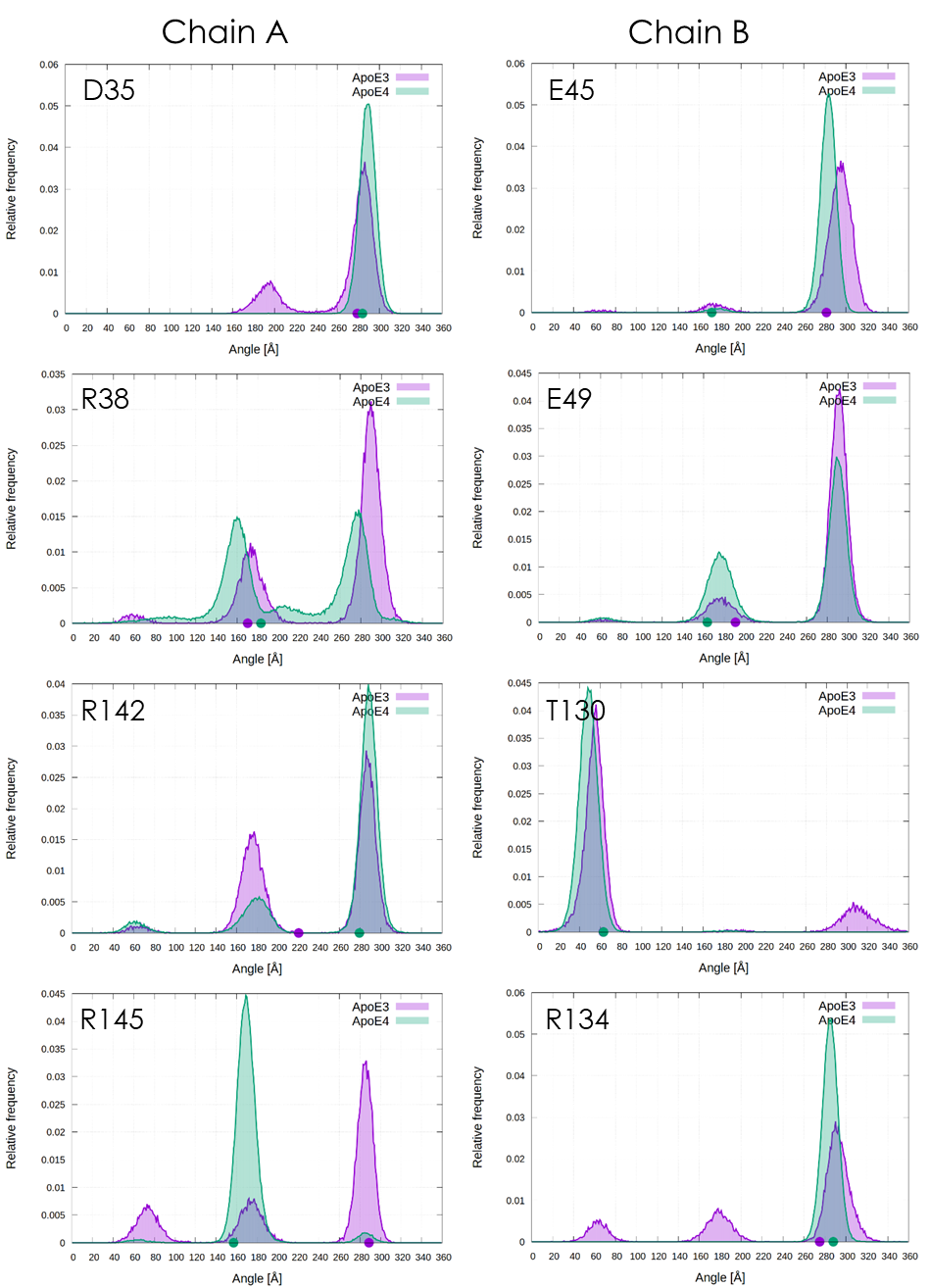

**Supplementary Figure 27**. Distribution of the side chain orientations of the main interface residues (χ1 dihedral angle) during the simulations of ApoE3 (purple) and ApoE4 (green). Left column: chain A residues D35, R38, R142, and R145; right column: chain B residues E45, E49, T130 and R134. The respective angles found in the crystal structures are represented by the circles with the same colours on the X-axis.

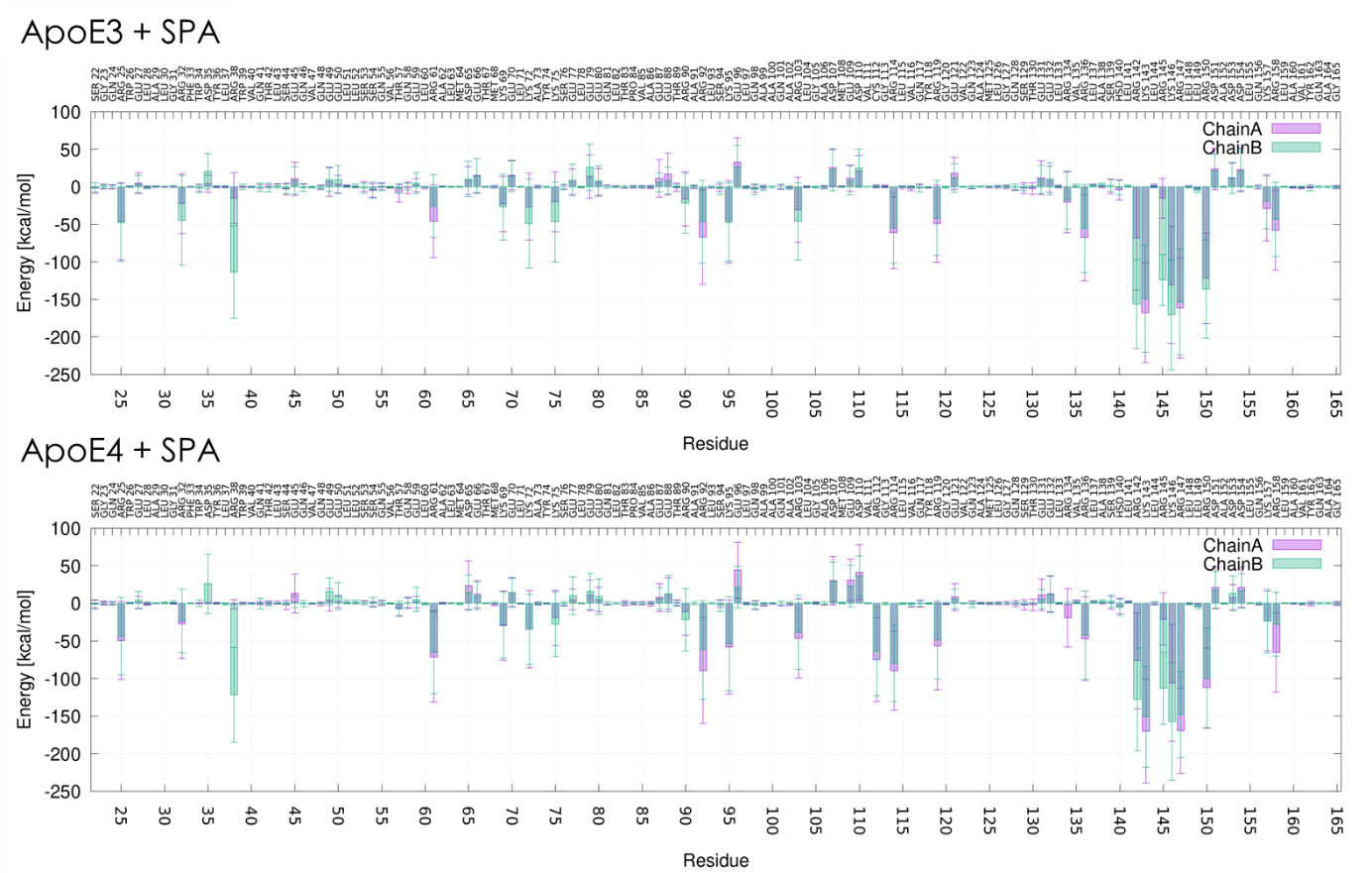

**Supplementary Figure 28.** Interactions of SPA with the dimers of ApoE3 and ApoE4. Electrostatic component of the linear interaction energy (LIE) of all the 100 molecules of SPA with each residue, calculated for the first 500 ns of the concatenated adaptive MDs. The error bars represent the standard deviations from the mean interaction energies.

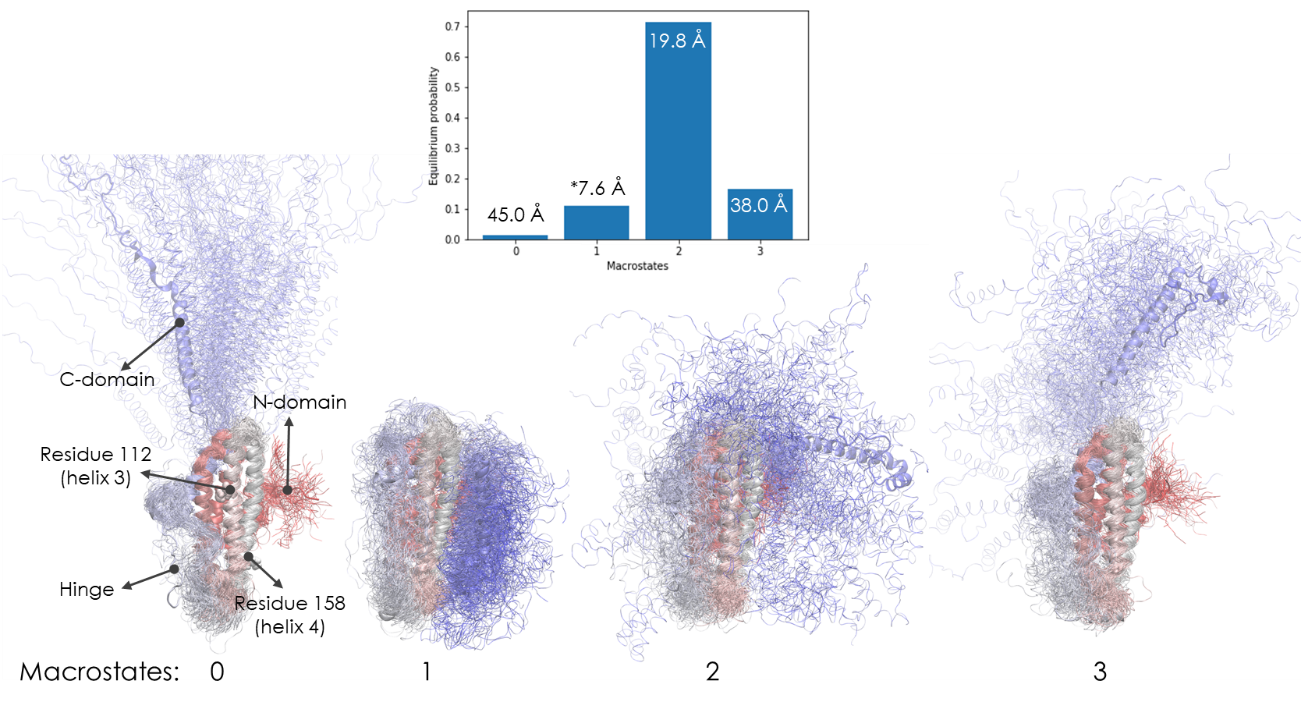

**Supplementary Figure 29.** Equilibrium distribution and illustrative structures of the macrostates obtained for ApoE3_M_. The respective average RMSD values are included by the labels; the symbol * identifies the most folded state. The cartoons are coloured by the sequence (red for N-terminus and blue for C-terminus); 100 random structures from each cluster are superimposed to illustrate the structural variability. Important regions are labelled for macrostate 0.

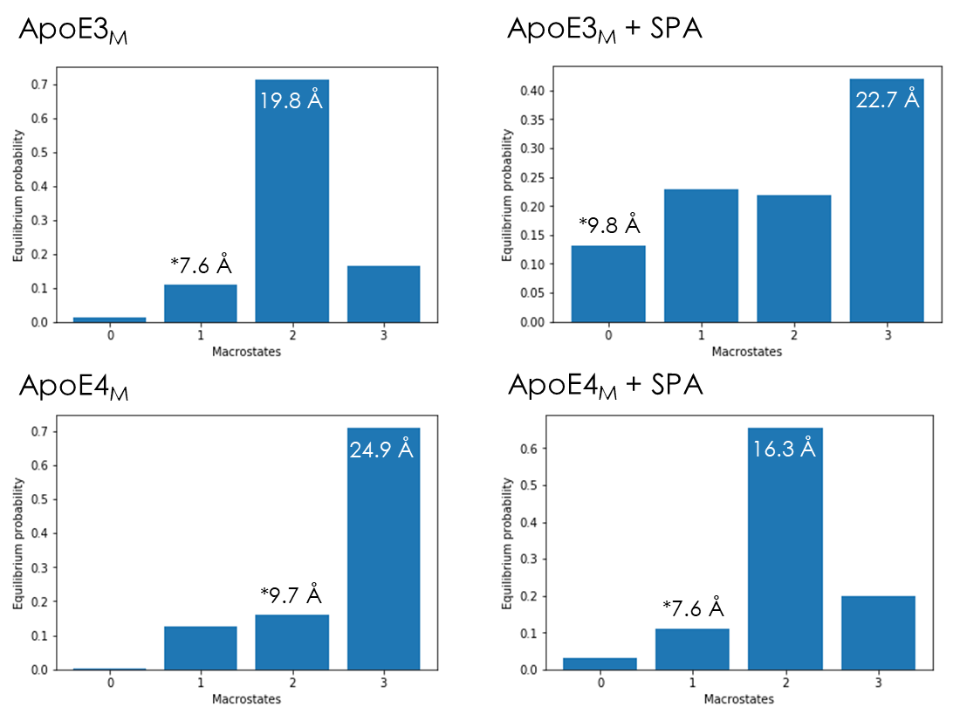

**Supplementary Figure 30.** Equilibrium populations of the MSMs for the simulations of ApoE3_M_ (top) and ApoE4_M_ (bottom), for the free systems (left) or with SPA (right). The labels indicate the average RMSD values of the closest folded state (marked with *) and the most populated state.

### SUPPLEMENTARY NOTES

#### Supplementary Note 1

**ApoE T-shaped dimeric unit dissociation**

To track the dissociation of the T-shaped ApoE dimeric units found in the crystal structures (**Supplementary Figure 22**), we performed molecular dynamics (MDs) simulations with the original crystal structures of those dimers. We note that we purposefully constructed the T-shaped and V-shaped dimers from the crystallographic structure to test our initial hypothesis that this dimeric form may have biological relevance. Herein we did not try other type of dimeric units that could be derived from the symmetry mates.

We used an adaptive sampling approach to survey the conformational diversity of the proteins, by specifying the root-mean-square deviation (RMSD) of the Cα atoms as the adaptive metric. By doing so, we encouraged the simulations to find different conformations of the protein backbone, and hence potentially accelerated the observation of dissociated states of the dimers. The simulations ran for a total combined time of 10 µs. We measured the RMSD of the backbone atoms of the protein during the adaptive simulations, after the concatenation and alignment of the sequential epochs of parallel MDs. We found that most of the studied systems *did* dissociate after 1.5-2 µs of combined simulation times. ApoE3 dissociated much faster (in ca. 2 µs) than ApoE4 (which *did not* dissociate at all during the 10 μs simulation). This can be assessed from the respective RMSD plots (**Supplementary Figure 23**).

In the presence of SPA, the ApoE dimers dissociated faster than without this molecule (**Supplementary Figure 22**). Moreover, the two monomeric units became involved by the SPA molecules, which disrupted the protein-protein interactions between the two chains and resulted in their drifting apart, often reaching much higher RMSD values (ca. 30-35 Å) than in the absence of SPA (ca. 15-20 Å). This is illustrated in **Supplementary Figure 24**.

**Inter-residue interactions in the ApoE T-shaped dimeric units**

To identify the residues contributing the most to holding the T-shaped dimeric units together, we computed the interactions of each residue with the opposite monomeric unit. For that, we also used the Linear Interaction Energy (LIE) method. The relevant interactions were dominated by the electrostatic component, with only a few exceptions (e.g., chain A: W39, R38, Y36; chain B: L126, G127, Q123; data not shown). The residues interacting with the opposite chains were different for the two chains (**Supplementary Figure 25**). This was expected, considering that the self-association interface is not symmetric for those two chains (**Supplementary Figure 22**).

Many residues also interacted differently in the two ApoE variants, which was a consequence of the respective dimers presenting different contacting interfaces (**Supplementary Figure 25-26**, and **Supplementary Table 8**). Interestingly, the interface residues in chain A are mostly positively charged, while in chain B they are mainly negatively charged.

Considering the total average LIE energies between the two chains (**Supplementary Table 8**), we found that the electrostatic interactions between the monomeric units in the dimer of ApoE4 were stronger and fluctuated less (*E*_elec_ = -396 ± 68 kcal/mol) than for ApoE3 (*E*_elec_ = -338 ± 84 kcal/mol). This difference is extremely significant (p-value < 10^-4^ from the *t*-test), and it may explain why the ApoE4 T-shaped dimeric unit did not dissociate during the simulation, in contrast to ApoE3. On the other hand, the dimer interactions were significantly reduced in the presence of SPA in both ApoE variants, compared to the absence of SPA, thus explaining their increased propensity to dissociate.

To investigate to which extent the main interface residues preserved their crystallographic positions during the simulations, we computed the χ1 dihedral angles (defined by atoms N-Cα–Cβ-Cγ) as a measure of the orientation of their side chains and plotted their population distributions. This analysis was performed for the residues: D35, R38, R142, and R145 in chain A; and E45, E49, T130 and R134 in chain B (residues with the strongest interactions, see Supplementary Table 8). In general, the results (Supplementary Figure 27) showed that: 1) the orientation of those residues was more dispersed (more scattered population peaks) in ApoE3 than in ApoE4; 2) ApoE4 preserved more closely the conformations found in the crystal structure than ApoE3. This finding is consistent with the dimer being more rigid and tightly bound for ApoE4 than ApoE3. There are only a few exceptions to these general observations among the reported residues, namely for R38 in chain A (it can adopt two main orientations in ApoE3 and ApoE4) and E45 in chain B (its dominant conformation in ApoE4 is different from the crystallographic one).

#### Supplementary Note 2

**Interactions of ApoE T-shaped dimeric units with SPA**

The interactions of SPA with the residues of the ApoE T-shaped dimeric units were assessed by the linear interaction energy (LIE), computed for all the 100 ligand molecules with each residue over the combined MDss (**Supplementary Figure 28**). The electrostatic component dominated the interactions due to the double negative charge of SPA at pH 7.4 (**Supplementary Figure 22**). Moreover, these interactions were much stronger with the charged residues compared to the neutral ones, in some cases attractive (ΔG < 0, with the positive residues) in other cases repulsive (ΔG > 0, with negative residues). Some of the largest differences between the interactions of SPA with the dimers of ApoE3 and ApoE4 were found for the mutated residue C112R, R61, and R134 and R158 (**Supplementary Figure 28**).

Some differences were observed between the SPA interactions with the same residues in chain A and chain B, in both proteins. These differences were mainly found in the residues located at the dimer interface, which were more shielded from the SPA molecules. Some of these residues, however, still interacted with SPA, which very likely contributed to decrease the protein-protein interactions formed by these residues.

#### Supplementary Note 3

**Markov state analysis and unfolding of the ApoE_M_**

The full-length monomers of ApoE3_M_ and ApoE4_M_ were simulated by adaptive MDs. As done for the dimers, also here we encouraged the sampling of conformational diversity by specifying the adaptive metric as the RMSD of the Cα atoms of the proteins. The simulations ran for a total combined time of 20 µs, and we constructed Markov state models (MSMs) for these simulations, using the RMSD of the protein C_α_ atoms. We obtained reasonably converged implied time scales (**Supplementary Figure 17**), which suggested 3-4 distinct macrostates. For this reason, we required 4 states in the analysis of all the systems.

The distribution of the RMSD values was extremely varied, mostly due to a high fluctuation of the C-domain, while the N-domain was very stable. For ApoE3_M_, the most populated state had an average RMSD of 19.8 Å, and the closest state to the initial folding had an RMSD of 7.6 Å, with only a 10.8% equilibrium population. The most extensively “unfolded” state (the most extended conformation), although little populated, showed an average RMSD of 45 Å and the C-domain stretching away from the N-domain (**Supplementary Figure 29** and **Supplementary Table 11**). The other systems showed similar trends (**Supplementary Figure 29**), with high “unfolding” rates and equilibrium that favours the unfolding compared to the folding process (**Supplementary Table 11**). Interestingly, ApoE4_M_ showed a higher propensity towards the folded state than ApoE3_M_, but in the presence of SPA, this trend was partially reverted. We note that we arbitrarily call the process described here “unfolding”, simply due to comparing the conformational states with the initial more globular “folded” state. Our results are in agreement with a recent study, which describes the ApoE4_M_ as an equilibrium of between the “closed”, “open” and “extended” conformations [2]. These conformations are similar to our macrostates between “folded” and “unfolded” conformations (**Supplementary Figure 29**).

#### Supplementary Note 4

The proposed arrangement of ApoE nanodiscs involving ApoE T-shaped dimeric units is consistent with the previous model [3]. In that model two ApoE molecules (or one dimer interact together via the NTDs and surround the lipid particle with CTDs. Hypothetically, if the ApoE T-shaped dimeric unit represented such a dimer it would explain why ApoE4 is less lipidated than ApoE3 [4], since only a monomeric ApoE binds lipids [5]. The angle difference between ApoE3 and ApoE4 T-shaped dimeric units could also rationalize the experimental observation that ApoE4 lipid particles are more ellipsoidal compared to the ApoE3 particles. Interestingly, the difference in the distances of the two ends of the ApoE3 and ApoE4 NTDs (92 - 78Å = 14 Å) corresponds to the difference in the short semi-axes of the nanodisc diameters (106 - 93Å = 13Å).

### SUPPLEMENTARY REFERENCES

1. Waldie S, Sebastiani F, Moulin M, Del Giudice R, Paracini N, Roosen-Runge F, et al. ApoE and ApoE Nascent-Like HDL Particles at Model Cellular Membranes: Effect of Protein Isoform and Membrane Composition. Frontiers in Chemistry [Internet]. 2021 [cited 2022 Jun 3];9. Available from: https://www.frontiersin.org/article/10.3389/fchem.2021.630152

2. Stuchell-Brereton MD, Zimmerman MI, Miller JJ, Mallimadugula UL, Incicco JJ, Roy D, et al. Apolipoprotein E4 has extensive conformational heterogeneity in lipid free and bound forms [Internet]. bioRxiv; 2022 [cited 2023 Feb 22]. p. 2022.02.02.478828. Available from: https://www.biorxiv.org/content/10.1101/2022.02.02.478828v1

3. Henry N, Krammer E-M, Stengel F, Adams Q, Liefferinge FV, Hubin E, et al. Lipidated apolipoprotein E4 structure and its receptor binding mechanism determined by a combined cross-linking coupled to mass spectrometry and molecular dynamics approach. PLOS Computational Biology. Public Library of Science; 2018;14:e1006165.

4. Hu J, Liu C-C, Chen X-F, Zhang Y, Xu H, Bu G. Opposing effects of viral mediated brain expression of apolipoprotein E2 (apoE2) and apoE4 on apoE lipidation and Aβ metabolism in apoE4-targeted replacement mice. Molecular Neurodegeneration. 2015;10:6.

5. Garai K, Baban B, Frieden C. Dissociation of Apolipoprotein E Oligomers to Monomer Is Required for High-Affinity Binding to Phospholipid Vesicles. Biochemistry. American Chemical Society; 2011;50:2550–8.
